# Rational Design of Enhanced Nme2Cas9 and Nme2^Smu^Cas9 Nucleases and Base Editors

**DOI:** 10.1101/2024.10.30.620986

**Authors:** Nathan Bamidele, Aditya Ansodaria, Zexiang Chen, Haoyang Cheng, Rebecca Panwala, Eva Jazbec, Erik J. Sontheimer

## Abstract

CRISPR-Cas genome editing tools enable precise, RNA-guided modification of genomes within living cells. The most clinically advanced genome editors are Cas9 nucleases, but many nuclease technologies provide only limited control over genome editing outcomes. Adenine base editors (ABEs) and cytosine base editors (CBEs) enable precise and efficient nucleotide conversions of A:T-to-G:C and C:G-to-T:A base pairs, respectively. Therapeutic use of base editors (BEs) provides an avenue to correct approximately 30% of human pathogenic variants. Nonetheless, factors such as protospacer adjacent motif (PAM) availability, accuracy, product purity, and delivery limit the full therapeutic potential of BEs. We previously developed Nme2Cas9 and its BE derivatives, including ABEs compatible with single adeno-associated virus (AAV) vector delivery, in part to enable editing near N_4_CC PAMs. Further engineering yielded domain-inlaid BEs with enhanced activity, as well as Nme2Cas9/SmuCas9 chimeras that target single-cytidine (N_4_C) PAMs. Here we further enhance Nme2Cas9 and Nme2^Smu^Cas9 editing effectors for improved efficiency and vector compatibility through site-directed mutagenesis and deaminase linker optimization. Finally, we define the editing and specificity profiles of the resulting variants by using paired guide-target libraries.

## Introduction

By enabling the alteration and correction of genes that contribute to disease, CRISPR-Cas genome editing tools are quickly advancing in the field of genetic medicine ^1,2^. Emerging tools have provided unparalleled control over the types of genetic manipulations that are possible ^3^. Base editors (BEs), composed of a nickase nCas9 fused to a nucleotide deaminase, are precision genome editors that enable single-nucleotide conversions ^4–6^. The most commonly used classes of BEs include adenine BEs (ABEs) and cytidine BEs (CBEs) that enable A:T-to-G:C or C:G-to-T:A base pair conversions, respectively ^3^. In principle, these two classes of BEs can together correct a large majority of disease-causing single nucleotide polymorphisms ^3^.

Despite the promise of genome editors to correct diseases at their root causes, in many cases *in vivo* delivery represents a major hurdle for the application of BEs and other editing systems ^7,8,8^. Adeno-associated virus (AAV) vectors are valuable delivery vehicles for genetic cargo to extra-hepatic tissues but are constrained by packaging capacity, among other limitations. Commonly used Cas9 variants such as SpyCas9 are large enough to greatly complicate AAV delivery, especially when used in combination with deaminases ^4–6,8^ or other fusion domains. The use of compact Cas9s such as Nme2Cas9 for nuclease ^9^ and BE ^10,11^ systems enhance the utility of gene editing tools from a variety of standpoints, including: (1) compact size, streamlining *in vivo* delivery through the use of single AAV vectors; (2) a cytidine dinucleotide PAM requirement, enabling access to pyrimidine-rich genome targets, and (3) editing specificity. Despite these benefits, initial Nme2Cas9-derived ABE systems exhibited modest editing efficiencies ^10,11^. Although the N_4_CC dinucleotide PAM provided additional flexibility when compared to other natural compact Cas9 variants ^12–15^, the PAM still restricts accessible BE editing windows.

Our initial efforts to improve the activity and targeting scope of Nme2Cas9 editing systems focused on BE applications ^16^. These efforts included internal rather than terminal deaminase domain insertion, which improved editing efficiencies while providing additional control over editing windows. We also developed Nme2^Smu^Cas9 ABE systems in which the single-cytidine PAM-interacting domain (PID) of SmuCas9 ^17^ was transplanted into Nme2Cas9 to enable targeting of N_4_C PAMs ^16^. Independently, phage evolution also yielded enhanced Nme2 variants (the eNme2-C ABE and the eNme2-C.NR nuclease) with N_4_C PAM compatibility ^18^. Despite these advances, when targeting sites with an N_4_CC PAM, Nme2^Smu^-ABEs suffered from diminished activity when compared to their WT PID counterparts. Other groups developing their own Nme2^Smu^Cas9 systems for nuclease and BE applications have reported similar findings ^19,20^. Interestingly, this decrease in activity at target sites with canonical PAMs is a common trend observed among PAM-relaxed Cas9 variants ^21–25^. Although our use of the transplanted PID from SmuCas9 increased PAM targetability, for ABE applications it came with a modest increase in size, potentially affecting single-vector AAV delivery with our format. Here we have further optimized domain-inlaid Nme2^Smu^Cas9 ABE systems to improve activity and minimize size for AAV deliverability. We also show that our editing activity improvements extend to Nme2^Smu^Cas9 nuclease as well as Nme2Cas9 systems with native PAM specificities.

## Results

### Rational design of Nme2^Smu^Cas9 nuclease and base editing mutants with increased activity

Nme2^Smu^Cas9’s subpar activities at N_4_CC targets sites prompted us to improve its activities through rational engineering. First, to recapitulate these findings, we compared Nme2Cas9 and Nme2^Smu^Cas9 editors in either their nuclease format or the domain-inlaid ABE8e-i1 architecture using a small subset of N_4_CC target sites (**Supplementary Fig. 1a, b**). Following plasmid transfection in HEK293T cells across three N_4_CC target sites, we observed a 2.2-and 2-fold decrease in editing efficiencies for the nuclease and ABE8e-i1 Nme2^Smu^Cas9 editors vs. their WT PID counterparts, respectively.

The enhancement of non-specific interactions between Cas effectors and the target nucleic acid is a common strategy employed to improve the activities of various genome editing reagents ^21,24,26,27^. We hypothesized that introducing arginine residues in proximity to either the DNA heteroduplex [target strand (TS) or non-target strand (NTS)] or the sgRNA could improve Nme2^Smu^Cas9 editing activity at N_4_CC and potentially N_4_CD (D = not C) target sites. To test this hypothesis, we selected candidate Nme2^Smu^Cas9 residues within 5-10 angstroms of guides or substrates using our previously generated SWISS homology model (**Supplementary Fig. 1c; Supplementary Table 1**). In addition to the Arg mutants, we also included a T72Y substitution analogous to the L58Y substitution of enCjeCas9 and reported to enhance base stacking interactions with the sgRNA ^27^. T72Y is located in the bridge helix and situated within the core of the Nme2Cas9/sgRNA/DNA ternary structure ^28^, likely forming a base-stacking interaction with U64 of the sgRNA. We incorporated a total of 33 single mutations into plasmids encoding the domain-inlaid Nme2^Smu^-ABE8e-i1 effector, and transfected them along with sgRNA-expressing plasmids into an established HEK293T ABE mCherry reporter cell line ^11,29^.

Among the TS-DNA interacting mutants, the E520R and D873R mutations substantially outperformed Nme2^Smu^-ABE8e-i1 (WT), increasing activation of the fluorescent reporter (**Supplementary Fig. 2a**). In addition, the panel of sgRNA and NTS-DNA proximity mutants produced eight variants with higher editing activities (**Supplementary Fig. 2a**). Mutations of note include the E932R, D56R and D1048R substitutions. Following our initial experiment, we re-evaluated the top two performing mutants from the TS-DNA panel and top eight variants from the sgRNA and NTS-DNA panel at additional N_4_CD PAM target sites within the ABE mCherry reporter (**Supplementary Fig. 2b**). In agreement with our pilot experiments, all mutations outperformed wildtype Nme2^Smu^-ABE8e-i1 (WT).

Encouraged by these results, we tested whether these mutations would increase the nuclease activity of Nme2^Smu^Cas9. We cloned the E520R and D873R substitutions, in addition to the entire panel of sgRNA and NTS-DNA mutants, into plasmids encoding Nme2^Smu^Cas9 nuclease. Subsequently, we transfected the panel of mutants along with sgRNA-expressing plasmids targeting the integrated traffic light reporter TLR-MCV1 in HEK293T cells ^30^ to report on DSB activity. For this assay, we screened editing activities at four target sites, varying the nucleotide in the 6th PAM position to cover all N_4_CN PAMs targetable by the SmuCas9 PID. As an additional benchmark of activity, we included Nme2Cas9 nuclease with the WT PID. As expected, Nme2Cas9 outperformed Nme2^Smu^Cas9 at the N_4_CC PAM target site (**Supplementary Fig. 2c**).

Encouragingly, the top-performing variants identified in our initial ABE8e screens also enhanced nuclease Nme2^Smu^Cas9 performance across all tested target sites, displaying a positive correlation in activity increase compared to the WT effector (Pearson *r* = 0.898) (**Fig. 1a**). Subsequently, we combined the top five mutations identified from our preliminary screens: E932R, D873R, D56R, E520R and D1048R. Intriguingly, 4 out of the 5 mutants spatially clustered around the PAM DNA duplex, while E520R was located closer to the PAM-distal end of the TS-DNA (**Fig. 1b, 1c**). We then screened the Nme2^Smu^Cas9 combinatorial variants in nuclease format. We anticipated that the enhancements that improved DSB activity would be most likely to translate to our domain-inlaid BE architectures. We repeated the TLR-MCV experiments for nuclease activity with 31 out of the 32 possible Nme2^Smu^Cas9 permutations including WT (**Supplementary Fig. 3**). As additional activity benchmarks, we included Nme2Cas9 (WT) and the nuclease eNme2-C.NR. To enable controlled comparisons, eNme2-C.NR was either assessed with the originally described NLS-linker architecture (eNme2-C.NR vLiu)^18^ or with an NLS-linker architecture identical to that of our Nme2^Smu^Cas9 plasmids (eNme2-C.NR vEJS). As previously demonstrated, Nme2Cas9 performed best at the N_4_CC target site, with background levels of activity at the N_4_CD target sites (**Supplementary Fig. 3**). In contrast, eNme2-C.NR and Nme2^Smu^Cas9 derivatives performed well at all N_4_CN PAM target sites. Notably, at N_4_CD PAM target sites, we observed similar TLR-MCV activation when comparing Nme2^Smu^Cas9 (WT) and eNme2-C.NR (**Supplementary Fig. 3**). Lastly, 23 out of the 30 Nme2^Smu^Cas9 combination mutants performed comparably to or better than their WT counterpart. The top performers in the first quartile consisted of single and double mutants (**Supplementary Fig. 3**).

**Figure 1.**
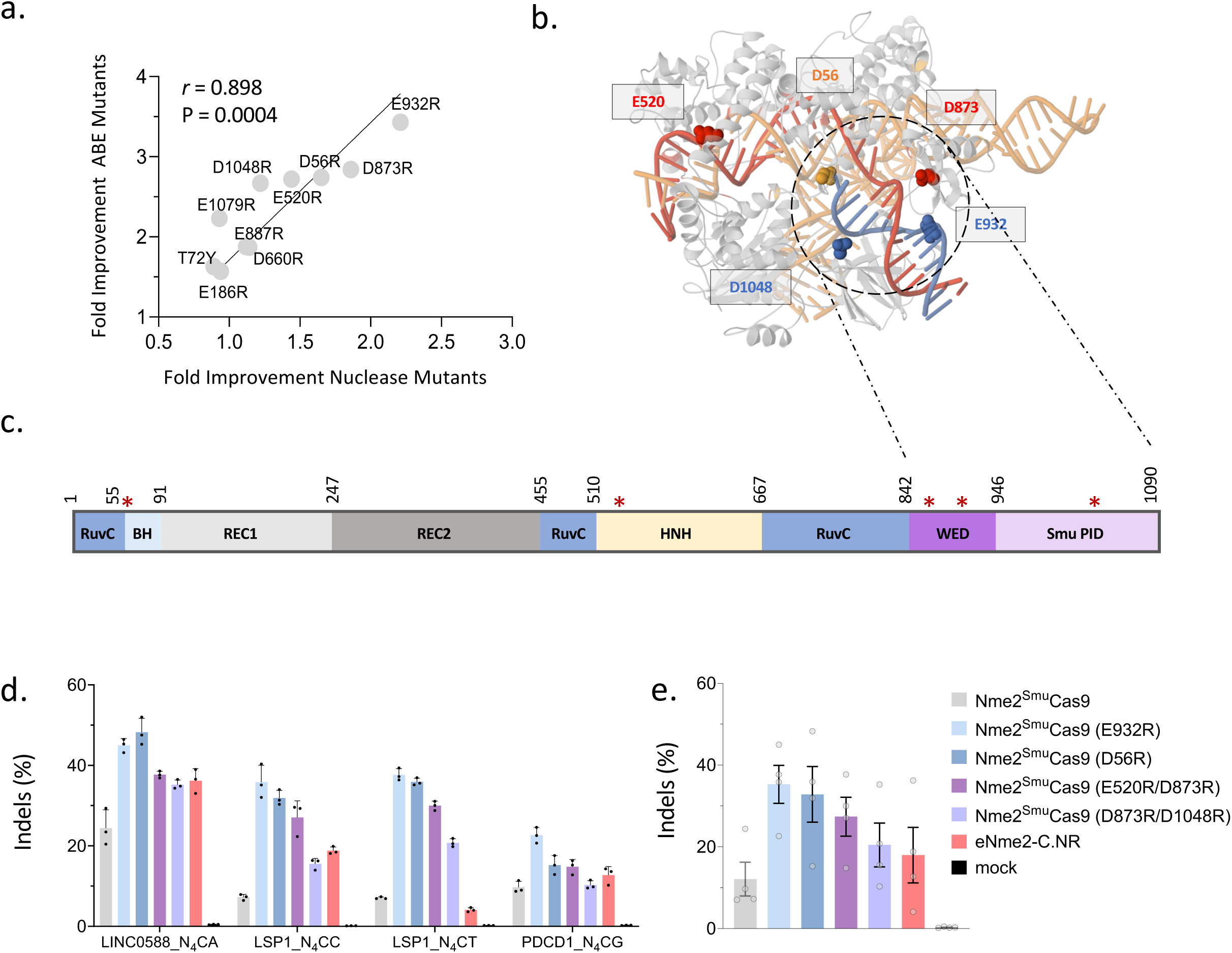
Arginine mutagenesis improves Nme2^Smu^Cas9 nuclease and BE activity. (**a**) Fold improvement of top 10 single-arginine mutants for Nme2^Smu^Cas9 nuclease and Nme2^Smu^-ABE8e-i1 vs. WT variants. Data represent mean activities of four guide RNAs activating either the TLR-MCV reporter (nuclease activity) or mCherry ABE reporter (ABE activity) at N_4_CC, N_4_CT, N_4_CA or N_4_CG PAM targets. Editing efficiencies were measured by flow cytometry (n = 2-3 biological replicates per guide; compiled from **Supplementary Figures 2a-2c**). r = Pearson correlation. (**b**) Homology model of Nme2^Smu^Cas9/sgRNA/DNA ternary structure and (**c**) linear domain organization (not drawn to scale) depicting the relative positions of the top 5 arginine mutants (red asterisks) within Nme2^Smu^Cas9. (**d**) Nuclease editing by Nme2^Smu^Cas9 (WT and E932R, D56R, E520R/D873R, and D873R/D1048R variants) and eNme2-C.NR at endogenous HEK293T genomic loci. The editing efficiency for each target was plotted. Editing efficiencies were measured by amplicon deep sequencing (n = 3 biological replicates; data represent mean ± SD). (**b**) Data from (a) were aggregated and replotted, with each data point representing the editing efficiency of an individual target site, as measured by amplicon deep sequencing (n = 3 biological replicates; data represent mean ± SEM).

We next selected four single or double Nme2^Smu^Cas9 mutants exhibiting increased activity for additional characterization: E932R, E520R/D873R, D873R/D1048R, and D56R. As an initial test, we measured editing efficiencies across endogenous targets within HEK293T cells. Plasmids encoding the Nme2^Smu^Cas9 variants and eNme2-C.NR were transfected along with sgRNA into HEK293T cells (**Fig. 1d, 1e**). In agreement with our screening assays, all Nme2^Smu^Cas9 mutants were more active than the WT nuclease. Furthermore, the E932R and D56R mutants outperformed eNme2-C.NR across all target sites (**Fig. 1d, 1e**).

### Activities of Nme2- and Nme2^Smu^Cas9 and nuclease variants

To better elucidate the activities of the enhanced Nme2^Smu^Cas9 mutants, we assessed editing at a battery of target sites in the context of a self-targeting library. We previously generated HEK293T cells with an integrated 200-member library expressing Nme2Cas9 sgRNAs targeted to adjacent N_4_CN PAM sites ^16^. In this experiment, we analyzed indel activities at 173 target sites, following plasmid transfection and antibiotic selection of cells expressing Nme2^Smu^Cas9 (WT, E932R, D56R and D873R/D1048R) variants, Nme2Cas9 (WT) or eNme2-C.NR. Four days post-transfection, we collected cells and measured editing activities by amplicon sequencing. Across the entire panel of N_4_CN PAM targets, the activity ranks determined by mean indel production tracked consistently with our endogenous target site data: Nme2^Smu^Cas9 (E932R), 12.58% > Nme2^Smu^Cas9 (D56R), 12.48% > Nme2^Smu^Cas9 (WT), 7.78% > Nme2^Smu^Cas9 (D873R/D1048R), 6.76% > eNme2-C.NR, 5.55% > Nme2Cas9 (WT), 1.52% (**Supplementary Fig. 4**). Across the N_4_CD PAM target sites, the trends remained the same with Nme2^Smu^Cas9 (E932R) and (D56R) variants respectively ranking first and second for activity. In contrast, across N_4_CC PAM target sites, Nme2Cas9 (WT) and eNme2-C.NR ranked 3^rd^ and 4^th^ for activity respectively, outperforming Nme2^Smu^Cas9 (WT and D873R/D1048R) (**Supplementary Fig. 4**). Overall, the Nme2^Smu^Cas9 (E932R) and (D56R) variants both demonstrated ∼1.6-fold increase in activity compared to their otherwise WT counterpart.

We considered whether the increases in activity observed with the D56R and E932R mutations would carry over to Nme2Cas9 (WT) ^9^. To test this hypothesis, we measured efficiencies of indel formation using the HEK293T activity library across 191 N_4_CN PAM target sites for Nme2- and Nme2^Smu^Cas9 variants (WT, E932R or D56R) in addition to eNme2-C.NR (**Fig. 2a**). As expected, eNme2-C and the Nme2^Smu^ PID-swapped variants demonstrated the highest editing efficiencies at N_4_CD PAM target sites, while Nme2Cas9 variants exhibited minimal activities at these PAMs (**Fig. 2a**). Conversely, across target sites bearing canonical N_4_CC PAMs, Nme2Cas9 variants exhibited mean editing efficiencies of 3.47% (WT), 9.2% (E932R) and 7.05% (D56R), respectively, yielding a 2.6- and 2-fold increase in activity in comparison to the WT protein (**Fig. 2a**). When comparing Nme2Cas9 vs. Nme2^Smu^Cas9 variants with the same background mutation, the WT PID effector still performed best at N_4_CC PAM target sites (**Fig. 2a**). These results indicate that the enhancing mutations improve the editing activity of Nme2^Smu^Cas9 and extend to Nme2Cas9, enabling enhanced genome editing potential at a range of single- and double-cytidine PAM target sites.

**Figure 2.**
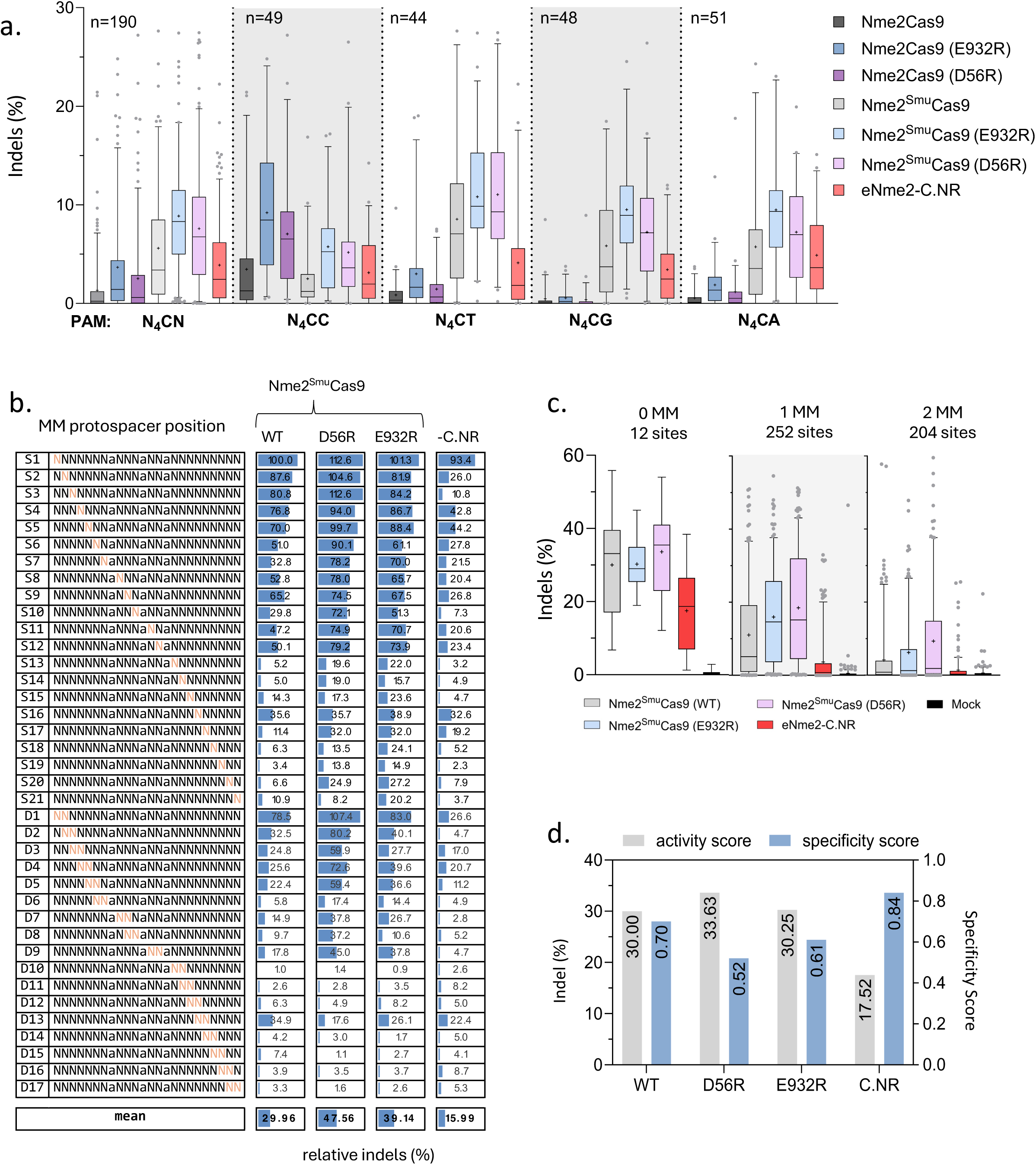
Activities and specificities of Nme2- and Nme2^Smu^Cas9 nuclease variants. (**a**) Nuclease-induced indels in the nuclease experimental panel 2 of the guide-target activity library following plasmid transfection of Nme2Cas9 (WT and E932R, D56R, E520R/D873R variants), Nme2^Smu^Cas9 (WT and E932R, D56R, and E520R/D873R variants) or eNme2-C.NR into HEK293T cells with integrated guide-target sites harboring N_4_CN PAMs. The editing efficiencies for 190 target sites were plotted. (**b**) Average indel frequencies of Nme2^Smu^Cas9 or eNme2-C.NR across single-(*S*) or di-nucleotide (*D*) mismatched target sites within the guide-target mismatch library. Activities for each mismatched target were normalized to the activity of their respective perfectly matched target site. Orange nucleotides represent the protospacer position(s) of the transversion mutation(s) present within the mismatched target site. (**c**) Bulk indel frequencies of the nuclease variants within the mismatch library for 12 perfectly matched target sites (0 MM), 252 single-mismatched target sites (1 MM), or 204 double-mismatched target sites (2 MM). (**d**) Indel vs. specificity scores for Nme2^Smu^Cas9 variants and eNme2-C.NR across the mismatched guide-target library. Indel efficiency was compiled from data for the 12 perfectly matched target sites (0 MM) in (**c**). The specificity score was calculated as one minus the tiled mismatched editing mean in (**b**), normalized to a scale of one to 100. Data were measured by amplicon sequencing (n = 3 biological replicates; boxplots represent median and interquartile ranges; whiskers indicate 5th and 95th percentiles and the cross represents the mean).

### Specificities of Nme2- and Nme2^Smu^Cas9 and nuclease variants

Following the characterization of nuclease activity, we next examined the specificities of the enhanced Nme2Cas9 and Nme2^Smu^Cas9 variants. For this experiment, we developed a library to assess the mismatch tolerance profile of Nme2Cas9 variants at varied positions across its protospacer (**Fig. 2b**). The library consisted of 12 guide RNAs, paired with either a perfectly matched protospacer target or mismatched targets. Each of the 12 guide RNAs were paired with a total of 40 target sites (two identical perfectly-matched targets, 21 single-nucleotide mismatches, and 17 dinucleotide mismatches), for a total of 480 library members (**Fig. 2b**). To mitigate potential PAM-specific effector preferences, three guides per possible N_4_CN PAM sequences were used. In anticipation of using the library for ABE specificity experiments, we incorporated adenines into protospacer positions A8, A12 and A15 and held them constant, as they fall within the editing windows of previously validated Nme2Cas9 derived base editors^16,18,20^.

After Tol2 transposon-mediated integration of the mismatch library into HEK293T cells ^16,31^, we assessed the mismatch tolerance profiles of Nme2Cas9, Nme2^Smu^Cas9, their derivatives, and the eNme2-C.NR nuclease. Following nuclease plasmid transfection and selection, we measured editing via deep sequencing at an average depth of ∼2,300 reads per library member (**Fig. 2b**). At the 12 perfectly matched N_4_CN PAM target sites, the Nme2^Smu^Cas9 variants performed comparably, with mean indel formation ∼30%, while eNme2-C.NR was slightly less efficient, inducing ∼17% indels on average (**Fig. 2c**). Similar trends were observed at the three N_4_CC target sites, with Nme2- and Nme2^Smu^Cas9 arginine mutants (E932R and D56R) performing comparably to or better than their WT counterparts. eNme2-C.NR again had the lowest rates of indel formation for the perfectly matched N_4_CC PAM target sites. In line with previous reports for Nme2Cas9 and other Cas9 effectors, single and double mismatches within the protospacer decreased editing efficiencies ^9,22^, with double mismatches having a more pronounced effect than single mismatches (**Fig. 2b, 2c, Supplementary Fig. 5a**), as expected.

Next, we turned our attention to relative efficiencies of indel formation at mismatched targets (compared to their perfectly matched counterparts) across protospacer positions. In agreement with previous reports ^9,22^, all nucleases tested had general trends for increased tolerance to mismatches at the PAM-distal ends of the protospacer, and decreased mismatch tolerance within the PAM-proximal or seed regions of the protospacer (**Fig. 2b**, **Supplementary Fig. 5a**). Using the mismatch library data, we calculated nuclease specificity scores that enable comparisons between the activity and specificity of the respective editors (**Fig. 2d**, **Supplementary Fig. 5b**). As expected, we observed a negative correlation between indel efficiency and specificity. For example, eNme2-C.NR was the most sensitive to mismatches with the highest specificity score (∼0.8) and the lowest indel efficiency (∼18%), whereas Nme2^Smu^Cas9 (D56R) had the lowest specificity score (0.52) and the highest mean indel efficiency (∼34%) (**Fig. 2d**, **Supplementary Fig. 5b**).

### Optimizing linker lengths for domain-inlaid Nme2^Smu^-ABEs

In our initial development of domain-inlaid Nme2Cas9 BEs ^16^, the larger SmuCas9 PID added 24 bp [8 amino acids (AA)] to the length of the ORF. Consequently, while WT PID Nme2-ABEs could be packaged within a single AAV vector, our design using the 251bp U1a promoter and Nme2^Smu^Cas9-ABEs exceeds the ∼5kb packaging limit of AAV (**Fig. 3a**). Although the use of alternative short promoters, such as EF-1α short (EFS, 212bp) ^10^ could enable packaging of these larger ABEs, we decided to augment this strategy with a more generalized approach to minimize the overall size of the editing reagent. In our initial design of the domain-inlaid Nme2-ABEs, we introduced 20AA linkers between the N- and C-termini of the inserted deaminase domain and the nickase Cas9 (**Fig. 3b**). Previous research has demonstrated that optimization of linker property and length is important for engineering Cas9 fusion proteins ^4,5,32,33^. We anticipated that truncation of the N- and C-terminal linkers flanking the inlaid deaminase domains would enable a reduction in overall protein size to facilitate single-vector AAV packaging (**Fig. 3a, 3b**).

**Figure 3.**
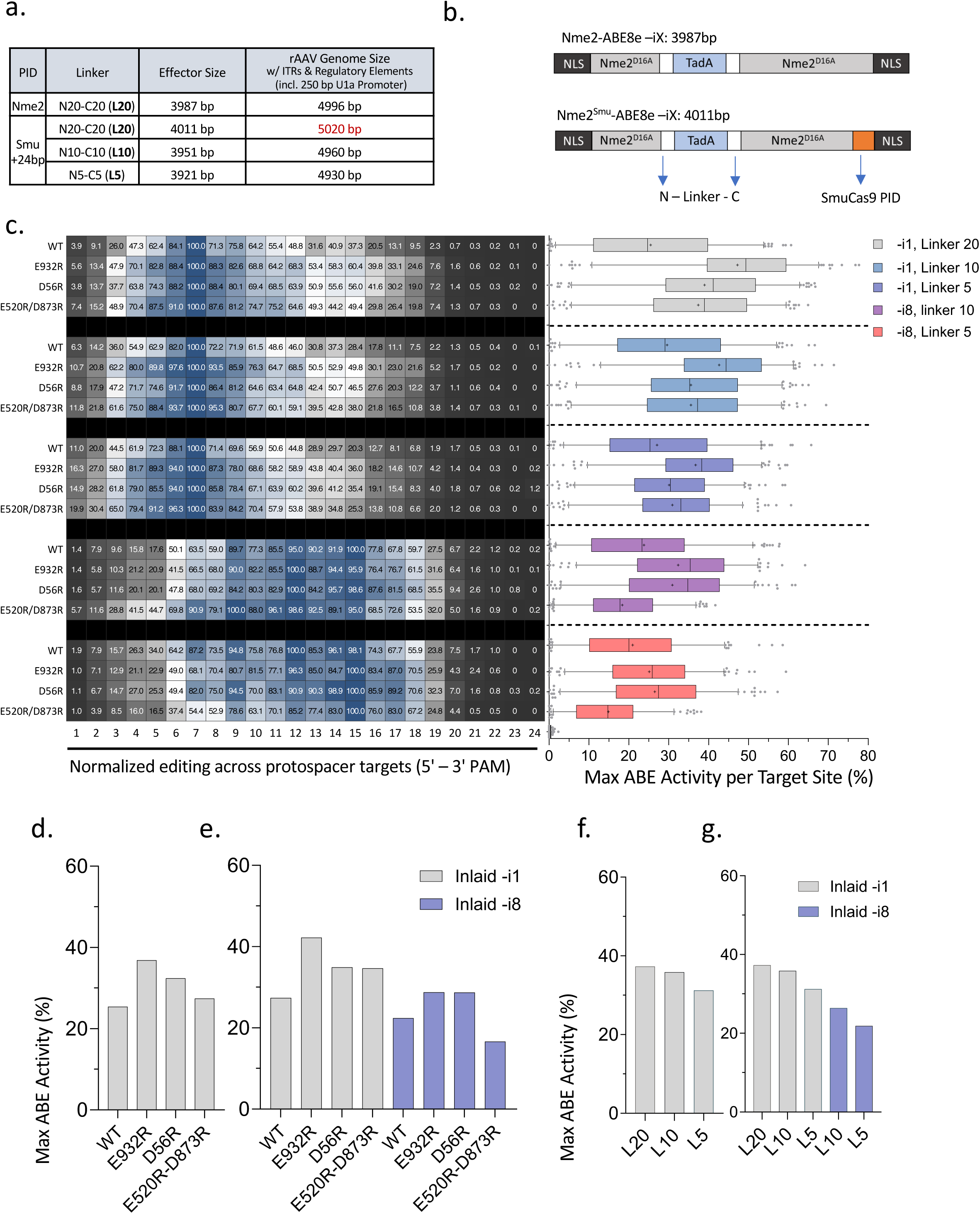
Editing windows and activities of domain-inlaid Nme2^Smu^-ABE8e variants. (**a**) Table depicting rAAV genome size in bp for respective domain-inlaid editors with linker variants and associated regulatory elements (right). Regulatory and coding elements for a representative all-in-one AAV vector include ITRs, U1a promoter, ABE8e editor, U6 promoter, and sgRNA cassette. (a) Cartoon schematic depicting open reading frame length (in bp) of domain-inlaid Nme2-ABE8e with Nme2Cas9 PID (left, top) or Nme2^Smu^-ABE8e with the SmuCas9 PID (left, bottom) with 20AA linkers flanking the N- and C-termini of Tad8e. (**c**) Assessment of editing windows and activities from ABE experimental panel 1 of the guide-target activity library (183 sites) for Nme2^Smu^-ABE8e –i1 or –i8, with or without arginine mutations (E932R, D56R, E520R/D873R), in combination with varied deaminase linker lengths (L20, L10, L5). Following plasmid transfection of the ABE variants into HEK293T cells with the integrated guide-target library, editing activities were measured by amplicon sequencing. Left: average editing windows across the target sites, normalized on a scale of 0 – 100 (%) normalized to the adenine position with the highest observed edited efficiencies within the window. Right: editing activities at the maximally edited adenine for each target were plotted (n = 3 biological replicates; boxplots represent median and interquartile ranges; whiskers indicate 5th and 95th percentiles and the cross represents the mean). (**d-g**) Summary data from self-targeting library maximal activity, aggregated from (**a**). (**d**) Nme2^Smu^-ABE8e and arginine mutant activity independent of domain insertion site and linker length, or (**e**) separated by the position of domain insertion. (**f**) Nme2^Smu^-ABE8e and linker variant activity independent of domain insertion site and arginine mutation, or (**g**) separated by the position of domain insertion.

To probe the effects of linker minimization, we designed and cloned a panel of domain-inlaid Nme2^Smu^-ABE8e (WT) plasmids in -i1 or -i8 formats with combinations of N- or C-terminal linker lengths (Nx - Cx) ranging from 20 to 5 AAs in 5AA steps (**Supplementary Fig. 6**). In addition to the 8 combinations of Nx – Cx linkers ranging from 20 to 5 AA, we included a N0 - C0 linker variant as an extra test subject and the N20-C20 variant as a benchmark. Following testing in HEK293T cells at four endogenous target sites, we observed that most of the linker variants performed comparably to the original N20-C20 linker for both -i1 and -i8 domain-inlaid ABE8e formats (**Supplementary Fig. 6**). With marginal impact on the editing activity at the tested targets, linker minimization of domain-inlaid Nme2^Smu^-ABEs will facilitate single-vector AAV packaging.

To more closely examine size-minimized Nme2^Smu^-ABE effectors, we combined the N10-C10 (L10) and N5-C5 (L5) linker variants with the top-performing enhancing mutations (E932R, D56R, E520R/D873R and D873R/D1048R). Using the same endogenous target sites from our nuclease test, we transfected plasmids encoding combinations of the Nme2^Smu^-ABE8e -i1 or -i8, the L5 or L10 linkers, and the Arg mutants for initial activity testing (**Supplementary Fig. 7a, 7b**). In addition to our domain-inlaid variants, eNme2-C with its original NLS and linker architecture was used for comparison ^18^. Across the four target sites, we observed that all combinations of Tad8e insertion sites, linkers, and mutations were functional with varied rates of activity depending on the combination (**Supplementary Fig. 7a, 7b**). When compared to eNme2-C, the domain-inlaid editors performed comparably across the four target sites on average; however, there were target site-specific differences in activity (**Supplementary Fig. 7a, 7b**). Overall, independent of the site of deaminase insertion or linker length, the E932R and D56R mutations ranked first and second with 1.7- and 1.4-fold activity increases, respectively, compared with their non-mutated counterparts (**Supplementary Fig. 7c**). When reassessing the data for each deaminase insertion site and arginine mutation, independent of linker length, we observed differences in the effect of different effector mutants. For example, in the domain-inlaid -i1 format, E932R and the E520R/D873R mutations ranked first and second for activity, respectively, while in the -i8 format the top two performers were the E932R and D56R mutations (**Supplementary Fig. 7d**).

### Evaluating size-optimized and activity-enhanced domain-inlaid Nme2^Smu^-ABE8e effectors

We next used the guide-target library to investigate the impact of arginine mutation and shortened linkers on the editing windows and activities of domain-inlaid Nme2^Smu^-ABE8e variants. For this experiment we assessed the editing efficiencies for a range of Nme2^Smu^-ABE8e variants across 183 target sites within the library. The effector variants included in this panel consisted of a combination of the following domain-inlaid Nme2^Smu^-ABE8e formats: -i1 or -i8 inlaid architecture, WT or arginine mutants (E932R, D56R, or E520R/D873R), and deaminase linker lengths (L20, L10, L5). For WT Nme2^Smu^-ABE8e editors not bearing arginine mutations, linker variation had minimal impact on the editing window (positions exhibiting activity >50% of the maximum editing efficiency of any adenine within the window) (**Fig. 3c**). A small increase in the mean maximum observed activity were observed for these WT linker variants. For example, Nme2^Smu^-ABE8e-i1 (WT) with L20, L10 or L5 linkers had editing windows spanning protospacer nucleotide (nt) positions 4-11, +/-1 nt. Simultaneously, the mean maximum activity marginally increased with L10 performing the best (∼29%) and L20 the worst (∼25%) (**Fig. 3c)**. In contrast, comparison of the truncated linkers with the same arginine mutant background slightly reduced activity. For instance, linker variants L20, L10 and L5 of Nme2^Smu^-ABE8e-i1 (E932R) had mean maximum activities of approximately 47%, 43% and 37% respectively (**Fig. 3c**). A similar trend was also observed with the -i8 architecture in the background of the E932R and D56R mutations (**Fig. 3c**). Despite this, arginine mutations generally had positive effects on domain-inlaid Nme2^Smu^-ABE8e’s for both the -i1 and -i8 architectures. The effects of the enhancing mutations followed a consistent trend for both the editing window and activity. For example, across the inlaid -i1 linker variants, the E932R mutation led to a widened window accompanied by a considerable increase in activity (+4nt window and 1.9-fold activity in the -i1, L20 format) (**Fig. 3c**). Installation of D56R and E520R/D873R also led to editing window and activity increases for Nme2^Smu^-ABE8e-i1 editors, with these changes being less pronounced (**Fig. 3c)**. In the -i8 architecture, both the E932R and D56R improved activity ∼1.4-fold compared to the WT effector. By contrast, the D520/E873R double mutant performed poorly in the -i8 architecture, exhibiting comparable or lower activity than the WT counterpart (**Fig. 3c**).

Overall, within the library editing experiments, the E932R and D56R mutations resulted in the highest increases in activities (∼1.5- and 1.3-fold respectively) when assessing editing outcomes independent of the deaminase insertion site or linker length (**Fig. 3d**). Importantly, although the E520R/D873R double mutant in the -i1 architecture performed comparably to the D56R substitution, it performed poorly for the -i8 architecture independent of the linker used (**Fig. 3e**).

In summary, the installation of arginine mutants in the domain-inlaid Nme2^Smu^-ABE8e variants enhanced their activities at endogenous target sites and in the context of guide-target libraries. Additionally, when considering linker length for applications without cargo size limits, the L20 variants are likely the variant of choice (**Fig. 3c, 3f, 3g**) based upon their marginally increased activities. Nevertheless, when applying domain-inlaid Nme2^Smu^-ABEs in size-constrained applications, the minimized linkers provide additional flexibility, with minimal losses in overall activity (**Fig. 3f, 3g**).

### Specificities of size-optimized and activity-enhanced domain-inlaid Nme2^Smu^-ABE8e effectors

We repeated the mismatch library experiments to determine the specificities of the size- and activity-optimized domain-inlaid Nme2-ABE8e variants. The effector variants included a combination of the following domain-inlaid formats: Nme2- or Nme2^Smu^Cas9, -i1 or -i8 inlaid architecture (with L10 linkers), WT or arginine mutants (E932R and D56R), and eNme2-C as an additional benchmark.

At the 12 perfectly matched N_4_CN target sites, base editors with arginine mutations outperformed their WT counterparts. Inlaid architectures with the -i1 format performed better than those in the -i8 format, and all domain-inlaid Nme2^Smu^-ABE8e variants induced higher editing efficiencies than eNme2-C (**Fig. 4a**). Similar to the nuclease panel, we observed an inverse correlation with editing efficiency when more mismatches are present, and a similar trend of decreased mismatch tolerance within the seed region (protospacer positions 15-24 counting from the 5’ end of the guide (**Fig. 4b**). Finally, the rankings of specificity scores were also similar to those of the nucleases; eNme2-C exhibited the best specificity and worst editing activity, whereas Nme2^Smu^-ABE8e (D56R) mutants in the -i1 and -i8 architectures had the best activities and worst specificity scores (**Fig. 4b**). Another trend we observed was the unusually high specificity of the -i8 inlaid variants in comparison to the -i1 inlaid variants (**Fig. 4b**). In some cases, -i8 editors exhibited some of the best activities across the 12 perfectly matched target sites with specificity scores (∼0.7), approaching that of eNme2-C (which had the lowest overall ABE activity) (**Fig. 4c**). Assessment of Nme2- and Nme2^Smu^-ABE8e variants with only the N_4_CC targeting guides within the library revealed similar trends to the entire N_4_CN mismatch library (**Supplementary Fig. 8a**). However, Nme2Cas9 variants were generally more effective than Nme2^Smu^ variants at the three N_4_CC perfectly matched targets, though with similar or better specificity scores (**Supplementary Fig. 8b**).

**Figure 4.**
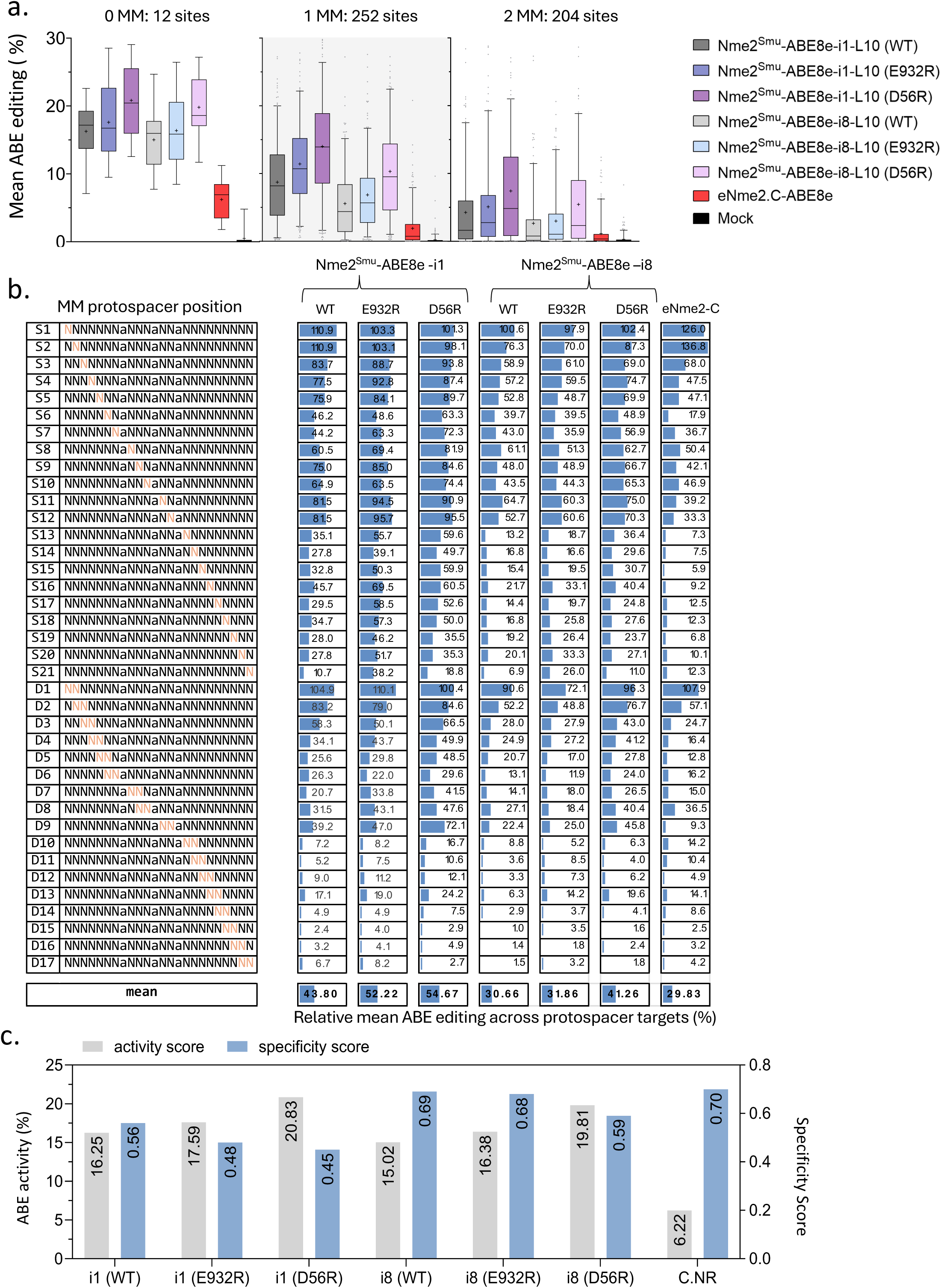
Specificities of domain-inlaid Nme2^Smu^-ABE8e variants. (**a**) Mean A-to-G editing efficiency across the targets within the mismatch library for domain-inlaid Nme2^Smu^-ABE8e variants or eNme2-C. Data was separated by the number of mismatches between guide and target site: 12 perfectly matched sites (0 MM), 252 single-mismatched sites (1 MM), and 204 double-mismatched sites (2 MM). Each data point represents the average A-to-G editing observed across a protospacer of an individual library member. (**b**) Mean A-to-G editing frequencies of domain-inlaid Nme2^Smu^-ABE8e variants or eNme2-C across single-(S) or di-nucleotide (D) mismatched target sites within the guide-target mismatch library. Activities for each mismatched target were normalized to the mean efficiency of their respective perfectly matched target site. Orange nucleotides represent the protospacer position(s) of the transversion mutation(s) present within the mismatched target site. (**d**) ABE activity vs. specificity scores for base editing variants in (**a-b**) across the mismatched guide-target library. ABE activity was compiled from editing data for the perfectly matched target sites (0 MM) in (**a**). The specificity score was calculated as one minus the tiled mismatched editing mean in (**b**), normalized to a scale of one to 100. Data were measured by amplicon sequencing (n = 3 biological replicates; boxplots represent median and interquartile ranges; whiskers indicate 5th and 95th percentiles and the cross represents the mean).

### Testing narrow-window deaminases with the domain-inlaid Nme2^Smu^-ABE architecture

Although we were able to improve the performance of domain-inlaid Nme2^Smu^-ABE8e, in cases where bystander editing is unavoidable, wide editing windows could become problematic. Engineered variants of TadA8e have been established that narrow the editing window by altering residues within or near the catalytic pocket of the deaminase ^34,35^. We tested both Tad9 ^34^ and Tad9e ^35^ variants in combination with our compact Nme2^Smu^-ABE architectures using the self-targeting library. For this experimental panel, we assessed the editing windows and activities across 193 target sites (**Supplementary Fig. 9**). In our hands, the Tad9 ^34^ variants failed to exhibit editing activity above background (**data not shown**). The other Nme2^Smu^-ABE variants tested included a combination of variants: Tad8e or Tad9e deaminase, -i1 or -i8 inlaid architecture, arginine mutants (E932R or D56R), and shortened deaminase linker lengths (L10, L5). Analysis of ABE editing outcomes for the Tad9e effectors confirmed that they have substantially smaller editing windows. This came at the cost of editing activity, a trend consistent with all formats tested (**Supplementary Fig. 9**). For example, the editing window and activity of Nme2^Smu^-ABE8e (L10, E932R), spanned nucleotides 3 – 15 with mean maximum activity of ∼30% (**Supplementary Fig. 9**). In contrast, the Nme2^Smu^-ABE9e (L10, E932R), exhibited an editing window across positions 5-10, with ∼15% maximum activity (**Supplementary Fig. 9**). Likewise, Nme2^Smu^-ABE9e (-i8, L10, E932R) exhibited a narrower window (pos. 10-18) than its ABE8e counterpart, though with a further shift to editing PAM-proximal nucleotides (**Supplementary Fig. 9**). These results demonstrate that Tad9e is compatible with domain-inlaid Nme2^Smu^-ABEs and can increase their precision. Although ABE9e effectors exhibited reduced activity in the domain-inlaid format, in cases where bystander editing must be avoided, they represent a potentially valuable option in base editing reagent.

### Assessment of arginine mutations with WT PID domain-inlaid Nme2-ABEs

Next, we wanted to determine whether the arginine mutations also enhance Nme2Cas9-derived ABEs that natively target an N_4_CC PAM, and how they compare to Nme2^Smu^ chimeric effectors. In addition to the arginine mutations, we also tested whether Nme2Cas9-ABEs (WT PID) were compatible with the minimal linker format (L10). We first constructed plasmids expressing domain-inlaid Nme2-ABE8e effectors in the following formats: inlaid architecture (-i1 and -i8), the L10 linker, and the E932R or D56R mutants. After cloning, we used the self-targeting library to compare domain-inlaid Nme2-ABEs, Nme2^Smu^-ABEs, and eNme2-C (used as a benchmark) across 192 N_4_CN targets (**Supplementary Fig. 10a**).

We first assessed the editing windows and activities of the Nme2-ABE8e derivatives across the 49 canonical N_4_CC PAM target sites within the self-targeting library (**Supplementary Fig. 10a**). While editing windows were somewhat more variable among this subset of target sites, the assay adequately captured the previously described window of eNme2-C ^18^. For editing window analysis, we noticed that in most cases, arginine mutants led to an increase in the editing windows for both Nme2- and Nme2^Smu^-ABEs, a trend consistent with the previous experiments (**Fig. 3a, Supplementary Fig. 10a**). For example, the observed editing window for Nme2-ABE-i1 (WT) and the E932R variant ranged from positions 3-13 and 3-16 respectively (**Supplementary Fig. 10a**). Although the windows for Nme2-ABE-i8 (WT and E932R) didn’t change dramatically in size (spanning positions 8-17 and 9-18 respectively), the arginine mutants increased editing near window boundaries (**Supplementary Fig. 10a**). Again, when installing the arginine mutants, we observed higher editing activities associated with an increased window size. For illustration, the mean maximum activities of Nme2Cas9-ABEs with the inlaid -i1 format were approximately 21% (WT), 31% (E932R), and 26% (D56R) (**Supplementary Fig. 10a**). Similarly, Nme2Cas9 ABEs with the inlaid -i8 format had mean maximum activities of approximately 15% (WT), 27% (E932R), and 20% (D56R) (**Supplementary Fig. 10a**). Altogether, at N_4_CC PAM targets, the E932R and D56R mutants increased editing of the domain-inlaid Nme2-ABE8es over their non-mutant counterparts by 1.6- and 1.2-fold. Furthermore, at all N_4_CC PAMs, the average activities of Nme2-ABE8e effectors were higher than the PID-chimeric Nme2^Smu^-ABEs and eNme2-C.

We also assessed the editing activities of the inlaid Nme2-ABE8e effectors at N_4_CD PAM target sites. Similar to our nuclease experiments, installation of the E932R and D56R mutants increased the activities of inlaid Nme2-ABEs at non-canonical (N_4_CD) PAM target sites (**Supplementary Fig. 10b**). Intriguingly, although inlaid Nme2^Smu^-ABE8e effectors outperformed the Nme2Cas9 (WT PID) counterparts at all N_4_CD PAM targets, the Nme2Cas9-ABE-i1 (E932R) variant approached similar activity levels across N_4_CT targets, albeit with less consistency than the corresponding PID-swapped variant [Nme2 PID, (∼20% ± SD of 14%) vs. Smu PID (∼24% ± SD of 10%)] (**Supplementary Fig. 10b**). These results demonstrate that the enhancing mutations are also applicable to domain-inlaid Nme2-ABE8e effectors and potentially relax their PAM requirements.

### Activities and windows of engineered Nme2Cas9 variants in various ABE8e formats

While this work was in progress, other groups reported on alternative engineering avenues to increase the efficiency and alter the PAM targeting scope of Nme2Cas9 editing systems towards N_4_CN PAM targets ^18,36^. We thus wanted to compare these PAM-relaxed nickase Cas9 domains for base editing applications at N_4_C PAM target sites. The Cas9 variants in this panel consisted of Nme2 (WT), Nme2^Smu^ (WT), and their mutant derivatives (E932R, E932R/D56R), in addition to eNme2-C and a rationally engineered Nme2Cas9 variant with enhanced DNA unwinding capabilities iNme2 ^36^. Since iNme2 had no mutations within the PID domain, we also cloned an iNme2^Smu^ Cas9 derivative, to determine whether the alternate PID would further enhance efficiency at N_4_CD PAM target sites. In this experiment, we tested the engineered Cas9 base editing variants in either the N-terminal or domain-inlaid -i1 (linker L10) ABE8e architectures using the aforementioned activity library across 181 N_4_CN target sites (**Fig. 5**, **Supplementary Figs. 11, 12**).

**Figure 5.**
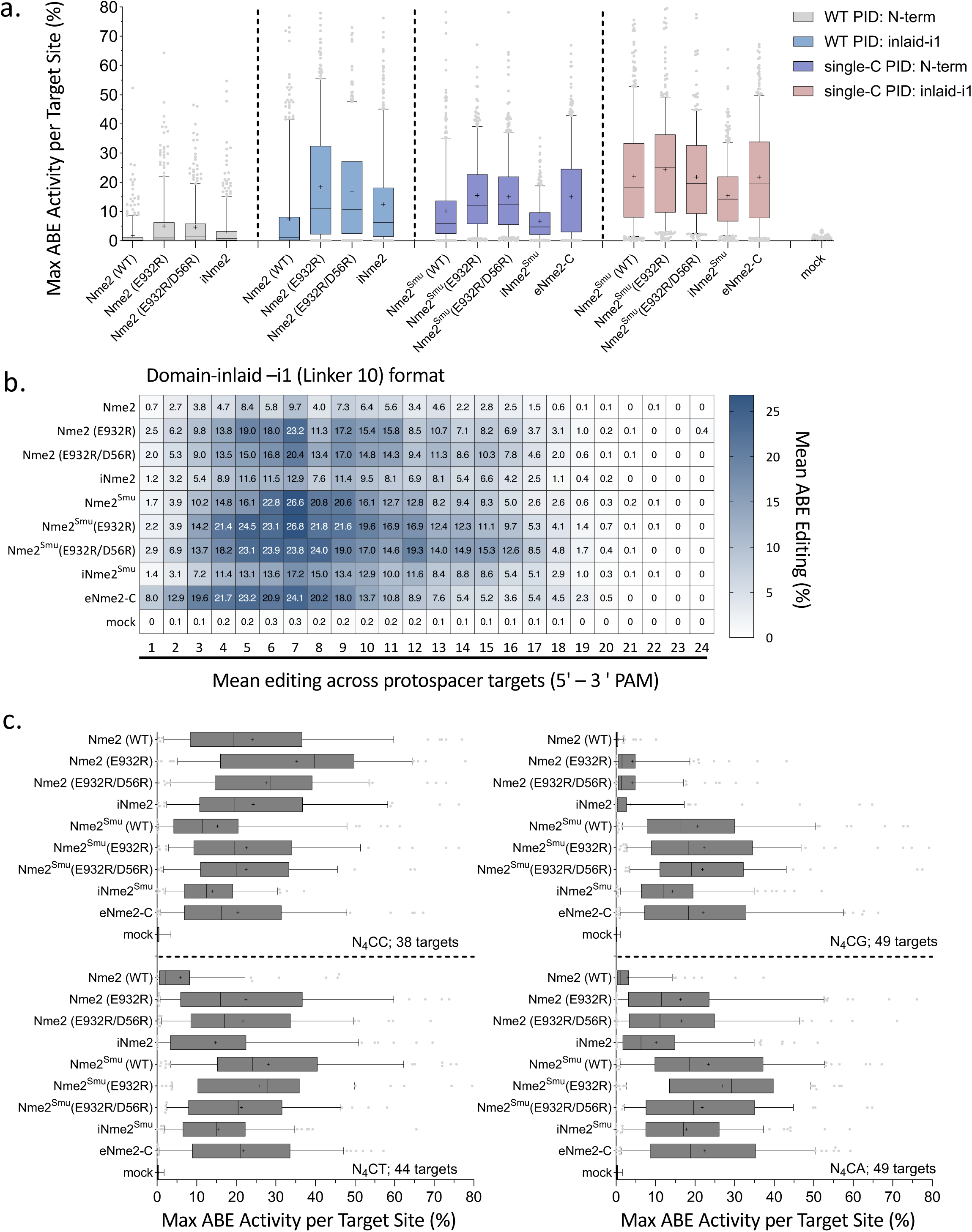
Activities and editing windows of engineered Nme2Cas9 variants in various ABE8e formats. Assessment of editing activities and windows from experimental panel 4 of the guide-target activity library (181 target sites) for Nme2-, Nme2^Smu-^, iNme2-, iNme2^Smu-^, and eNme2-C variants in either the N-terminally fused or inlaid-i1 (linker 10) format. (**a**) Efficiency at the maximally edited adenine for each target was plotted for all N_4_CN PAM target sites. ABEs with a WT Nme2Cas9 PID (WT PID) or N_4_CN targeting PID (single -C PID) are depicted by color. (**b**) Mean A-to-G editing activities and windows across protospacer positions in the activity guide-target library for engineered Nme2-ABE8e variants in the domain-inlaid -i1 (linker 10) format. (**c**) Data in (**a**) separated by target site PAM identity (N_4_CC, N_4_CT, N_4_CG, N_4_CA) for the engineered Nme2-ABE8e variants in the domain-inlaid -i1 (linker 10) format. The editing efficiency of the maximally edited adenine for each target was plotted. The number of target sites per PAM is indicated (n = 3 biological replicates; boxplots represent median and interquartile ranges; whiskers indicate 5th and 95th percentiles and the cross represents the mean).

In all cases, Nme2^Smu^Cas9 PID variants performed better than their WT PID counterparts across the N_4_CN PAM targets. In a similar fashion, the domain-inlaid -i1 architecture also increased the editing efficiency of all Cas9 variants tested within the panel (**Fig. 5a**). We also observed that the -i1 inlaid architecture mediated shifts in editing windows towards PAM-distal adenines, an effect most prominent with the eNme2-C Cas9 domain (**Fig. 5b, Supplementary Fig. 11**). Since the ABE8e editors in the -i1 inlaid format had the highest activities, we focused our subsequent analyses on these variants. The top-ranking editor for mean maximum ABE activity across all N_4_CN target sites was Nme2^Smu^(E932R) (∼24%), closely followed by Nme2^Smu^ (WT, E932R/D56R) and eNme2-C, with approximate editing efficiencies ∼22% (**Fig. 5a**). iNme2 had average ABE editing activity approximately 12%, while iNme2^Smu^ performed marginally better (∼15%) (**Fig. 5a**).

Next, we looked more closely at the average ABE editing activities of different Cas9 editors when varying the 6^th^ nucleotide of the N_4_CN PAM. For the N_4_CC target sites, Cas9 mutants with the WT Nme2 PID performed best (**Fig. 5c**). For example, Nme2-ABE8e-i1 (E932R) induced mean maximum activity ∼35%, whereas its PID-swapped counterpart had lower activity (∼22%). For these target sites, the Nme2 (E932R/D56R) and iNme2 Cas9 editors ranked second (∼28%) and third (∼24%) respectively (**Fig. 5c**). Top-performing Cas9 variants for mean maximum ABE editing at N_4_CD target sites were Nme2^Smu^ (E932R) > Nme2^Smu^ (WT) > eNme2-C, with respective mean maximum editing rates of ∼25%, ∼24% and 22% (**Fig. 5c**). Across N_4_CT target sites, Nme2^Smu^ (WT) ranked first, whereas Nme2^Smu^ (E932R) performed the best at N_4_CG and N_4_CA target sites (**Fig. 5c**). In agreement with our previous results, Nme2 (E932R) had high activity across N_4_CT target sites (∼22%), ranking third for activity against the other Cas9 editors (**Fig. 5c**). These results establish deaminase domain insertion as a viable approach to improve the general activities for a variety of engineered Nme2Cas9-derived base-editing systems.

## Discussion

In an effort to relax the PAM preference of Nme2Cas9 editors from N_4_CC to N_4_CN, we ^16^ and others ^19,20^ engineered Nme2Cas9 nuclease and BE derivatives with a transplanted PAM interacting domain (PID) from SmuCas9 ^17^. Although active at N_4_CN PAM target sites, the chimeric effectors are often outperformed by Nme2Cas9 at target sites with a canonical N_4_CC PAM. This trend is commonly observed with other PAM-relaxed Cas9 variants ^21,24,23,22,25^. Furthermore, when considering *in vivo* ABE applications, the SmuCas9 PID swap increases the sizes of our domain-inlaid architectures, which are already very close to the packaging capacities of AAV vector capsids.

We continued our advances to optimize Nme2^Smu^ nuclease and ABE systems for activity and single-vector AAV deliverability. We first used arginine mutagenesis to enhance interactions with either sgRNA, TS DNA, or NTS DNA. Of note, the E932R and D56R mutations significantly improved the editing efficiencies of Nme2^Smu^ nuclease systems on all N_4_CN PAM targets. After finding Nme2^Smu^ mutants with improved activity, we transferred them into our domain-inlaid ABE architectures, optimizing for linker length in the process to facilitate single-vector AAV compatibility. Finally, we defined the editing activities, PAM compatibilities, editing windows, and specificities of the resulting editors using endogenous genomic sites as well as self-targeting guide-target libraries. In addition to the wide windows of ABE8e deaminases, we found that ABE9e variants are also compatible with our domain-inlaid ABE architectures, allowing for narrow-window applications (albeit at the cost of somewhat reduced editing efficiency).

As an additional benefit, the activity-enhancing properties of the arginine mutants also extended to WT Nme2Cas9 effectors, increasing their activities at N_4_CC PAM targets. In contrast to the self-targeting library nuclease results (**Fig. 2a**), we observed a particularly stark activity increase at N_4_CW (W = A or T) PAM target sites for domain-inlaid Nme2-ABE-i1 (**Fig. 5c**, **Supplementary Fig. 10b**). This suggests that the enhancing mutations may relax the PAM requirements of this ABE. Nonetheless, in terms of PAM selection, Nme2^Smu^Cas9 effectors are the variants of choice for N_4_CD targets, while Nme2Cas9 effectors likely have the best activity at targets with N_4_CC PAMs. Finally, although inlaid Nme2-ABEs with a WT PID can already be packaged into single AAV vectors ^10,11^, the additional space saved with minimized linkers may further their utility when delivered via AAV vectors.

In terms of alternative engineering approaches for Nme2Cas9 systems, the Liu group used phage-assisted evolution to develop eNme2-C base editing systems ^18^, whereas the Doudna group employed a mixture of rational design and homology modelling focused on WED domain engineering to develop iNme2Cas9 ^36^. Notably, our top-performing arginine mutant (E932R) overlaps with one of the identified amino acid positions observed to improve Nme2Cas9 activity during PACE evolution (E932K for eNme2-C) for nuclease and base editing (Huang et al., 2022); or during a bacterial evolution campaign combined with rational design (E932K) for iNme2Cas9 nuclease ^36^. Biochemical characterization in the iNme2Cas9 study ^36^ points to increased unwinding rates as a prospective mechanism for the enhanced editing activity seen with some nucleic acid interacting mutants (E932R, D873R, D1048R).

Enhanced R-loop formation also potentially explains the PAM relaxation effect observed with WT PID Nme2-ABE8e-i1. iNme2Cas9 was shown to have robust activity at the non-canonical PAMs N_4_CT and N_4_TC at a limited number of target sites, without mutation of residues directly interacting with the PAM ^36^. It is interesting to speculate whether activating mutations such as E932R or those found within the iNme2Cas9 editors could further relax the preferences of Nme2^Smu^Cas9 effectors. Finally, although the activity-specificity balance of Nme2Cas9-derived editors leads to intrinsically reduced off-target editing than many other Cas9 orthologs, the broadening of PAM compatibility paired with improved activity are likely to be inversely correlated with specificity ^37,38^. Additional studies will be necessary to carefully define the off-target profiles of these enhanced Nme2Cas9 and Nme2^Smu^Cas9 variants, especially in cases of therapeutic application.

In summary, we have improved the editing potential of Nme2Cas9- and Nme2^Smu^Cas9-derived editors for nuclease and ABE applications. Altogether, the domain-inlaid Nme2^Smu^-ABE8e’s (-i1 and -i8) with editing windows that enable targeting from positions 4 – 18 (counting from the 5’ end of the protospacer NTS, **Fig. 6a**), in combination with a single-cytidine PAM preference, enables targeting of 88% of adenines within the hg38 reference genome (**Fig. 6b**). The expanded windows of the domain-inlaid variants resulted in a 62% and 19% increase in targeting scope compared to N-terminally fused variants targeting dinucleotide N_4_CC PAMs (Nme2-ABE8e) ^10,11^ or single-cytidine N_4_CN PAMs (eNme2-C) ^18^ (**Fig. 6b**). Furthermore, we were able to port our domain-inlaid designs for use with eNme2-C and iNme2 Cas9 domains. The use of alternately engineered Nme2Cas9 ABE editing systems could prove beneficial when fine-tuning parameters such as editing window, activity, and off-target profile during therapeutic implementation. Overall, the minimal PAM requirements, tunable editing windows, and AAV compatibilities of these enhanced domain-inlaid ABE systems provide greater control over targetable genomic adenines with improved utility for *in vivo* applications.

**Figure 6.**
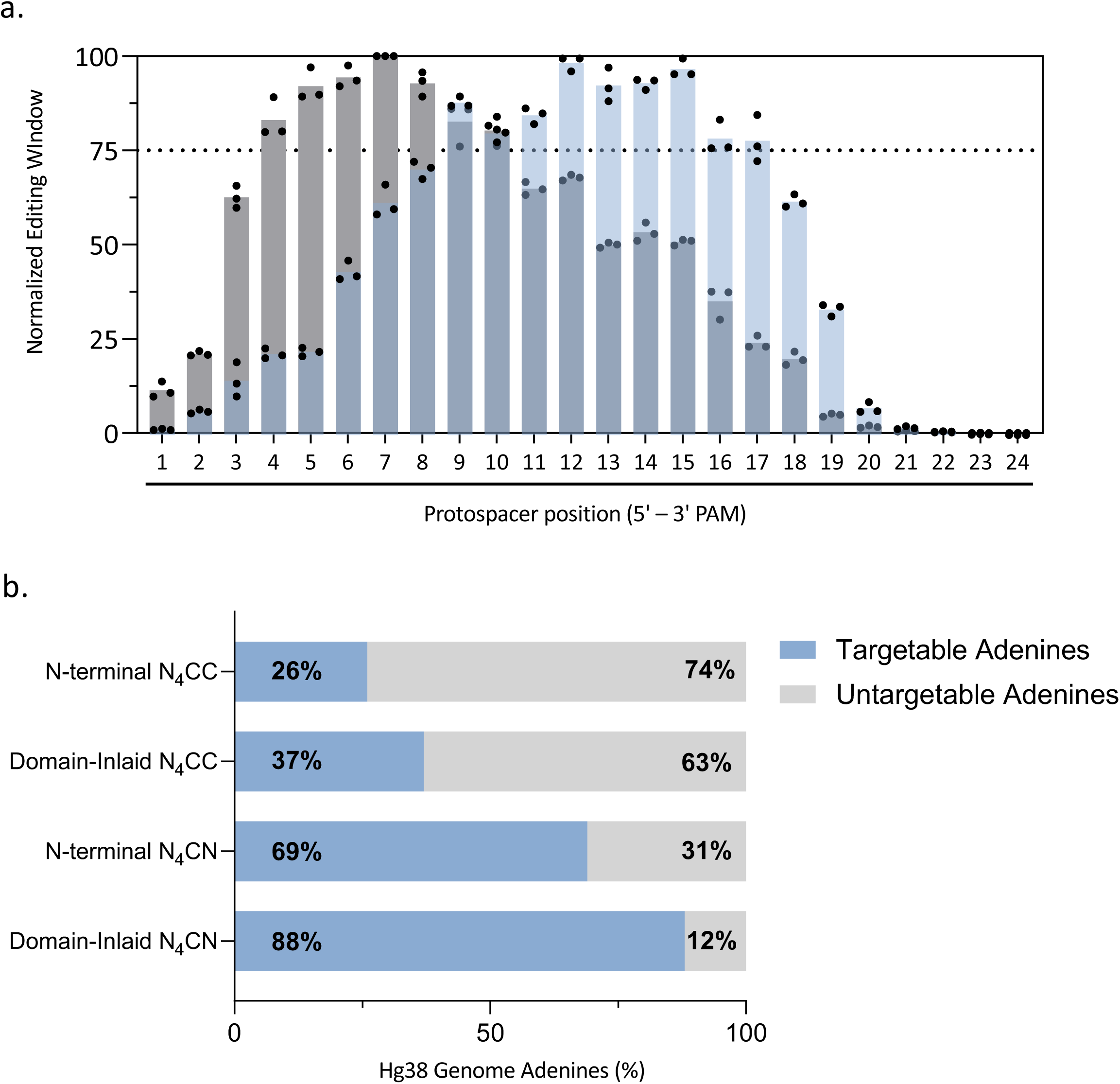
Summary of editing windows and genomic targetable adenines by various Nme2Cas9-derived ABEs. (**a**) Summary of editing windows of Nme2^Smu^-ABE8e-i1 or Nme2^Smu^-ABE8e-i8 with the L10 linker format and E932R mutation. The data represent the normalized editing rates across the window from three independent self-targeting library experimental panels, compiled from Fig. 3a and **Supplementary Figures 9 and 10a**. Each experimental panel consisted of 3 biological replicates. (**b**) Adenines targetable within the hg38 reference human genome by Nme2Cas9-derived ABE8e variants in various formats. Editing windows to calculate the targetable adenines within the reference genome consisted of the previously described window for N-terminally fused Nme2-ABE8e ^10^, or the editing windows observed here with the guide-target library assay for N-terminally fused eNme2-C or domain-inlaid –i1 or –i8 Nme2^Smu^-ABE8e editors from (**a**). Targetable adenine calculations were also made for whether the ABE uses dinucleotide (N_4_CC) or single nucleotide cytidine (N_4_CN) PAMs. Editing activity above 75% of the maximum position in the window was the cutoff criteria for window selection. Code used to generate this data was adapted from ^10^.

## Methods

### Molecular cloning

Nucleotide sequences of key nucleases and base editors described in this manuscript are provided in the **Supplementary Notes**. Effectors used for endogenous target site editing were cloned into a CMV promoter plasmid backbone (Addgene #201510) for expression. Conversely, for the guide-target library experiments, nucleases and base editors were cloned into the p2T-CMV-ABEmax-BlastR backbone (Addgene, #152989) via Gibson assembly. Plasmids expressing nuclease and base editor arginine mutants were generated by introduction of point mutations by site-directed mutagenesis (SDM) oligos in the **Supplementary Oligonucleotides** file, using NEB’s KLD enzyme mix (NEB #M0554S) with the appropriate Nme2Cas9 effector plasmid used as a template. The truncated Nx-Cx linker variants were constructed via Gibson assembly with ssDNA bridge oligos containing the linker of interest, as described in the **Supplementary Notes**.

### Transient transfection for fluorescent reporter and endogenous target site experiments

HEK293T (ATCC #CRL-3216) cells and the fluorescent reporter derivatives described in this manuscript were cultured in Dulbecco’s Modified Eagle’s Medium (DMEM; Genesee Scientific #25-500) with 10% fetal bovine serum (FBS; Gibco #26140079). For transient plasmid transfection, ∼15,000 cells were seeded in 96-well plates and incubated overnight. The following day, approximately 0.75 µl Lipofectamine 2000 (ThermoFisher #11668019) was used to transfect cells with ∼100 ng of the editor plasmid and ∼100 ng of sgRNA according to the manufacturer’s recommended protocols. In all cases cells were incubated at 37°C with 5% CO_2_.

### Flow cytometry

For measurements of editing activities with fluorescent reporters, cells were trypsinized, collected, and washed with FACS buffer (chilled PBS and 3% fetal bovine serum) 72 hours post-transfection. After washing, cells were resuspended in 300 µl FACS buffer for flow cytometry analysis using the MACSQuant VYB system. A total of 10,000 cells were counted for analysis with Flowjo v10.

### Amplicon sequencing and data analysis of endogenous target sites

Amplicon sequencing, library preparation, and analysis were performed as previously described^16^. We used NEBNext Ultra II Q5® Master Mix (NEB #M0544) to amplify genomic DNA for library preparation, followed by pooling and two rounds of left-sided size selection with SPRIselect beads (Beckman Coulter #B23317) and subsequent agarose gel analysis. Pooled amplicons were sequenced on an Illumina MiniSeq system (300 cycles, Illumina sequencing kit #FC-420-1004) following the manufacturer’s protocol. Sequencing data was analyzed with CRISPResso248 (version 2.0.40) in nuclease or BE output batch mode with the following flags: Nuclease (-wc −3, -w 1, -q 30), BE (-w 12, -wc −12, -q 30).

### Transient transfection for guide-target library experiments

Construction and editing experiments for the activity guide-target library assay were as previously described ^16^. In brief,∼200,000 (activity library) or ∼400,000 (mismatch library) cells were plated into 12- or 6-well plates, respectively, in non-selective medium and incubated overnight. The next day, cells were transfected with ∼1.8 μg (activity library) or ∼3.6 μg (mismatch library) of editing effector plasmid using Lipofectamine 2000 following the standard manufacturer-recommended protocol. One day post-transfection, culture media were supplemented with Blasticidin S [10 μg/ml]. After 3 days, genomic DNA was extracted from cells with QuickExtract (Lucigen #QE0905), column-purified (Zymo Research #C1102-50), and used for NGS library preparation.

### Amplicon sequencing and data analysis of library target sites

NGS library preparation was performed as described previously ^16^ and akin to the *Amplicon sequencing and data analysis of endogenous target sites* section listed above, with the following amendments. 1 μg (activity library) or 2 μg (mismatch library) of input DNA was used for library PCRs and sequenced on an Illumina NextSeq 2000 system (200 cycles, Illumina sequencing kit #20046812). Sequencing data for the activity library was demultiplexed using our previously published custom script ^16^ while sequencing data for the mismatch library was demultiplexed with a new custom script (see **Supplemental Code**). Demultiplexed files were analyzed with CRISPResso2 (version 2.0.40) in the standard batch output modes as described above. Library members with <40 reads were omitted from analysis in all samples. The following flags were used: Nuclease (-wc −3, -w 1, -q 10), BE (-w 12, -wc −12, -q 10).

### Statistical analysis

Data analysis and plotting was performed in GraphPad Prism 9.4.0.

## Data Availability

Sequencing data that support the findings of this study will be made available in the NCBI SRA (bioproject #PRJNA1098767) upon publication. Source data for figures are provided within the **Source Data** and **Supplementary Data** files. Sequences of target sites and oligonucleotides (primers, guide-target library oligos) used in this study are provided in **Supplementary Data 9, Oligonucleotides** file. Key plasmids described in this paper are being made available via Addgene. All other data will be made available upon request.

## Supporting information

Inventory of Supplementary Items

## Acknowledgments

We thank members of the Sontheimer, Watts, Wolfe, Xue, and Khvorova laboratories for their time, advice, and feedback during the preparation of this manuscript. Support for this work was provided by the National Institutes of Health (F31GM143879 and R25GM113686 to N.B. and R01GM150273 to E.J.S.), the Rett Syndrome Research Trust (E.J.S. and J.K.W.), and the Leducq Foundation (to E.J.S.).

## Author Contributions

N.B., Z.C., and E.J.S. conceived the study and designed experiments with input from all authors. N.B., A.A., E.J., and Z.C. performed and analyzed cell culture experiments. H.C. and R.P. developed and implemented scripts for computational analysis of sequencing data. N.B. and E.J.S. wrote the manuscript with editing contributions from all co-authors.

## Supplementary Note 1. Nucleotide sequence of key nuclease and base editing constructs described in this manuscript

**Nme2Cas9 (WT)**: <u>BPSV40-NLS</u>, Nme2Cas9, Linkers

MKRTADGSEFESPKKKRKVEDMAAFKPNPINYILGLDIGIASVGWAMVEIDEEENPIRLIDLGVRVFERAEVPKTGDSLAMARRLARSVRR LTRRRAHRLLRARRLLKREGVLQAADFDENGLIKSLPNTPWQLRAAALDRKLTPLEWSAVLLHLIKHRGYLSQRKNEGETADKELGALLKG VANNAHALQTGDFRTPAELALNKFEKESGHIRNQRGDYSHTFSRKDLQAELILLFEKQKEFGNPHVSGGLKEGIETLLMTQRPALSGDAV QKMLGHCTFEPAEPKAAKNTYTAERFIWLTKLNNLRILEQGSERPLTDTERATLMDEPYRKSKLTYAQARKLLGLEDTAFFKGLRYGKDNA EASTLMEMKAYHAISRALEKEGLKDKKSPLNLSSELQDEIGTAFSLFKTDEDITGRLKDRVQPEILEALLKHISFDKFVQISLKALRRIVPLMEQ GKRYDEACAEIYGDHYGKKNTEEKIYLPPIPADEIRNPVVLRALSQARKVINGVVRRYGSPARIHIETAREVGKSFKDRKEIEKRQEENRKDR EKAAAKFREYFPNFVGEPKSKDILKLRLYEQQHGKCLYSGKEINLVRLNEKGYVEIDHALPFSRTWDDSFNNKVLVLGSENQNKGNQTPYE YFNGKDNSREWQEFKARVETSRFPRSKKQRILLQKFDEDGFKECNLNDTRYVNRFLCQFVADHILLTGKGKRRVFASNGQITNLLRGFWG LRKVRAENDRHHALDAVVVACSTVAMQQKITRFVRYKEMNAFDGKTIDKETGKVLHQKTHFPQPWEFFAQEVMIRVFGKPDGKPEFEE ADTPEKLRTLLAEKLSSRPEAVHEYVTPLFVSRAPNRKMSGAHKDTLRSAKRFVKHNEKISVKRVWLTEIKLADLENMVNYKNGREIELYEA LKARLEAYGGNAKQAFDPKDNPFYKKGGQLVKAVRVEKTQESGVLLNKKNAYTIADNGDMVRVDVFCKVDKKGKNQYFIVPIYAWQVA ENILPDIDCKGYRIDDSYTFCFSLHKYDLIAFQKDEKSKVEFAYYINCDSSNGRFYLAWHDKGSKEQQFRISTQNLVLIQKYQVNELGKEIRPC RLKKRPPVREDKRTADGSEFEPKKKRKV

Atgaaacggacagccgacggaagcgagttcgagtcaccaaagaagaagcggaaagtcgaagatatggccgccttcaagcctaacccaatcaattacatcctgggactgga catcggaatcgcatccgtgggatgggctatggtggagatcgacgaggaggagaatcctatccggctgatcgatctgggcgtgagagtgtttgagagggccgaggtgccaaag accggcgattctctggctatggcccggagactggcacggagcgtgaggcgcctgacacggagaagggcacacaggctgctgagggcacgccggctgctgaagagagaggg cgtgctgcaggcagcagacttcgatgagaatggcctgatcaagagcctgccaaacaccccctggcagctgagagcagccgccctggacaggaagctgacaccactggagtg gtctgccgtgctgctgcacctgatcaagcaccgcggctacctgagccagcggaagaacgagggagagacagcagacaaggagctgggcgccctgctgaagggagtggcca acaatgcccacgccctgcagaccggcgatttcaggacacctgccgagctggccctgaataagtttgagaaggagtccggccacatcagaaaccagaggggcgactatagcc acaccttctcccgcaaggatctgcaggccgagctgatcctgctgttcgagaagcagaaggagtttggcaatccacacgtgagcggaggcctgaaggagggaatcgagaccct gctgatgacacagaggcctgccctgtccggcgacgcagtgcagaagatgctgggacactgcaccttcgagcctgcagagccaaaggccgccaagaacacctacacagccga gcggtttatctggctgacaaagctgaacaatctgagaatcctggagcagggatccgagaggccactgaccgacacagagagggccaccctgatggatgagccttaccggaa gtctaagctgacatatgcccaggccagaaagctgctgggcctggaggacaccgccttctttaagggcctgagatacggcaaggataatgccgaggcctccacactgatggag atgaaggcctatcacgccatctctcgcgccctggagaaggagggcctgaaggacaagaagtcccccctgaacctgagctccgagctgcaggatgagatcggcaccgccttct ctctgtttaagaccgacgaggatatcacaggccgcctgaaggacagggtgcagcctgagatcctggaggccctgctgaagcacatctctttcgataagtttgtgcagatcagc ctgaaggccctgagaaggatcgtgccactgatggagcagggcaagcggtacgacgaggcctgcgccgagatctacggcgatcactatggcaagaagaacacagaggaga agatctatctgccccctatccctgccgacgagatcagaaatcctgtggtgctgagggccctgtcccaggcaagaaaagtgatcaacggagtggtgcgccggtacggatctcca gcccggatccacatcgagaccgccagagaagtgggcaagagcttcaaggaccggaaggagatcgagaagagacaggaggagaatcgcaaggatcgggagaaggccgcc gccaagtttagggagtacttccctaactttgtgggcgagccaaagtctaaggacatcctgaagctgcgcctgtacgagcagcagcacggcaagtgtctgtatagcggcaagg agatcaatctggtgcggctgaacgagaagggctatgtggagatcgatcacgccctgcctttctccagaacctgggacgattcttttaacaataaggtgctggtgctgggcagcg agaaccagaataagggcaatcagacaccatacgagtatttcaatggcaaggacaactccagggagtggcaggagttcaaggcccgcgtggagacctctagatttcccagga gcaagaagcagcggatcctgctgcagaagttcgacgaggatggctttaaggagtgcaacctgaatgacaccagatacgtgaaccggttcctgtgccagtttgtggccgatca catcctgctgaccggcaagggcaagagaagggtgttcgcctctaatggccagatcacaaacctgctgaggggattttggggactgaggaaggtgcgggcagagaatgacag acaccacgcactggatgcagtggtggtggcatgcagcaccgtggcaatgcagcagaagatcacaagattcgtgaggtataaggagatgaacgcctttgacggcaagaccat cgataaggagacaggcaaggtgctgcaccagaagacccacttcccccagccttgggagttctttgcccaggaagtgatgatccgggtgttcggcaagccagacggcaagcct gagtttgaggaggccgataccccagagaagctgaggacactgctggcagagaagctgtctagcaggccagaggcagtgcacgagtacgtgaccccactgttcgtgtccagg gcacccaatcggaagatgtctggcgcccacaaggacacactgagaagcgccaagaggtttgtgaagcacaacgagaagatctccgtgaagagagtgtggctgaccgagat caagctggccgatctggagaacatggtgaattacaagaacggcagggagatcgagctgtatgaggccctgaaggcaaggctggaggcctacggaggaaatgccaagcagg ccttcgacccaaaggataaccccttttataagaagggaggacagctggtgaaggccgtgcgggtggagaagacccaggagagcggcgtgctgctgaataagaagaacgcc tacacaatcgccgacaatggcgatatggtgagagtggacgtgttctgtaaggtggataagaagggcaagaatcagtactttatcgtgcctatctatgcctggcaggtggccga

**Nme2Cas9 (E932R)**: <u>BPSV40-NLS</u>, Nme2Cas9, Linkers

MKRTADGSEFESPKKKRKVEDMAAFKPNPINYILGLDIGIASVGWAMVEIDEEENPIRLIDLGVRVFERAEVPKTGDSLAMARRLARSVRR LTRRRAHRLLRARRLLKREGVLQAADFDENGLIKSLPNTPWQLRAAALDRKLTPLEWSAVLLHLIKHRGYLSQRKNEGETADKELGALLKG VANNAHALQTGDFRTPAELALNKFEKESGHIRNQRGDYSHTFSRKDLQAELILLFEKQKEFGNPHVSGGLKEGIETLLMTQRPALSGDAV QKMLGHCTFEPAEPKAAKNTYTAERFIWLTKLNNLRILEQGSERPLTDTERATLMDEPYRKSKLTYAQARKLLGLEDTAFFKGLRYGKDNA EASTLMEMKAYHAISRALEKEGLKDKKSPLNLSSELQDEIGTAFSLFKTDEDITGRLKDRVQPEILEALLKHISFDKFVQISLKALRRIVPLMEQ GKRYDEACAEIYGDHYGKKNTEEKIYLPPIPADEIRNPVVLRALSQARKVINGVVRRYGSPARIHIETAREVGKSFKDRKEIEKRQEENRKDR EKAAAKFREYFPNFVGEPKSKDILKLRLYEQQHGKCLYSGKEINLVRLNEKGYVEIDHALPFSRTWDDSFNNKVLVLGSENQNKGNQTPYE YFNGKDNSREWQEFKARVETSRFPRSKKQRILLQKFDEDGFKECNLNDTRYVNRFLCQFVADHILLTGKGKRRVFASNGQITNLLRGFWG LRKVRAENDRHHALDAVVVACSTVAMQQKITRFVRYKEMNAFDGKTIDKETGKVLHQKTHFPQPWEFFAQEVMIRVFGKPDGKPEFEE ADTPEKLRTLLAEKLSSRPEAVHEYVTPLFVSRAPNRKMSGAHKDTLRSAKRFVKHNEKISVKRVWLTEIKLADLENMVNYKNGREIELYEA LKARLEAYGGNAKQAFDPKDNPFYKKGGQLVKAVRVEKTQRSGVLLNKKNAYTIADNGDMVRVDVFCKVDKKGKNQYFIVPIYAWQVA ENILPDIDCKGYRIDDSYTFCFSLHKYDLIAFQKDEKSKVEFAYYINCDSSNGRFYLAWHDKGSKEQQFRISTQNLVLIQKYQVNELGKEIRPC RLKKRPPVREDKRTADGSEFEPKKKRKV

Atgaaacggacagccgacggaagcgagttcgagtcaccaaagaagaagcggaaagtcgaagatatggccgccttcaagcctaacccaatcaattacatcctgggactgga catcggaatcgcatccgtgggatgggctatggtggagatcgacgaggaggagaatcctatccggctgatcgatctgggcgtgagagtgtttgagagggccgaggtgccaaag accggcgattctctggctatggcccggagactggcacggagcgtgaggcgcctgacacggagaagggcacacaggctgctgagggcacgccggctgctgaagagagaggg cgtgctgcaggcagcagacttcgatgagaatggcctgatcaagagcctgccaaacaccccctggcagctgagagcagccgccctggacaggaagctgacaccactggagtg gtctgccgtgctgctgcacctgatcaagcaccgcggctacctgagccagcggaagaacgagggagagacagcagacaaggagctgggcgccctgctgaagggagtggcca acaatgcccacgccctgcagaccggcgatttcaggacacctgccgagctggccctgaataagtttgagaaggagtccggccacatcagaaaccagaggggcgactatagcc acaccttctcccgcaaggatctgcaggccgagctgatcctgctgttcgagaagcagaaggagtttggcaatccacacgtgagcggaggcctgaaggagggaatcgagaccct gctgatgacacagaggcctgccctgtccggcgacgcagtgcagaagatgctgggacactgcaccttcgagcctgcagagccaaaggccgccaagaacacctacacagccga gcggtttatctggctgacaaagctgaacaatctgagaatcctggagcagggatccgagaggccactgaccgacacagagagggccaccctgatggatgagccttaccggaa gtctaagctgacatatgcccaggccagaaagctgctgggcctggaggacaccgccttctttaagggcctgagatacggcaaggataatgccgaggcctccacactgatggag atgaaggcctatcacgccatctctcgcgccctggagaaggagggcctgaaggacaagaagtcccccctgaacctgagctccgagctgcaggatgagatcggcaccgccttct ctctgtttaagaccgacgaggatatcacaggccgcctgaaggacagggtgcagcctgagatcctggaggccctgctgaagcacatctctttcgataagtttgtgcagatcagc ctgaaggccctgagaaggatcgtgccactgatggagcagggcaagcggtacgacgaggcctgcgccgagatctacggcgatcactatggcaagaagaacacagaggaga agatctatctgccccctatccctgccgacgagatcagaaatcctgtggtgctgagggccctgtcccaggcaagaaaagtgatcaacggagtggtgcgccggtacggatctcca gcccggatccacatcgagaccgccagagaagtgggcaagagcttcaaggaccggaaggagatcgagaagagacaggaggagaatcgcaaggatcgggagaaggccgcc gccaagtttagggagtacttccctaactttgtgggcgagccaaagtctaaggacatcctgaagctgcgcctgtacgagcagcagcacggcaagtgtctgtatagcggcaagg agatcaatctggtgcggctgaacgagaagggctatgtggagatcgatcacgccctgcctttctccagaacctgggacgattcttttaacaataaggtgctggtgctgggcagcg agaaccagaataagggcaatcagacaccatacgagtatttcaatggcaaggacaactccagggagtggcaggagttcaaggcccgcgtggagacctctagatttcccagga gcaagaagcagcggatcctgctgcagaagttcgacgaggatggctttaaggagtgcaacctgaatgacaccagatacgtgaaccggttcctgtgccagtttgtggccgatca catcctgctgaccggcaagggcaagagaagggtgttcgcctctaatggccagatcacaaacctgctgaggggattttggggactgaggaaggtgcgggcagagaatgacag acaccacgcactggatgcagtggtggtggcatgcagcaccgtggcaatgcagcagaagatcacaagattcgtgaggtataaggagatgaacgcctttgacggcaagaccat cgataaggagacaggcaaggtgctgcaccagaagacccacttcccccagccttgggagttctttgcccaggaagtgatgatccgggtgttcggcaagccagacggcaagcct gagtttgaggaggccgataccccagagaagctgaggacactgctggcagagaagctgtctagcaggccagaggcagtgcacgagtacgtgaccccactgttcgtgtccagg gcacccaatcggaagatgtctggcgcccacaaggacacactgagaagcgccaagaggtttgtgaagcacaacgagaagatctccgtgaagagagtgtggctgaccgagat caagctggccgatctggagaacatggtgaattacaagaacggcagggagatcgagctgtatgaggccctgaaggcaaggctggaggcctacggaggaaatgccaagcagg ccttcgacccaaaggataaccccttttataagaagggaggacagctggtgaaggccgtgcgggtggagaagacccagCGTagcggcgtgctgctgaataagaagaacgcc tacacaatcgccgacaatggcgatatggtgagagtggacgtgttctgtaaggtggataagaagggcaagaatcagtactttatcgtgcctatctatgcctggcaggtggccga

**Nme2Cas9 (D56R)**: <u>BPSV40-NLS</u>, Nme2Cas9, Linkers

MKRTADGSEFESPKKKRKVEDMAAFKPNPINYILGLDIGIASVGWAMVEIDEEENPIRLIDLGVRVFERAEVPKTGRSLAMARRLARSVRR LTRRRAHRLLRARRLLKREGVLQAADFDENGLIKSLPNTPWQLRAAALDRKLTPLEWSAVLLHLIKHRGYLSQRKNEGETADKELGALLKG VANNAHALQTGDFRTPAELALNKFEKESGHIRNQRGDYSHTFSRKDLQAELILLFEKQKEFGNPHVSGGLKEGIETLLMTQRPALSGDAV QKMLGHCTFEPAEPKAAKNTYTAERFIWLTKLNNLRILEQGSERPLTDTERATLMDEPYRKSKLTYAQARKLLGLEDTAFFKGLRYGKDNA EASTLMEMKAYHAISRALEKEGLKDKKSPLNLSSELQDEIGTAFSLFKTDEDITGRLKDRVQPEILEALLKHISFDKFVQISLKALRRIVPLMEQ GKRYDEACAEIYGDHYGKKNTEEKIYLPPIPADEIRNPVVLRALSQARKVINGVVRRYGSPARIHIETAREVGKSFKDRKEIEKRQEENRKDR EKAAAKFREYFPNFVGEPKSKDILKLRLYEQQHGKCLYSGKEINLVRLNEKGYVEIDHALPFSRTWDDSFNNKVLVLGSENQNKGNQTPYE YFNGKDNSREWQEFKARVETSRFPRSKKQRILLQKFDEDGFKECNLNDTRYVNRFLCQFVADHILLTGKGKRRVFASNGQITNLLRGFWG LRKVRAENDRHHALDAVVVACSTVAMQQKITRFVRYKEMNAFDGKTIDKETGKVLHQKTHFPQPWEFFAQEVMIRVFGKPDGKPEFEE ADTPEKLRTLLAEKLSSRPEAVHEYVTPLFVSRAPNRKMSGAHKDTLRSAKRFVKHNEKISVKRVWLTEIKLADLENMVNYKNGREIELYEA LKARLEAYGGNAKQAFDPKDNPFYKKGGQLVKAVRVEKTQESGVLLNKKNAYTIADNGDMVRVDVFCKVDKKGKNQYFIVPIYAWQVA ENILPDIDCKGYRIDDSYTFCFSLHKYDLIAFQKDEKSKVEFAYYINCDSSNGRFYLAWHDKGSKEQQFRISTQNLVLIQKYQVNELGKEIRPC RLKKRPPVREDKRTADGSEFEPKKKRKV

Atgaaacggacagccgacggaagcgagttcgagtcaccaaagaagaagcggaaagtcgaagatatggccgccttcaagcctaacccaatcaattacatcctgggactgga catcggaatcgcatccgtgggatgggctatggtggagatcgacgaggaggagaatcctatccggctgatcgatctgggcgtgagagtgtttgagagggccgaggtgccaaag accggcCGTtctctggctatggcccggagactggcacggagcgtgaggcgcctgacacggagaagggcacacaggctgctgagggcacgccggctgctgaagagagagg gcgtgctgcaggcagcagacttcgatgagaatggcctgatcaagagcctgccaaacaccccctggcagctgagagcagccgccctggacaggaagctgacaccactggagt ggtctgccgtgctgctgcacctgatcaagcaccgcggctacctgagccagcggaagaacgagggagagacagcagacaaggagctgggcgccctgctgaagggagtggcc aacaatgcccacgccctgcagaccggcgatttcaggacacctgccgagctggccctgaataagtttgagaaggagtccggccacatcagaaaccagaggggcgactatagc cacaccttctcccgcaaggatctgcaggccgagctgatcctgctgttcgagaagcagaaggagtttggcaatccacacgtgagcggaggcctgaaggagggaatcgagacc ctgctgatgacacagaggcctgccctgtccggcgacgcagtgcagaagatgctgggacactgcaccttcgagcctgcagagccaaaggccgccaagaacacctacacagcc gagcggtttatctggctgacaaagctgaacaatctgagaatcctggagcagggatccgagaggccactgaccgacacagagagggccaccctgatggatgagccttaccgg aagtctaagctgacatatgcccaggccagaaagctgctgggcctggaggacaccgccttctttaagggcctgagatacggcaaggataatgccgaggcctccacactgatgg agatgaaggcctatcacgccatctctcgcgccctggagaaggagggcctgaaggacaagaagtcccccctgaacctgagctccgagctgcaggatgagatcggcaccgcctt ctctctgtttaagaccgacgaggatatcacaggccgcctgaaggacagggtgcagcctgagatcctggaggccctgctgaagcacatctctttcgataagtttgtgcagatcag cctgaaggccctgagaaggatcgtgccactgatggagcagggcaagcggtacgacgaggcctgcgccgagatctacggcgatcactatggcaagaagaacacagaggag aagatctatctgccccctatccctgccgacgagatcagaaatcctgtggtgctgagggccctgtcccaggcaagaaaagtgatcaacggagtggtgcgccggtacggatctcc agcccggatccacatcgagaccgccagagaagtgggcaagagcttcaaggaccggaaggagatcgagaagagacaggaggagaatcgcaaggatcgggagaaggccgc cgccaagtttagggagtacttccctaactttgtgggcgagccaaagtctaaggacatcctgaagctgcgcctgtacgagcagcagcacggcaagtgtctgtatagcggcaagg agatcaatctggtgcggctgaacgagaagggctatgtggagatcgatcacgccctgcctttctccagaacctgggacgattcttttaacaataaggtgctggtgctgggcagcg agaaccagaataagggcaatcagacaccatacgagtatttcaatggcaaggacaactccagggagtggcaggagttcaaggcccgcgtggagacctctagatttcccagga gcaagaagcagcggatcctgctgcagaagttcgacgaggatggctttaaggagtgcaacctgaatgacaccagatacgtgaaccggttcctgtgccagtttgtggccgatca catcctgctgaccggcaagggcaagagaagggtgttcgcctctaatggccagatcacaaacctgctgaggggattttggggactgaggaaggtgcgggcagagaatgacag acaccacgcactggatgcagtggtggtggcatgcagcaccgtggcaatgcagcagaagatcacaagattcgtgaggtataaggagatgaacgcctttgacggcaagaccat cgataaggagacaggcaaggtgctgcaccagaagacccacttcccccagccttgggagttctttgcccaggaagtgatgatccgggtgttcggcaagccagacggcaagcct gagtttgaggaggccgataccccagagaagctgaggacactgctggcagagaagctgtctagcaggccagaggcagtgcacgagtacgtgaccccactgttcgtgtccagg gcacccaatcggaagatgtctggcgcccacaaggacacactgagaagcgccaagaggtttgtgaagcacaacgagaagatctccgtgaagagagtgtggctgaccgagat caagctggccgatctggagaacatggtgaattacaagaacggcagggagatcgagctgtatgaggccctgaaggcaaggctggaggcctacggaggaaatgccaagcagg ccttcgacccaaaggataaccccttttataagaagggaggacagctggtgaaggccgtgcgggtggagaagacccaggagagcggcgtgctgctgaataagaagaacgcc tacacaatcgccgacaatggcgatatggtgagagtggacgtgttctgtaaggtggataagaagggcaagaatcagtactttatcgtgcctatctatgcctggcaggtggccga gaacatcctgccagacatcgattgcaagggctacagaatcgacgatagctatacattctgtttttccctgcacaagtatgacctgatcgccttccagaaggatgagaagtccaa ggtggagtttgcctactatatcaattgcgactcctctaacggcaggttctacctggcctggcacgataagggcagcaaggagcagcagtttcgcatctccacccagaatctggt gctgatccagaagtatcaggtgaacgagctgggcaaggagatcaggccatgtcggctgaagaagcgcccacccgtgcgggaggataaaagaaccgccgacggcagcgaa ttcgagcccaagaagaagaggaaagtc

**Nme2^Smu^Cas9 (WT)**: <u>BPSV40-NLS</u>, Nme2Cas9 – delta PID, <u>SmuCas9 PID</u>, Linkers

MKRTADGSEFESPKKKRKVEDMAAFKPNPINYILGLDIGIASVGWAMVEIDEEENPIRLIDLGVRVFERAEVPKTGDSLAMARRLARSVRR LTRRRAHRLLRARRLLKREGVLQAADFDENGLIKSLPNTPWQLRAAALDRKLTPLEWSAVLLHLIKHRGYLSQRKNEGETADKELGALLKG VANNAHALQTGDFRTPAELALNKFEKESGHIRNQRGDYSHTFSRKDLQAELILLFEKQKEFGNPHVSGGLKEGIETLLMTQRPALSGDAV QKMLGHCTFEPAEPKAAKNTYTAERFIWLTKLNNLRILEQGSERPLTDTERATLMDEPYRKSKLTYAQARKLLGLEDTAFFKGLRYGKDNA EASTLMEMKAYHAISRALEKEGLKDKKSPLNLSSELQDEIGTAFSLFKTDEDITGRLKDRVQPEILEALLKHISFDKFVQISLKALRRIVPLMEQ GKRYDEACAEIYGDHYGKKNTEEKIYLPPIPADEIRNPVVLRALSQARKVINGVVRRYGSPARIHIETAREVGKSFKDRKEIEKRQEENRKDR EKAAAKFREYFPNFVGEPKSKDILKLRLYEQQHGKCLYSGKEINLVRLNEKGYVEIDHALPFSRTWDDSFNNKVLVLGSENQNKGNQTPYE YFNGKDNSREWQEFKARVETSRFPRSKKQRILLQKFDEDGFKECNLNDTRYVNRFLCQFVADHILLTGKGKRRVFASNGQITNLLRGFWG LRKVRAENDRHHALDAVVVACSTVAMQQKITRFVRYKEMNAFDGKTIDKETGKVLHQKTHFPQPWEFFAQEVMIRVFGKPDGKPEFEE ADTPEKLRTLLAEKLSSRPEAVHEYVTPLFVSRAPNRKMSGAHKDTLRSAKRFVKHNEKISVKRVWLTEIKLADLENMVNYKNGREIELYEA LKARLEAYGGNAKQAFDPKDNPFYKKGGQLVKAVRVEKTQESGVLLNKKNAYTIADNATMVRVDVYTKAGKNYLVPVYVWQVAQGILP NRAVTSGKSEADWDLIDESFEFKFSLSRGDLVEMISNKGRIFGYYNGLDRANGSIGIREHDLEKSKGKDGVHRVGVKTATAFNKYHVDPLG KEIHRCSSEPRPTLKIKSKKEDKRTADGSEFEPKKKRKV

atgaaacggacagccgacggaagcgagttcgagtcaccaaagaagaagcggaaagtcgaagatatggccgccttcaagcctaacccaatcaattacatcctgggactgga catcggaatcgcatccgtgggatgggctatggtggagatcgacgaggaggagaatcctatccggctgatcgatctgggcgtgagagtgtttgagagggccgaggtgccaaag accggcgattctctggctatggcccggagactggcacggagcgtgaggcgcctgacacggagaagggcacacaggctgctgagggcacgccggctgctgaagagagaggg cgtgctgcaggcagcagacttcgatgagaatggcctgatcaagagcctgccaaacaccccctggcagctgagagcagccgccctggacaggaagctgacaccactggagtg gtctgccgtgctgctgcacctgatcaagcaccgcggctacctgagccagcggaagaacgagggagagacagcagacaaggagctgggcgccctgctgaagggagtggcca acaatgcccacgccctgcagaccggcgatttcaggacacctgccgagctggccctgaataagtttgagaaggagtccggccacatcagaaaccagaggggcgactatagcc acaccttctcccgcaaggatctgcaggccgagctgatcctgctgttcgagaagcagaaggagtttggcaatccacacgtgagcggaggcctgaaggagggaatcgagaccct gctgatgacacagaggcctgccctgtccggcgacgcagtgcagaagatgctgggacactgcaccttcgagcctgcagagccaaaggccgccaagaacacctacacagccga gcggtttatctggctgacaaagctgaacaatctgagaatcctggagcagggatccgagaggccactgaccgacacagagagggccaccctgatggatgagccttaccggaa gtctaagctgacatatgcccaggccagaaagctgctgggcctggaggacaccgccttctttaagggcctgagatacggcaaggataatgccgaggcctccacactgatggag atgaaggcctatcacgccatctctcgcgccctggagaaggagggcctgaaggacaagaagtcccccctgaacctgagctccgagctgcaggatgagatcggcaccgccttct ctctgtttaagaccgacgaggatatcacaggccgcctgaaggacagggtgcagcctgagatcctggaggccctgctgaagcacatctctttcgataagtttgtgcagatcagc ctgaaggccctgagaaggatcgtgccactgatggagcagggcaagcggtacgacgaggcctgcgccgagatctacggcgatcactatggcaagaagaacacagaggaga agatctatctgccccctatccctgccgacgagatcagaaatcctgtggtgctgagggccctgtcccaggcaagaaaagtgatcaacggagtggtgcgccggtacggatctcca gcccggatccacatcgagaccgccagagaagtgggcaagagcttcaaggaccggaaggagatcgagaagagacaggaggagaatcgcaaggatcgggagaaggccgcc gccaagtttagggagtacttccctaactttgtgggcgagccaaagtctaaggacatcctgaagctgcgcctgtacgagcagcagcacggcaagtgtctgtatagcggcaagg agatcaatctggtgcggctgaacgagaagggctatgtggagatcgatcacgccctgcctttctccagaacctgggacgattcttttaacaataaggtgctggtgctgggcagcg agaaccagaataagggcaatcagacaccatacgagtatttcaatggcaaggacaactccagggagtggcaggagttcaaggcccgcgtggagacctctagatttcccagga gcaagaagcagcggatcctgctgcagaagttcgacgaggatggctttaaggagtgcaacctgaatgacaccagatacgtgaaccggttcctgtgccagtttgtggccgatca catcctgctgaccggcaagggcaagagaagggtgttcgcctctaatggccagatcacaaacctgctgaggggattttggggactgaggaaggtgcgggcagagaatgacag acaccacgcactggatgcagtggtggtggcatgcagcaccgtggcaatgcagcagaagatcacaagattcgtgaggtataaggagatgaacgcctttgacggcaagaccat cgataaggagacaggcaaggtgctgcaccagaagacccacttcccccagccttgggagttctttgcccaggaagtgatgatccgggtgttcggcaagccagacggcaagcct gagtttgaggaggccgataccccagagaagctgaggacactgctggcagagaagctgtctagcaggccagaggcagtgcacgagtacgtgaccccactgttcgtgtccagg gcacccaatcggaagatgtctggcgcccacaaggacacactgagaagcgccaagaggtttgtgaagcacaacgagaagatctccgtgaagagagtgtggctgaccgagat caagctggccgatctggagaacatggtgaattacaagaacggcagggagatcgagctgtatgaggccctgaaggcaaggctggaggcctacggaggaaatgccaagcagg ccttcgacccaaaggataaccccttttataagaagggaggacagctggtgaaggccgtgcgggtggagaagacccaggagagcggcgtgctgctgaataagaagaacgcc tacacaatcgccgacaacgccaccatggtgcgggtggacgtgtacaccaaggccggcaagaactacctggttcctgtgtacgtgtggcaggtggcccagggcatcttaccca accgcgccgtgaccagcggcaagtccgaggctgactgggacctgatcgatgagagcttcgagttcaagttctctctgtcccggggagatctcgtggaaatgatctccaacaag ggcagaatcttcggctactacaacggcctggacagagccaacggctctattggaattagagagcacgacctagagaagagcaagggcaaagacggcgtgcatagagtggg agtgaaaacagctacagcatttaacaagtaccacgtggatcccctgggcaaagagatccacagatgcagcagcgaacccagacctacactgaaaatcaagtctaagaagg aggataaaagaaccgccgacggcagcgaattcgagcccaagaagaagaggaaagtc

**Nme2^Smu^Cas9 (E932R)**: <u>BPSV40-NLS</u>, Nme2Cas9 – delta PID, <u>SmuCas9 PID</u>, Linkers

MKRTADGSEFESPKKKRKVEDMAAFKPNPINYILGLDIGIASVGWAMVEIDEEENPIRLIDLGVRVFERAEVPKTGDSLAMARRLARSVRR LTRRRAHRLLRARRLLKREGVLQAADFDENGLIKSLPNTPWQLRAAALDRKLTPLEWSAVLLHLIKHRGYLSQRKNEGETADKELGALLKG VANNAHALQTGDFRTPAELALNKFEKESGHIRNQRGDYSHTFSRKDLQAELILLFEKQKEFGNPHVSGGLKEGIETLLMTQRPALSGDAV QKMLGHCTFEPAEPKAAKNTYTAERFIWLTKLNNLRILEQGSERPLTDTERATLMDEPYRKSKLTYAQARKLLGLEDTAFFKGLRYGKDNA EASTLMEMKAYHAISRALEKEGLKDKKSPLNLSSELQDEIGTAFSLFKTDEDITGRLKDRVQPEILEALLKHISFDKFVQISLKALRRIVPLMEQ GKRYDEACAEIYGDHYGKKNTEEKIYLPPIPADEIRNPVVLRALSQARKVINGVVRRYGSPARIHIETAREVGKSFKDRKEIEKRQEENRKDR EKAAAKFREYFPNFVGEPKSKDILKLRLYEQQHGKCLYSGKEINLVRLNEKGYVEIDHALPFSRTWDDSFNNKVLVLGSENQNKGNQTPYE YFNGKDNSREWQEFKARVETSRFPRSKKQRILLQKFDEDGFKECNLNDTRYVNRFLCQFVADHILLTGKGKRRVFASNGQITNLLRGFWG LRKVRAENDRHHALDAVVVACSTVAMQQKITRFVRYKEMNAFDGKTIDKETGKVLHQKTHFPQPWEFFAQEVMIRVFGKPDGKPEFEE ADTPEKLRTLLAEKLSSRPEAVHEYVTPLFVSRAPNRKMSGAHKDTLRSAKRFVKHNEKISVKRVWLTEIKLADLENMVNYKNGREIELYEA LKARLEAYGGNAKQAFDPKDNPFYKKGGQLVKAVRVEKTQRSGVLLNKKNAYTIADNATMVRVDVYTKAGKNYLVPVYVWQVAQGILP NRAVTSGKSEADWDLIDESFEFKFSLSRGDLVEMISNKGRIFGYYNGLDRANGSIGIREHDLEKSKGKDGVHRVGVKTATAFNKYHVDPLG KEIHRCSSEPRPTLKIKSKKEDKRTADGSEFEPKKKRKV

atgaaacggacagccgacggaagcgagttcgagtcaccaaagaagaagcggaaagtcgaagatatggccgccttcaagcctaacccaatcaattacatcctgggactgga catcggaatcgcatccgtgggatgggctatggtggagatcgacgaggaggagaatcctatccggctgatcgatctgggcgtgagagtgtttgagagggccgaggtgccaaag accggcgattctctggctatggcccggagactggcacggagcgtgaggcgcctgacacggagaagggcacacaggctgctgagggcacgccggctgctgaagagagaggg cgtgctgcaggcagcagacttcgatgagaatggcctgatcaagagcctgccaaacaccccctggcagctgagagcagccgccctggacaggaagctgacaccactggagtg gtctgccgtgctgctgcacctgatcaagcaccgcggctacctgagccagcggaagaacgagggagagacagcagacaaggagctgggcgccctgctgaagggagtggcca acaatgcccacgccctgcagaccggcgatttcaggacacctgccgagctggccctgaataagtttgagaaggagtccggccacatcagaaaccagaggggcgactatagcc acaccttctcccgcaaggatctgcaggccgagctgatcctgctgttcgagaagcagaaggagtttggcaatccacacgtgagcggaggcctgaaggagggaatcgagaccct gctgatgacacagaggcctgccctgtccggcgacgcagtgcagaagatgctgggacactgcaccttcgagcctgcagagccaaaggccgccaagaacacctacacagccga gcggtttatctggctgacaaagctgaacaatctgagaatcctggagcagggatccgagaggccactgaccgacacagagagggccaccctgatggatgagccttaccggaa gtctaagctgacatatgcccaggccagaaagctgctgggcctggaggacaccgccttctttaagggcctgagatacggcaaggataatgccgaggcctccacactgatggag atgaaggcctatcacgccatctctcgcgccctggagaaggagggcctgaaggacaagaagtcccccctgaacctgagctccgagctgcaggatgagatcggcaccgccttct ctctgtttaagaccgacgaggatatcacaggccgcctgaaggacagggtgcagcctgagatcctggaggccctgctgaagcacatctctttcgataagtttgtgcagatcagc ctgaaggccctgagaaggatcgtgccactgatggagcagggcaagcggtacgacgaggcctgcgccgagatctacggcgatcactatggcaagaagaacacagaggaga agatctatctgccccctatccctgccgacgagatcagaaatcctgtggtgctgagggccctgtcccaggcaagaaaagtgatcaacggagtggtgcgccggtacggatctcca gcccggatccacatcgagaccgccagagaagtgggcaagagcttcaaggaccggaaggagatcgagaagagacaggaggagaatcgcaaggatcgggagaaggccgcc gccaagtttagggagtacttccctaactttgtgggcgagccaaagtctaaggacatcctgaagctgcgcctgtacgagcagcagcacggcaagtgtctgtatagcggcaagg agatcaatctggtgcggctgaacgagaagggctatgtggagatcgatcacgccctgcctttctccagaacctgggacgattcttttaacaataaggtgctggtgctgggcagcg agaaccagaataagggcaatcagacaccatacgagtatttcaatggcaaggacaactccagggagtggcaggagttcaaggcccgcgtggagacctctagatttcccagga gcaagaagcagcggatcctgctgcagaagttcgacgaggatggctttaaggagtgcaacctgaatgacaccagatacgtgaaccggttcctgtgccagtttgtggccgatca catcctgctgaccggcaagggcaagagaagggtgttcgcctctaatggccagatcacaaacctgctgaggggattttggggactgaggaaggtgcgggcagagaatgacag acaccacgcactggatgcagtggtggtggcatgcagcaccgtggcaatgcagcagaagatcacaagattcgtgaggtataaggagatgaacgcctttgacggcaagaccat cgataaggagacaggcaaggtgctgcaccagaagacccacttcccccagccttgggagttctttgcccaggaagtgatgatccgggtgttcggcaagccagacggcaagcct gagtttgaggaggccgataccccagagaagctgaggacactgctggcagagaagctgtctagcaggccagaggcagtgcacgagtacgtgaccccactgttcgtgtccagg gcacccaatcggaagatgtctggcgcccacaaggacacactgagaagcgccaagaggtttgtgaagcacaacgagaagatctccgtgaagagagtgtggctgaccgagat caagctggccgatctggagaacatggtgaattacaagaacggcagggagatcgagctgtatgaggccctgaaggcaaggctggaggcctacggaggaaatgccaagcagg ccttcgacccaaaggataaccccttttataagaagggaggacagctggtgaaggccgtgcgggtggagaagacccagCGTagcggcgtgctgctgaataagaagaacgcc tacacaatcgccgacaacgccaccatggtgcgggtggacgtgtacaccaaggccggcaagaactacctggttcctgtgtacgtgtggcaggtggcccagggcatcttaccca accgcgccgtgaccagcggcaagtccgaggctgactgggacctgatcgatgagagcttcgagttcaagttctctctgtcccggggagatctcgtggaaatgatctccaacaag ggcagaatcttcggctactacaacggcctggacagagccaacggctctattggaattagagagcacgacctagagaagagcaagggcaaagacggcgtgcatagagtggg agtgaaaacagctacagcatttaacaagtaccacgtggatcccctgggcaaagagatccacagatgcagcagcgaacccagacctacactgaaaatcaagtctaagaagg aggataaaagaaccgccgacggcagcgaattcgagcccaagaagaagaggaaagtc

**Nme2^Smu^Cas9 (D56R)**: <u>BPSV40-NLS</u>, Nme2Cas9 – delta PID, <u>SmuCas9 PID</u>, Linkers

MKRTADGSEFESPKKKRKVEDMAAFKPNPINYILGLDIGIASVGWAMVEIDEEENPIRLIDLGVRVFERAEVPKTGRSLAMARRLARSVRR LTRRRAHRLLRARRLLKREGVLQAADFDENGLIKSLPNTPWQLRAAALDRKLTPLEWSAVLLHLIKHRGYLSQRKNEGETADKELGALLKG VANNAHALQTGDFRTPAELALNKFEKESGHIRNQRGDYSHTFSRKDLQAELILLFEKQKEFGNPHVSGGLKEGIETLLMTQRPALSGDAV QKMLGHCTFEPAEPKAAKNTYTAERFIWLTKLNNLRILEQGSERPLTDTERATLMDEPYRKSKLTYAQARKLLGLEDTAFFKGLRYGKDNA EASTLMEMKAYHAISRALEKEGLKDKKSPLNLSSELQDEIGTAFSLFKTDEDITGRLKDRVQPEILEALLKHISFDKFVQISLKALRRIVPLMEQ GKRYDEACAEIYGDHYGKKNTEEKIYLPPIPADEIRNPVVLRALSQARKVINGVVRRYGSPARIHIETAREVGKSFKDRKEIEKRQEENRKDR EKAAAKFREYFPNFVGEPKSKDILKLRLYEQQHGKCLYSGKEINLVRLNEKGYVEIDHALPFSRTWDDSFNNKVLVLGSENQNKGNQTPYE YFNGKDNSREWQEFKARVETSRFPRSKKQRILLQKFDEDGFKECNLNDTRYVNRFLCQFVADHILLTGKGKRRVFASNGQITNLLRGFWG LRKVRAENDRHHALDAVVVACSTVAMQQKITRFVRYKEMNAFDGKTIDKETGKVLHQKTHFPQPWEFFAQEVMIRVFGKPDGKPEFEE ADTPEKLRTLLAEKLSSRPEAVHEYVTPLFVSRAPNRKMSGAHKDTLRSAKRFVKHNEKISVKRVWLTEIKLADLENMVNYKNGREIELYEA LKARLEAYGGNAKQAFDPKDNPFYKKGGQLVKAVRVEKTQESGVLLNKKNAYTIADNATMVRVDVYTKAGKNYLVPVYVWQVAQGILP NRAVTSGKSEADWDLIDESFEFKFSLSRGDLVEMISNKGRIFGYYNGLDRANGSIGIREHDLEKSKGKDGVHRVGVKTATAFNKYHVDPLG KEIHRCSSEPRPTLKIKSKKEDKRTADGSEFEPKKKRKV

atgaaacggacagccgacggaagcgagttcgagtcaccaaagaagaagcggaaagtcgaagatatggccgccttcaagcctaacccaatcaattacatcctgggactgga catcggaatcgcatccgtgggatgggctatggtggagatcgacgaggaggagaatcctatccggctgatcgatctgggcgtgagagtgtttgagagggccgaggtgccaaag accggcCGTtctctggctatggcccggagactggcacggagcgtgaggcgcctgacacggagaagggcacacaggctgctgagggcacgccggctgctgaagagagagg gcgtgctgcaggcagcagacttcgatgagaatggcctgatcaagagcctgccaaacaccccctggcagctgagagcagccgccctggacaggaagctgacaccactggagt ggtctgccgtgctgctgcacctgatcaagcaccgcggctacctgagccagcggaagaacgagggagagacagcagacaaggagctgggcgccctgctgaagggagtggcc aacaatgcccacgccctgcagaccggcgatttcaggacacctgccgagctggccctgaataagtttgagaaggagtccggccacatcagaaaccagaggggcgactatagc cacaccttctcccgcaaggatctgcaggccgagctgatcctgctgttcgagaagcagaaggagtttggcaatccacacgtgagcggaggcctgaaggagggaatcgagacc ctgctgatgacacagaggcctgccctgtccggcgacgcagtgcagaagatgctgggacactgcaccttcgagcctgcagagccaaaggccgccaagaacacctacacagcc gagcggtttatctggctgacaaagctgaacaatctgagaatcctggagcagggatccgagaggccactgaccgacacagagagggccaccctgatggatgagccttaccgg aagtctaagctgacatatgcccaggccagaaagctgctgggcctggaggacaccgccttctttaagggcctgagatacggcaaggataatgccgaggcctccacactgatgg agatgaaggcctatcacgccatctctcgcgccctggagaaggagggcctgaaggacaagaagtcccccctgaacctgagctccgagctgcaggatgagatcggcaccgcctt ctctctgtttaagaccgacgaggatatcacaggccgcctgaaggacagggtgcagcctgagatcctggaggccctgctgaagcacatctctttcgataagtttgtgcagatcag cctgaaggccctgagaaggatcgtgccactgatggagcagggcaagcggtacgacgaggcctgcgccgagatctacggcgatcactatggcaagaagaacacagaggag aagatctatctgccccctatccctgccgacgagatcagaaatcctgtggtgctgagggccctgtcccaggcaagaaaagtgatcaacggagtggtgcgccggtacggatctcc agcccggatccacatcgagaccgccagagaagtgggcaagagcttcaaggaccggaaggagatcgagaagagacaggaggagaatcgcaaggatcgggagaaggccgc cgccaagtttagggagtacttccctaactttgtgggcgagccaaagtctaaggacatcctgaagctgcgcctgtacgagcagcagcacggcaagtgtctgtatagcggcaagg agatcaatctggtgcggctgaacgagaagggctatgtggagatcgatcacgccctgcctttctccagaacctgggacgattcttttaacaataaggtgctggtgctgggcagcg agaaccagaataagggcaatcagacaccatacgagtatttcaatggcaaggacaactccagggagtggcaggagttcaaggcccgcgtggagacctctagatttcccagga gcaagaagcagcggatcctgctgcagaagttcgacgaggatggctttaaggagtgcaacctgaatgacaccagatacgtgaaccggttcctgtgccagtttgtggccgatca catcctgctgaccggcaagggcaagagaagggtgttcgcctctaatggccagatcacaaacctgctgaggggattttggggactgaggaaggtgcgggcagagaatgacag acaccacgcactggatgcagtggtggtggcatgcagcaccgtggcaatgcagcagaagatcacaagattcgtgaggtataaggagatgaacgcctttgacggcaagaccat cgataaggagacaggcaaggtgctgcaccagaagacccacttcccccagccttgggagttctttgcccaggaagtgatgatccgggtgttcggcaagccagacggcaagcct gagtttgaggaggccgataccccagagaagctgaggacactgctggcagagaagctgtctagcaggccagaggcagtgcacgagtacgtgaccccactgttcgtgtccagg gcacccaatcggaagatgtctggcgcccacaaggacacactgagaagcgccaagaggtttgtgaagcacaacgagaagatctccgtgaagagagtgtggctgaccgagat caagctggccgatctggagaacatggtgaattacaagaacggcagggagatcgagctgtatgaggccctgaaggcaaggctggaggcctacggaggaaatgccaagcagg ccttcgacccaaaggataaccccttttataagaagggaggacagctggtgaaggccgtgcgggtggagaagacccaggagagcggcgtgctgctgaataagaagaacgcc tacacaatcgccgacaacgccaccatggtgcgggtggacgtgtacaccaaggccggcaagaactacctggttcctgtgtacgtgtggcaggtggcccagggcatcttaccca accgcgccgtgaccagcggcaagtccgaggctgactgggacctgatcgatgagagcttcgagttcaagttctctctgtcccggggagatctcgtggaaatgatctccaacaag ggcagaatcttcggctactacaacggcctggacagagccaacggctctattggaattagagagcacgacctagagaagagcaagggcaaagacggcgtgcatagagtggg agtgaaaacagctacagcatttaacaagtaccacgtggatcccctgggcaaagagatccacagatgcagcagcgaacccagacctacactgaaaatcaagtctaagaagg aggataaaagaaccgccgacggcagcgaattcgagcccaagaagaagaggaaagtc

**eNme2-C.NR (vLiu)**: <u>BPSV40-NLS</u>, eNme2-C.NR, Linkers

David Liu Lab evolved Nme2Cas9 nuclease for N4CN PAM targeting without alterations

MKRTADGSEFESPKKKRKVAAFKPNPINYILGLDIGIASVGWAMVEIDEEENPIRLIDLGVRVFERAEVPKTGDSLAMARRLARSVRRLTRRRAHRLLRARRLLKREGVLQAADFDENGLITSLPNTPWQLRAAALDRKLTPLEWSAVLLHLIKHRGYLSQRKNEGETAAKELGALLKGVANNAHALQTGDFRTPAELALNKFEKESGHIRNQRGDYSHTFSRKDLQAELILLFEKQKEFGNPHVSGGLKEGIETLLMTQRPALSGDAVQKMLGHCTLEPTEPKAAKNTYTAERFIWLTKLNNLRILEQGSERPLTDTERSTLMDEPYRKSKLTYAQARKLLGLEDTAFFKGLRYGKDNAEASTLMEMKAYHAISRALEKEGLKDKKSPLNLSSELQDEIGTAFSLFKTDEDITGRLKDRVQPEILEALLKHISFDKFVQISLKALRRIVPLMEQGKRYDEACAEIYGVHYGKKNTEEKIYLPPIPADEIRNPVVLRALSQARKVINGVVRRYGSPARIHIETAREVGKSFKDRKEIEKRQEENRKDREKAAAKFREYFPNFVGEPKSKDILKLRLYEQQHGKCLYSGKEINLVRLNEKGYVEIDHALPFSRTWDDSFNNKVLVLGSENQNKGNQTPYEYFNGKDNSREWQEFKARVETSRFPRSKKQRILLQKFDEDGFKECNLNDTRYVNRFLCQFVADHILLTGKGKRRVFASNGQITNLLRGFWGLRKVRAENDRHHALDAVVVACSTVAMQQKITRFVRYKEMNAFDGKTIDKETGKVLHQKTHFPQPWEFFAQEVMIRVFGKPDGKPEFEEADTPEKLRTLLAEKLSSRPEAVHEYVTPLFVSRAPNRKMSGAHKDTLRSAKRFVKHNEKISVKRVWLTEIKLADLENMVNYKNGREIELYEALKARLEAYGGNAKQAFDPKDNPFYKKGGQLVKAVRVEKTQKSGVLLNKKNAYTIADNGDMVRVDVFCKVDKKGKNQYFIVPIYAWQVAENILPDIDCKGYRIDDSYTFCFSLHKYDLIAFQKDEKSKVEFAYYINCDSSSGGFYLAWHDKGSREQRFRISTQNLALIQKYQVNELGKEIRPCRLKKRPPVRSGGSKRTADGSEFEPKKKRKV

atgaaacggacagccgacggaagcgagttcgagtcaccaaagaagaagcggaaagtcacagcattcaagcctaacccaatcaattacatcctgggactggatatcggaatcgcatccgtgggatgggctatggtggagatcgacgaggaggagaatcctatccggctgatcgatctgggcgtgagagtgtttgagagggccgaggtgccaaagaccggcgattctctggctatggcccggagactggcacggagcgtgaggcgcctgacacggagaagggcacacaggctgctgagggcacgccggctgctgaagagagagggcgtgctgcaggcagcagacttcgatgagaatggcctgatcacgagcttgccaaacaccccctggcagctgagagcagccgccctggacaggaagctgacaccactggagtggtctgccgtgctgctgcacctgatcaagcaccgcggctacctgagccagcggaagaacgagggagagacagcagccaaggagctgggcgccctgctgaagggagtggccaacaatgcccacgccctgcagaccggcgatttcaggacacctgccgagctggccctgaataagtttgagaaggagtccggccacatcagaaaccagaggggcgactatagccacaccttctcccgcaaggatctgcaggccgagctgatcctgctgttcgagaagcagaaggagtttggcaatccacacgtgagcggaggcctgaaggagggaatcgagaccctgctgatgacacagaggcctgccctgtccggcgacgcagtgcagaagatgctggggcactgcaccctcgagcctacagagccaaaggccgccaagaacacctacacagccgagcggtttatctggctgacaaagctgaacaatctgagaatcctggagcagggatccgagaggccactgaccgacacagagaggtccaccctgatggatgagccttaccggaagtctaaactgacatatgcccaggccagaaagctgctgggcctggaggacaccgccttctttaagggcctgagatacggcaaggataatgccgaggcctccacactgatggagatgaaggcctatcacgccatctctcgcgccctggagaaggagggcctgaaggacaagaagtcccccctgaacctgagctccgagctgcaggatgagatcggcaccgccttctctctgtttaagaccgacgaggatatcacaggccgcctgaaggacagggtgcagcctgagatcctggaggccctgctgaagcacatctctttcgataagtttgtgcagatcagcctgaaggccctgagaaggatcgtgccactgatggagcagggcaagcggtacgacgaggcctgcgccgagatctacggcgttcactatggcaagaagaacacagaggagaagatctatctgccccctatccctgccgacgagatcagaaatcctgtggtgctgagggccctgtcccaggcaagaaaagtgatcaacggagtggtgcgccggtacggatctccagcccggatccacatcgagaccgccagagaagtgggcaagagcttcaaggaccggaaggagatcgagaagagacaggaggagaatcgcaaggatcgggagaaggccgccgccaagtttagggagtacttccctaactttgtgggcgagccaaagtctaaggacatcctgaagctgcgcctgtacgagcagcagcacggcaagtgtctgtatagcggcaaggagatcaatctggtgcggctgaacgagaagggctatgtggagatcgatcacgccctgcctttctccagaacctgggacgattcttttaacaataaggtgctggtgctgggcagcgagaaccagaataagggcaatcagacaccatacgagtatttcaatggcaaggacaactccagggagtggcaggagttcaaggcccgcgtggagacctctagatttcccaggagcaagaagcagcggatcctgctgcagaagttcgacgaggatggctttaaggagtgcaacctgaatgacaccagatacgtgaaccggttcctgtgccagtttgtggccgatcacatcctgctgaccggcaagggcaagagaagggtgttcgcctctaatggccagatcacaaacctgctgaggggattttggggactgaggaaggtgcgggcagagaatgacagacaccacgcactggatgcagtggtggtggcatgcagcaccgtggcaatgcagcagaagatcacaagattcgtgaggtataaggagatgaacgcctttgacggcaagaccatcgataaggagacaggcaaggtgctgcaccagaagacccacttcccccagccttgggagttctttgcccaggaagtgatgatccgggtgttcggcaagccagacggcaagcctgagtttgaggaggccgataccccagagaagctgaggacactgctggcagagaagctgtctagcaggccagaggcagtgcacgagtacgtgaccccactgttcgtgtccagggcacccaatcggaagatgtctggcgcccacaaggacacactgagaagcgccaagaggtttgtgaagcacaacgagaagatctccgtgaagagagtgtggctgaccgagatcaagctggccgatctggagaacatggtgaattacaagaacggcagggagatcgagctgtatgaggccctgaaggcaaggctggaggcctacggaggaaatgccaagcaggccttcgacccaaaggataaccccttttataagaagggaggacagctggtgaaggccgtgcgggtggagaagacccagaagagcggcgtgctgctgaataagaagaacgcctacacaatcgccgacaatggtgatatggtgagagtggacgtgttctgtaaggtggataagaagggcaagaatcagtactttatcgtgcctatctatgcctggcaggtggccgagaacatcct gccagacatcgattgcaagggctacagaatcgacgatagctatacattctgtttttccctgcacaagtatgacctgatcgccttccagaaggatgagaagtccaaggtggagtt tgcctactatatcaattgcgactcctctagcggcgggttctacctggcctggcacgataagggcagcagggagcagcggtttcgcatctccacccagaatctggcgctgatcca gaagtatcaggtgaacgagctgggcaaggagatcaggccatgtcggctgaagaagcgcccacccgtgcggtctggcggctcaaaaagaaccgccgacggcagcgaattcg agcccaagaagaagaggaaagtc

**eNme2-C.NR (vEJS)**: <u>BPSV40-NLS</u>, eNme2-C.NR, Linkers

David Liu Lab evolved Nme2Cas9 nuclease for N4CN PAM targeting with linker and nuclear localization signals in the same framework as Nme2- and Nme2^Smu^Cas9 nucleases described in this work.

MKRTADGSEFESPKKKRKVEDAAFKPNPINYILGLDIGIASVGWAMVEIDEEENPIRLIDLGVRVFERAEVPKTGDSLAMARRLARSVRRLT RRRAHRLLRARRLLKREGVLQAADFDENGLITSLPNTPWQLRAAALDRKLTPLEWSAVLLHLIKHRGYLSQRKNEGETAAKELGALLKGVA NNAHALQTGDFRTPAELALNKFEKESGHIRNQRGDYSHTFSRKDLQAELILLFEKQKEFGNPHVSGGLKEGIETLLMTQRPALSGDAVQK MLGHCTLEPTEPKAAKNTYTAERFIWLTKLNNLRILEQGSERPLTDTERSTLMDEPYRKSKLTYAQARKLLGLEDTAFFKGLRYGKDNAEAS TLMEMKAYHAISRALEKEGLKDKKSPLNLSSELQDEIGTAFSLFKTDEDITGRLKDRVQPEILEALLKHISFDKFVQISLKALRRIVPLMEQGKR YDEACAEIYGVHYGKKNTEEKIYLPPIPADEIRNPVVLRALSQARKVINGVVRRYGSPARIHIETAREVGKSFKDRKEIEKRQEENRKDREKA AAKFREYFPNFVGEPKSKDILKLRLYEQQHGKCLYSGKEINLVRLNEKGYVEIDHALPFSRTWDDSFNNKVLVLGSENQNKGNQTPYEYFNGKDNSREWQEFKARVETSRFPRSKKQRILLQKFDEDGFKECNLNDTRYVNRFLCQFVADHILLTGKGKRRVFASNGQITNLLRGFWGLRKVRAENDRHHALDAVVVACSTVAMQQKITRFVRYKEMNAFDGKTIDKETGKVLHQKTHFPQPWEFFAQEVMIRVFGKPDGKPEFEEADTPEKLRTLLAEKLSSRPEAVHEYVTPLFVSRAPNRKMSGAHKDTLRSAKRFVKHNEKISVKRVWLTEIKLADLENMVNYKNGREIELYEALKARLEAYGGNAKQAFDPKDNPFYKKGGQLVKAVRVEKTQKSGVLLNKKNAYTIADNGDMVRVDVFCKVDKKGKNQYFIVPIYAWQVAENILPDIDCKGYRIDDSYTFCFSLHKYDLIAFQKDEKSKVEFAYYINCDSSSGGFYLAWHDKGSREQRFRISTQNLALIQKYQVNELGKEIRPCRLKKRPPVREDKRTADGSEFEPKKKRKV

atgaaacggacagccgacggaagcgagttcgagtcaccaaagaagaagcggaaagtcgaagatacagcattcaagcctaacccaatcaattacatcctgggactggatatcggaatcgcatccgtgggatgggctatggtggagatcgacgaggaggagaatcctatccggctgatcgatctgggcgtgagagtgtttgagagggccgaggtgccaaagaccggcgattctctggctatggcccggagactggcacggagcgtgaggcgcctgacacggagaagggcacacaggctgctgagggcacgccggctgctgaagagagagggcgtgctgcaggcagcagacttcgatgagaatggcctgatcacgagcttgccaaacaccccctggcagctgagagcagccgccctggacaggaagctgacaccactggagtggtctgccgtgctgctgcacctgatcaagcaccgcggctacctgagccagcggaagaacgagggagagacagcagccaaggagctgggcgccctgctgaagggagtggccaacaatgcccacgccctgcagaccggcgatttcaggacacctgccgagctggccctgaataagtttgagaaggagtccggccacatcagaaaccagaggggcgactatagccacaccttctcccgcaaggatctgcaggccgagctgatcctgctgttcgagaagcagaaggagtttggcaatccacacgtgagcggaggcctgaaggagggaatcgagaccctgctgatgacacagaggcctgccctgtccggcgacgcagtgcagaagatgctggggcactgcaccctcgagcctacagagccaaaggccgccaagaacacctacacagccgagcggtttatctggctgacaaagctgaacaatctgagaatcctggagcagggatccgagaggccactgaccgacacagagaggtccaccctgatggatgagccttaccggaagtctaaactgacatatgcccaggccagaaagctgctgggcctggaggacaccgccttctttaagggcctgagatacggcaaggataatgccgaggcctccacactgatggagatgaaggcctatcacgccatctctcgcgccctggagaaggagggcctgaaggacaagaagtcccccctgaacctgagctccgagctgcaggatgagatcggcaccgccttctctctgtttaagaccgacgaggatatcacaggccgcctgaaggacagggtgcagcctgagatcctggaggccctgctgaagcacatctctttcgataagtttgtgcagatcagcctgaaggccctgagaaggatcgtgccactgatggagcagggcaagcggtacgacgaggcctgcgccgagatctacggcgttcactatggcaagaagaacacagaggagaagatctatctgccccctatccctgccgacgagatcagaaatcctgtggtgctgagggccctgtcccaggcaagaaaagtgatcaacggagtggtgcgccggtacggatctccagcccggatccacatcgagaccgccagagaagtgggcaagagcttcaaggaccggaaggagatcgagaagagacaggaggagaatcgcaaggatcgggagaaggccgccgccaagtttagggagtacttccctaactttgtgggcgagccaaagtctaaggacatcctgaagctgcgcctgtacgagcagcagcacggcaagtgtctgtatagcggcaaggagatcaatctggtgcggctgaacgagaagggctatgtggagatcgatcacgccctgcctttctccagaacctgggacgattcttttaacaataaggtgctggtgctgggcagcgagaaccagaataagggcaatcagacaccatacgagtatttcaatggcaaggacaactccagggagtggcaggagttcaaggcccgcgtggagacctctagatttcccaggagcaagaagcagcggatcctgctgcagaagttcgacgaggatggctttaaggagtgcaacctgaatgacaccagatacgtgaaccggttcctgtgccagtttgtggccgatcacatcctgctgaccggcaagggcaagagaagggtgttcgcctctaatggccagatcacaaacctgctgaggggattttggggactgaggaaggtgcgggcagagaatgacagacaccacgcactggatgcagtggtggtggcatgcagcaccgtggcaatgcagcagaagatcacaagattcgtgaggtataaggagatgaacgcctttgacggcaagaccatcgataaggagacaggcaaggtgctgcaccagaagacccacttcccccagccttgggagttctttgcccaggaagtgatgatccgggtgttcggcaagccagacggcaagcctgagtttgaggaggccgataccccagagaagctgaggacactgctggcagagaagctgtctagcaggccagaggcagtgcacgagtacgtgaccccactgttcgtgtccagggcacccaatcggaagatgtctggcgcccacaaggacacactgagaagcgccaagaggtttgtgaagcacaacgagaagatctccgtgaagagagtgtggctgaccgagatcaagctggccgatctggagaacatggtgaattacaagaacggcagggagatcgagctgtatgaggccctgaaggcaaggctggaggcctacggaggaaatgccaagcaggcct tcgacccaaaggataaccccttttataagaagggaggacagctggtgaaggccgtgcgggtggagaagacccagaagagcggcgtgctgctgaataagaagaacgcctac acaatcgccgacaatggtgatatggtgagagtggacgtgttctgtaaggtggataagaagggcaagaatcagtactttatcgtgcctatctatgcctggcaggtggccgagaa catcctgccagacatcgattgcaagggctacagaatcgacgatagctatacattctgtttttccctgcacaagtatgacctgatcgccttccagaaggatgagaagtccaaggt ggagtttgcctactatatcaattgcgactcctctagcggcgggttctacctggcctggcacgataagggcagcagggagcagcggtttcgcatctccacccagaatctggcgct gatccagaagtatcaggtgaacgagctgggcaaggagatcaggccatgtcggctgaagaagcgcccacccgtgcgggaggataaaagaaccgccgacggcagcgaattc gagcccaagaagaagaggaaagtc

**Nme2-ABE8e-i1_linker10 (WT)**: <u>BPSV40-NLS</u>, nNme2Cas9, <u>TadA8e</u>, Linkers

MKRTADGSEFESPKKKRKVEDMAAFKPNPINYILGLAIGIASVGWAMVEIDEEENPIRLIDLGVRVFERAEVPKTGDSLAMARRLARSVRRLTRR RAHRLLRARRLLKREGVLQAADFDENGLIKSLPNTPWQLRAAALDRKLTPLEWSAVLLHLIKHRGYLSQRKNEGETADKELGALLKGVANNAHAL QTGDFRTPAELALNKFEKESGHIRNQRGDYSHTFSRKDLQAELILLFEKQKEFGNPHVSGGLKEGIETLLMTQRPALSGDAVQKMLGHCTFEPAE PKAAKNTYTAERFIWLTKLNNLRILEQSGGSGGSGGSSEVEFSHEYWMRHALTLAKRARDEREVPVGAVLVLNNRVIGEGWNRAIGLHDPTAH AEIMALRQGGLVMQNYRLIDATLYVTFEPCVMCAGAMIHSRIGRVVFGVRNSKRGAAGSLMNVLNYPGMNHRVEITEGILADECAALLCDFYR MPRQVFNAQKKAQSSINETPGTSESATGSERPLTDTERATLMDEPYRKSKLTYAQARKLLGLEDTAFFKGLRYGKDNAEASTLMEMKAYHAISR ALEKEGLKDKKSPLNLSSELQDEIGTAFSLFKTDEDITGRLKDRVQPEILEALLKHISFDKFVQISLKALRRIVPLMEQGKRYDEACAEIYGDHYGKKN TEEKIYLPPIPADEIRNPVVLRALSQARKVINGVVRRYGSPARIHIETAREVGKSFKDRKEIEKRQEENRKDREKAAAKFREYFPNFVGEPKSKDILKL RLYEQQHGKCLYSGKEINLVRLNEKGYVEIDHALPFSRTWDDSFNNKVLVLGSENQNKGNQTPYEYFNGKDNSREWQEFKARVETSRFPRSKK QRILLQKFDEDGFKECNLNDTRYVNRFLCQFVADHILLTGKGKRRVFASNGQITNLLRGFWGLRKVRAENDRHHALDAVVVACSTVAMQQKIT RFVRYKEMNAFDGKTIDKETGKVLHQKTHFPQPWEFFAQEVMIRVFGKPDGKPEFEEADTPEKLRTLLAEKLSSRPEAVHEYVTPLFVSRAPNRK MSGAHKDTLRSAKRFVKHNEKISVKRVWLTEIKLADLENMVNYKNGREIELYEALKARLEAYGGNAKQAFDPKDNPFYKKGGQLVKAVRVEKT QESGVLLNKKNAYTIADNGDMVRVDVFCKVDKKGKNQYFIVPIYAWQVAENILPDIDCKGYRIDDSYTFCFSLHKYDLIAFQKDEKSKVEFAYYIN CDSSNGRFYLAWHDKGSKEQQFRISTQNLVLIQKYQVNELGKEIRPCRLKKRPPVREDKRTADGSEFEPKKKRKV

atgaaacggacagccgacggaagcgagttcgagtcaccaaagaagaagcggaaagtcgaagatatggccgccttcaagcctaacccaatcaattacatcctgggactggccatcg gaatcgcatccgtgggatgggctatggtggagatcgacgaggaggagaatcctatccggctgatcgatctgggcgtgagagtgtttgagagggccgaggtgccaaagaccggcga ttctctggctatggcccggagactggcacggagcgtgaggcgcctgacacggagaagggcacacaggctgctgagggcacgccggctgctgaagagagagggcgtgctgcaggc agcagacttcgatgagaatggcctgatcaagagcctgccaaacaccccctggcagctgagagcagccgccctggacaggaagctgacaccactggagtggtctgccgtgctgctg cacctgatcaagcaccgcggctacctgagccagcggaagaacgagggagagacagcagacaaggagctgggcgccctgctgaagggagtggccaacaatgcccacgccctgca gaccggcgatttcaggacacctgccgagctggccctgaataagtttgagaaggagtccggccacatcagaaaccagaggggcgactatagccacaccttctcccgcaaggatctgc aggccgagctgatcctgctgttcgagaagcagaaggagtttggcaatccacacgtgagcggaggcctgaaggagggaatcgagaccctgctgatgacacagaggcctgccctgtc cggcgacgcagtgcagaagatgctgggacactgcaccttcgagcctgcagagccaaaggccgccaagaacacctacacagccgagcggtttatctggctgacaaagctgaacaat ctgagaatcctggagcagtctggcggttcaggtggatcaggcggtagctctgaggtggagttttcccacgagtactggatgagacatgccctgaccctggccaagagggcacgcga tgagagggaggtgcctgtgggagccgtgctggtgctgaacaatagagtgatcggcgagggctggaacagagccatcggcctgcacgacccaacagcccatgccgaaattatggcc ctgagacagggcggcctggtcatgcagaactacagactgattgacgccaccctgtacgtgacattcgagccttgcgtgatgtgcgccggcgccatgatccactctaggatcggccgc gtggtgtttggcgtgaggaacagcaaacggggcgccgcaggctccctgatgaacgtgctgaactaccccggcatgaatcaccgcgtcgaaattaccgagggaatcctggcagatg aatgtgccgccctgctgtgcgacttctaccggatgcctagacaggtgttcaatgctcagaagaaggcccagagctccatcaacgagacacctggcacaagcgagagcgcaacagg atccgagaggccactgaccgacacagagagggccaccctgatggatgagccttaccggaagtctaagctgacatatgcccaggccagaaagctgctgggcctggaggacaccgc cttctttaagggcctgagatacggcaaggataatgccgaggcctccacactgatggagatgaaggcctatcacgccatctctcgcgccctggagaaggagggcctgaaggacaag aagtcccccctgaacctgagctccgagctgcaggatgagatcggcaccgccttctctctgtttaagaccgacgaggatatcacaggccgcctgaaggacagggtgcagcctgagat cctggaggccctgctgaagcacatctctttcgataagtttgtgcagatcagcctgaaggccctgagaaggatcgtgccactgatggagcagggcaagcggtacgacgaggcctgcg ccgagatctacggcgatcactatggcaagaagaacacagaggagaagatctatctgccccctatccctgccgacgagatcagaaatcctgtggtgctgagggccctgtcccaggca agaaaagtgatcaacggagtggtgcgccggtacggatctccagcccggatccacatcgagaccgccagagaagtgggcaagagcttcaaggaccggaaggagatcgagaagag acaggaggagaatcgcaaggatcgggagaaggccgccgccaagtttagggagtacttccctaactttgtgggcgagccaaagtctaaggacatcctgaagctgcgcctgtacgag cagcagcacggcaagtgtctgtatagcggcaaggagatcaatctggtgcggctgaacgagaagggctatgtggagatcgatcacgccctgcctttctccagaacctgggacgattct tttaacaataaggtgctggtgctgggcagcgagaaccagaataagggcaatcagacaccatacgagtatttcaatggcaaggacaactccagggagtggcaggagttcaaggccc gcgtggagacctctagatttcccaggagcaagaagcagcggatcctgctgcagaagttcgacgaggatggctttaaggagtgcaacctgaatgacaccagatacgtgaaccggttc ctgtgccagtttgtggccgatcacatcctgctgaccggcaagggcaagagaagggtgttcgcctctaatggccagatcacaaacctgctgaggggattttggggactgaggaaggtg cgggcagagaatgacagacaccacgcactggatgcagtggtggtggcatgcagcaccgtggcaatgcagcagaagatcacaagattcgtgaggtataaggagatgaacgcctttg acggcaagaccatcgataaggagacaggcaaggtgctgcaccagaagacccacttcccccagccttgggagttctttgcccaggaagtgatgatccgggtgttcggcaagccaga cggcaagcctgagtttgaggaggccgataccccagagaagctgaggacactgctggcagagaagctgtctagcaggccagaggcagtgcacgagtacgtgaccccactgttcgtg tccagggcacccaatcggaagatgtctggcgcccacaaggacacactgagaagcgccaagaggtttgtgaagcacaacgagaagatctccgtgaagagagtgtggctgaccgag atcaagctggccgatctggagaacatggtgaattacaagaacggcagggagatcgagctgtatgaggccctgaaggcaaggctggaggcctacggaggaaatgccaagcaggcc ttcgacccaaaggataaccccttttataagaagggaggacagctggtgaaggccgtgcgggtggagaagacccaggagagcggcgtgctgctgaataagaagaacgcctacaca atcgccgacaatggcgatatggtgagagtggacgtgttctgtaaggtggataagaagggcaagaatcagtactttatcgtgcctatctatgcctggcaggtggccgagaacatcctg ccagacatcgattgcaagggctacagaatcgacgatagctatacattctgtttttccctgcacaagtatgacctgatcgccttccagaaggatgagaagtccaaggtggagtttgcct actatatcaattgcgactcctctaacggcaggttctacctggcctggcacgataagggcagcaaggagcagcagtttcgcatctccacccagaatctggtgctgatccagaagtatca ggtgaacgagctgggcaaggagatcaggccatgtcggctgaagaagcgcccacccgtgcgggaggataaaagaaccgccgacggcagcgaattcgagcccaagaagaagagg aaagtc

**Nme2-ABE8e-i1_linker10 (E932R)**: <u>BPSV40-NLS</u>, nNme2Cas9, <u>TadA8e</u>, Linkers

MKRTADGSEFESPKKKRKVEDMAAFKPNPINYILGLAIGIASVGWAMVEIDEEENPIRLIDLGVRVFERAEVPKTGDSLAMARRLARSVRRLTRR RAHRLLRARRLLKREGVLQAADFDENGLIKSLPNTPWQLRAAALDRKLTPLEWSAVLLHLIKHRGYLSQRKNEGETADKELGALLKGVANNAHAL QTGDFRTPAELALNKFEKESGHIRNQRGDYSHTFSRKDLQAELILLFEKQKEFGNPHVSGGLKEGIETLLMTQRPALSGDAVQKMLGHCTFEPAE PKAAKNTYTAERFIWLTKLNNLRILEQSGGSGGSGGSSEVEFSHEYWMRHALTLAKRARDEREVPVGAVLVLNNRVIGEGWNRAIGLHDPTAH AEIMALRQGGLVMQNYRLIDATLYVTFEPCVMCAGAMIHSRIGRVVFGVRNSKRGAAGSLMNVLNYPGMNHRVEITEGILADECAALLCDFYR MPRQVFNAQKKAQSSINETPGTSESATGSERPLTDTERATLMDEPYRKSKLTYAQARKLLGLEDTAFFKGLRYGKDNAEASTLMEMKAYHAISR ALEKEGLKDKKSPLNLSSELQDEIGTAFSLFKTDEDITGRLKDRVQPEILEALLKHISFDKFVQISLKALRRIVPLMEQGKRYDEACAEIYGDHYGKKN TEEKIYLPPIPADEIRNPVVLRALSQARKVINGVVRRYGSPARIHIETAREVGKSFKDRKEIEKRQEENRKDREKAAAKFREYFPNFVGEPKSKDILKL RLYEQQHGKCLYSGKEINLVRLNEKGYVEIDHALPFSRTWDDSFNNKVLVLGSENQNKGNQTPYEYFNGKDNSREWQEFKARVETSRFPRSKK QRILLQKFDEDGFKECNLNDTRYVNRFLCQFVADHILLTGKGKRRVFASNGQITNLLRGFWGLRKVRAENDRHHALDAVVVACSTVAMQQKIT RFVRYKEMNAFDGKTIDKETGKVLHQKTHFPQPWEFFAQEVMIRVFGKPDGKPEFEEADTPEKLRTLLAEKLSSRPEAVHEYVTPLFVSRAPNRK MSGAHKDTLRSAKRFVKHNEKISVKRVWLTEIKLADLENMVNYKNGREIELYEALKARLEAYGGNAKQAFDPKDNPFYKKGGQLVKAVRVEKT QRSGVLLNKKNAYTIADNGDMVRVDVFCKVDKKGKNQYFIVPIYAWQVAENILPDIDCKGYRIDDSYTFCFSLHKYDLIAFQKDEKSKVEFAYYIN CDSSNGRFYLAWHDKGSKEQQFRISTQNLVLIQKYQVNELGKEIRPCRLKKRPPVREDKRTADGSEFEPKKKRKV

atgaaacggacagccgacggaagcgagttcgagtcaccaaagaagaagcggaaagtcgaagatatggccgccttcaagcctaacccaatcaattacatcctgggactggccatcg gaatcgcatccgtgggatgggctatggtggagatcgacgaggaggagaatcctatccggctgatcgatctgggcgtgagagtgtttgagagggccgaggtgccaaagaccggcga ttctctggctatggcccggagactggcacggagcgtgaggcgcctgacacggagaagggcacacaggctgctgagggcacgccggctgctgaagagagagggcgtgctgcaggc agcagacttcgatgagaatggcctgatcaagagcctgccaaacaccccctggcagctgagagcagccgccctggacaggaagctgacaccactggagtggtctgccgtgctgctg cacctgatcaagcaccgcggctacctgagccagcggaagaacgagggagagacagcagacaaggagctgggcgccctgctgaagggagtggccaacaatgcccacgccctgca gaccggcgatttcaggacacctgccgagctggccctgaataagtttgagaaggagtccggccacatcagaaaccagaggggcgactatagccacaccttctcccgcaaggatctgc aggccgagctgatcctgctgttcgagaagcagaaggagtttggcaatccacacgtgagcggaggcctgaaggagggaatcgagaccctgctgatgacacagaggcctgccctgtc cggcgacgcagtgcagaagatgctgggacactgcaccttcgagcctgcagagccaaaggccgccaagaacacctacacagccgagcggtttatctggctgacaaagctgaacaat ctgagaatcctggagcagtctggcggttcaggtggatcaggcggtagctctgaggtggagttttcccacgagtactggatgagacatgccctgaccctggccaagagggcacgcga tgagagggaggtgcctgtgggagccgtgctggtgctgaacaatagagtgatcggcgagggctggaacagagccatcggcctgcacgacccaacagcccatgccgaaattatggcc ctgagacagggcggcctggtcatgcagaactacagactgattgacgccaccctgtacgtgacattcgagccttgcgtgatgtgcgccggcgccatgatccactctaggatcggccgc gtggtgtttggcgtgaggaacagcaaacggggcgccgcaggctccctgatgaacgtgctgaactaccccggcatgaatcaccgcgtcgaaattaccgagggaatcctggcagatg aatgtgccgccctgctgtgcgacttctaccggatgcctagacaggtgttcaatgctcagaagaaggcccagagctccatcaacgagacacctggcacaagcgagagcgcaacagg atccgagaggccactgaccgacacagagagggccaccctgatggatgagccttaccggaagtctaagctgacatatgcccaggccagaaagctgctgggcctggaggacaccgc cttctttaagggcctgagatacggcaaggataatgccgaggcctccacactgatggagatgaaggcctatcacgccatctctcgcgccctggagaaggagggcctgaaggacaag aagtcccccctgaacctgagctccgagctgcaggatgagatcggcaccgccttctctctgtttaagaccgacgaggatatcacaggccgcctgaaggacagggtgcagcctgagat cctggaggccctgctgaagcacatctctttcgataagtttgtgcagatcagcctgaaggccctgagaaggatcgtgccactgatggagcagggcaagcggtacgacgaggcctgcg ccgagatctacggcgatcactatggcaagaagaacacagaggagaagatctatctgccccctatccctgccgacgagatcagaaatcctgtggtgctgagggccctgtcccaggca agaaaagtgatcaacggagtggtgcgccggtacggatctccagcccggatccacatcgagaccgccagagaagtgggcaagagcttcaaggaccggaaggagatcgagaagag acaggaggagaatcgcaaggatcgggagaaggccgccgccaagtttagggagtacttccctaactttgtgggcgagccaaagtctaaggacatcctgaagctgcgcctgtacgag cagcagcacggcaagtgtctgtatagcggcaaggagatcaatctggtgcggctgaacgagaagggctatgtggagatcgatcacgccctgcctttctccagaacctgggacgattct tttaacaataaggtgctggtgctgggcagcgagaaccagaataagggcaatcagacaccatacgagtatttcaatggcaaggacaactccagggagtggcaggagttcaaggccc gcgtggagacctctagatttcccaggagcaagaagcagcggatcctgctgcagaagttcgacgaggatggctttaaggagtgcaacctgaatgacaccagatacgtgaaccggttc ctgtgccagtttgtggccgatcacatcctgctgaccggcaagggcaagagaagggtgttcgcctctaatggccagatcacaaacctgctgaggggattttggggactgaggaaggtg cgggcagagaatgacagacaccacgcactggatgcagtggtggtggcatgcagcaccgtggcaatgcagcagaagatcacaagattcgtgaggtataaggagatgaacgcctttg acggcaagaccatcgataaggagacaggcaaggtgctgcaccagaagacccacttcccccagccttgggagttctttgcccaggaagtgatgatccgggtgttcggcaagccaga cggcaagcctgagtttgaggaggccgataccccagagaagctgaggacactgctggcagagaagctgtctagcaggccagaggcagtgcacgagtacgtgaccccactgttcgtg tccagggcacccaatcggaagatgtctggcgcccacaaggacacactgagaagcgccaagaggtttgtgaagcacaacgagaagatctccgtgaagagagtgtggctgaccgag atcaagctggccgatctggagaacatggtgaattacaagaacggcagggagatcgagctgtatgaggccctgaaggcaaggctggaggcctacggaggaaatgccaagcaggcc ttcgacccaaaggataaccccttttataagaagggaggacagctggtgaaggccgtgcgggtggagaagacccagCGTagcggcgtgctgctgaataagaagaacgcctacaca atcgccgacaatggcgatatggtgagagtggacgtgttctgtaaggtggataagaagggcaagaatcagtactttatcgtgcctatctatgcctggcaggtggccgagaacatcctg ccagacatcgattgcaagggctacagaatcgacgatagctatacattctgtttttccctgcacaagtatgacctgatcgccttccagaaggatgagaagtccaaggtggagtttgcct actatatcaattgcgactcctctaacggcaggttctacctggcctggcacgataagggcagcaaggagcagcagtttcgcatctccacccagaatctggtgctgatccagaagtatca ggtgaacgagctgggcaaggagatcaggccatgtcggctgaagaagcgcccacccgtgcgggaggataaaagaaccgccgacggcagcgaattcgagcccaagaagaagagg aaagtc

**Nme2-ABE8e-i1_linker10 (D56R)**: <u>BPSV40-NLS</u>, nNme2Cas9, <u>TadA8e</u>, Linkers

MKRTADGSEFESPKKKRKVEDMAAFKPNPINYILGLAIGIASVGWAMVEIDEEENPIRLIDLGVRVFERAEVPKTGRSLAMARRLARSVRRLTRR RAHRLLRARRLLKREGVLQAADFDENGLIKSLPNTPWQLRAAALDRKLTPLEWSAVLLHLIKHRGYLSQRKNEGETADKELGALLKGVANNAHAL QTGDFRTPAELALNKFEKESGHIRNQRGDYSHTFSRKDLQAELILLFEKQKEFGNPHVSGGLKEGIETLLMTQRPALSGDAVQKMLGHCTFEPAE PKAAKNTYTAERFIWLTKLNNLRILEQSGGSGGSGGSSEVEFSHEYWMRHALTLAKRARDEREVPVGAVLVLNNRVIGEGWNRAIGLHDPTAH AEIMALRQGGLVMQNYRLIDATLYVTFEPCVMCAGAMIHSRIGRVVFGVRNSKRGAAGSLMNVLNYPGMNHRVEITEGILADECAALLCDFYR MPRQVFNAQKKAQSSINETPGTSESATGSERPLTDTERATLMDEPYRKSKLTYAQARKLLGLEDTAFFKGLRYGKDNAEASTLMEMKAYHAISR ALEKEGLKDKKSPLNLSSELQDEIGTAFSLFKTDEDITGRLKDRVQPEILEALLKHISFDKFVQISLKALRRIVPLMEQGKRYDEACAEIYGDHYGKKN TEEKIYLPPIPADEIRNPVVLRALSQARKVINGVVRRYGSPARIHIETAREVGKSFKDRKEIEKRQEENRKDREKAAAKFREYFPNFVGEPKSKDILKL RLYEQQHGKCLYSGKEINLVRLNEKGYVEIDHALPFSRTWDDSFNNKVLVLGSENQNKGNQTPYEYFNGKDNSREWQEFKARVETSRFPRSKK QRILLQKFDEDGFKECNLNDTRYVNRFLCQFVADHILLTGKGKRRVFASNGQITNLLRGFWGLRKVRAENDRHHALDAVVVACSTVAMQQKIT RFVRYKEMNAFDGKTIDKETGKVLHQKTHFPQPWEFFAQEVMIRVFGKPDGKPEFEEADTPEKLRTLLAEKLSSRPEAVHEYVTPLFVSRAPNRK MSGAHKDTLRSAKRFVKHNEKISVKRVWLTEIKLADLENMVNYKNGREIELYEALKARLEAYGGNAKQAFDPKDNPFYKKGGQLVKAVRVEKT QESGVLLNKKNAYTIADNGDMVRVDVFCKVDKKGKNQYFIVPIYAWQVAENILPDIDCKGYRIDDSYTFCFSLHKYDLIAFQKDEKSKVEFAYYIN CDSSNGRFYLAWHDKGSKEQQFRISTQNLVLIQKYQVNELGKEIRPCRLKKRPPVREDKRTADGSEFEPKKKRKV

atgaaacggacagccgacggaagcgagttcgagtcaccaaagaagaagcggaaagtcgaagatatggccgccttcaagcctaacccaatcaattacatcctgggactggccatcg gaatcgcatccgtgggatgggctatggtggagatcgacgaggaggagaatcctatccggctgatcgatctgggcgtgagagtgtttgagagggccgaggtgccaaagaccggcC GTtctctggctatggcccggagactggcacggagcgtgaggcgcctgacacggagaagggcacacaggctgctgagggcacgccggctgctgaagagagagggcgtgctgcag gcagcagacttcgatgagaatggcctgatcaagagcctgccaaacaccccctggcagctgagagcagccgccctggacaggaagctgacaccactggagtggtctgccgtgctgc tgcacctgatcaagcaccgcggctacctgagccagcggaagaacgagggagagacagcagacaaggagctgggcgccctgctgaagggagtggccaacaatgcccacgccctg cagaccggcgatttcaggacacctgccgagctggccctgaataagtttgagaaggagtccggccacatcagaaaccagaggggcgactatagccacaccttctcccgcaaggatct gcaggccgagctgatcctgctgttcgagaagcagaaggagtttggcaatccacacgtgagcggaggcctgaaggagggaatcgagaccctgctgatgacacagaggcctgccctg tccggcgacgcagtgcagaagatgctgggacactgcaccttcgagcctgcagagccaaaggccgccaagaacacctacacagccgagcggtttatctggctgacaaagctgaaca atctgagaatcctggagcagtctggcggttcaggtggatcaggcggtagctctgaggtggagttttcccacgagtactggatgagacatgccctgaccctggccaagagggcacgc gatgagagggaggtgcctgtgggagccgtgctggtgctgaacaatagagtgatcggcgagggctggaacagagccatcggcctgcacgacccaacagcccatgccgaaattatgg ccctgagacagggcggcctggtcatgcagaactacagactgattgacgccaccctgtacgtgacattcgagccttgcgtgatgtgcgccggcgccatgatccactctaggatcggcc gcgtggtgtttggcgtgaggaacagcaaacggggcgccgcaggctccctgatgaacgtgctgaactaccccggcatgaatcaccgcgtcgaaattaccgagggaatcctggcagat gaatgtgccgccctgctgtgcgacttctaccggatgcctagacaggtgttcaatgctcagaagaaggcccagagctccatcaacgagacacctggcacaagcgagagcgcaacag gatccgagaggccactgaccgacacagagagggccaccctgatggatgagccttaccggaagtctaagctgacatatgcccaggccagaaagctgctgggcctggaggacaccg ccttctttaagggcctgagatacggcaaggataatgccgaggcctccacactgatggagatgaaggcctatcacgccatctctcgcgccctggagaaggagggcctgaaggacaag aagtcccccctgaacctgagctccgagctgcaggatgagatcggcaccgccttctctctgtttaagaccgacgaggatatcacaggccgcctgaaggacagggtgcagcctgagat cctggaggccctgctgaagcacatctctttcgataagtttgtgcagatcagcctgaaggccctgagaaggatcgtgccactgatggagcagggcaagcggtacgacgaggcctgcg ccgagatctacggcgatcactatggcaagaagaacacagaggagaagatctatctgccccctatccctgccgacgagatcagaaatcctgtggtgctgagggccctgtcccaggca agaaaagtgatcaacggagtggtgcgccggtacggatctccagcccggatccacatcgagaccgccagagaagtgggcaagagcttcaaggaccggaaggagatcgagaagag acaggaggagaatcgcaaggatcgggagaaggccgccgccaagtttagggagtacttccctaactttgtgggcgagccaaagtctaaggacatcctgaagctgcgcctgtacgag cagcagcacggcaagtgtctgtatagcggcaaggagatcaatctggtgcggctgaacgagaagggctatgtggagatcgatcacgccctgcctttctccagaacctgggacgattct tttaacaataaggtgctggtgctgggcagcgagaaccagaataagggcaatcagacaccatacgagtatttcaatggcaaggacaactccagggagtggcaggagttcaaggccc gcgtggagacctctagatttcccaggagcaagaagcagcggatcctgctgcagaagttcgacgaggatggctttaaggagtgcaacctgaatgacaccagatacgtgaaccggttc ctgtgccagtttgtggccgatcacatcctgctgaccggcaagggcaagagaagggtgttcgcctctaatggccagatcacaaacctgctgaggggattttggggactgaggaaggtg cgggcagagaatgacagacaccacgcactggatgcagtggtggtggcatgcagcaccgtggcaatgcagcagaagatcacaagattcgtgaggtataaggagatgaacgcctttg acggcaagaccatcgataaggagacaggcaaggtgctgcaccagaagacccacttcccccagccttgggagttctttgcccaggaagtgatgatccgggtgttcggcaagccaga cggcaagcctgagtttgaggaggccgataccccagagaagctgaggacactgctggcagagaagctgtctagcaggccagaggcagtgcacgagtacgtgaccccactgttcgtg tccagggcacccaatcggaagatgtctggcgcccacaaggacacactgagaagcgccaagaggtttgtgaagcacaacgagaagatctccgtgaagagagtgtggctgaccgag atcaagctggccgatctggagaacatggtgaattacaagaacggcagggagatcgagctgtatgaggccctgaaggcaaggctggaggcctacggaggaaatgccaagcaggcc ttcgacccaaaggataaccccttttataagaagggaggacagctggtgaaggccgtgcgggtggagaagacccaggagagcggcgtgctgctgaataagaagaacgcctacaca atcgccgacaatggcgatatggtgagagtggacgtgttctgtaaggtggataagaagggcaagaatcagtactttatcgtgcctatctatgcctggcaggtggccgagaacatcctg ccagacatcgattgcaagggctacagaatcgacgatagctatacattctgtttttccctgcacaagtatgacctgatcgccttccagaaggatgagaagtccaaggtggagtttgcct actatatcaattgcgactcctctaacggcaggttctacctggcctggcacgataagggcagcaaggagcagcagtttcgcatctccacccagaatctggtgctgatccagaagtatca ggtgaacgagctgggcaaggagatcaggccatgtcggctgaagaagcgcccacccgtgcgggaggataaaagaaccgccgacggcagcgaattcgagcccaagaagaagagg aaagtc

**iNme2-ABE8e-i1_linker10**: BPSV40-NLS, nNme2Cas9, TadA8e, Linkers Jenifer Dounda Lab, iNme2Cas9 (D16A) variant in domain-inlaid-i1 format

MKRTADGSEFESPKKKRKVEDMAAFKPNPINYILGLAIGIASVGWAMVEIDEEENPIRLIDLGVRVFERAEVPKTGDSLAMARRLARSVRRLTRR RAHRLLRARRLLKREGVLQAADFDENGLIKSLPNTPWQLRAAALDRKLTPLEWSAVLLHLIKHRGYLSQRKNEGETADKELGALLKGVANNAHAL QTGDFRTPAELALNKFEKESGHIRNQRGDYSHTFSRKDLQAELILLFEKQKEFGNPHVSGGLKEGIETLLMTQRPALSGDAVQKMLGHCTFEPAE PKAAKNTYTAERFIWLTKLNNLRILEQSGGSGGSGGSSEVEFSHEYWMRHALTLAKRARDEREVPVGAVLVLNNRVIGEGWNRAIGLHDPTAH AEIMALRQGGLVMQNYRLIDATLYVTFEPCVMCAGAMIHSRIGRVVFGVRNSKRGAAGSLMNVLNYPGMNHRVEITEGILADECAALLCDFYR MPRQVFNAQKKAQSSINETPGTSESATGSERPLTDTERATLMDEPYRKSKLTYAQARKLLGLEDTAFFKGLRYGKDNAEASTLMEMKAYHAISR ALEKEGLKDKKSPLNLSSELQDEIGTAFSLFKTDEDITGRLKDRVQPEILEALLKHISFDKFVQISLKALRRIVPLMEQGKRYDEACAEIYGDHYGKKN TEEKIYLPPIPADEIRNPVVLRALSQARKVINGVVRRYGSPARIHIETAREVGKSFKDRKEIEKRQEENRKDREKAAAKFREYFPNFVGEPKSKDILKL RLYEQQHGKCLYSGKEINLVRLNEKGYVEIDHALPFSRTWDDSFNNKVLVLGSENQNKGNQTPYEYFNGKDNSREWQEFKARVETSRFPRSKK QRILLQKFDEDGFKECNLNDTRYVNRFLCQFVADHILLTGKGKRRVFASNGQITNLLRGFWGLRKVRAENDRHHALDAVVVACSTVAMQQKIT RFVRYKEMNAFDGKTIDKETGKVLHQKTHFPQPWEFFAQEVMIRVFGKPDGKPEFEEADTPEKLRTLLAEKLSSRPEAVHEYVTPLFVSRAPNRK MSGAHKGTLRSAKRFVKHNEKISVKRVWLTKIRLAALENMVNYKNGREIELYEALKARLEAYGGNAKQAFDPKGNPFYKKGGQLVKAVRVERT QKSGVLLNKKNAYTIADNGDMVRVDVFCKVDKKGKNQYFIVPIYAWQVAENILPDIDCKGYRIDDSYTFCFSLHKYDLIAFQKDEKSKVEFAYYIN CDSSNGRFYLAWHDKGSKEQQFRISTQNLVLIQKYQVNELGKEIRPCRLKKRPPVREDKRTADGSEFEPKKKRKV

atgaaacggacagccgacggaagcgagttcgagtcaccaaagaagaagcggaaagtcgaagatatggccgccttcaagcctaacccaatcaattacatcctgggactggccatcg gaatcgcatccgtgggatgggctatggtggagatcgacgaggaggagaatcctatccggctgatcgatctgggcgtgagagtgtttgagagggccgaggtgccaaagaccggcga ttctctggctatggcccggagactggcacggagcgtgaggcgcctgacacggagaagggcacacaggctgctgagggcacgccggctgctgaagagagagggcgtgctgcaggc agcagacttcgatgagaatggcctgatcaagagcctgccaaacaccccctggcagctgagagcagccgccctggacaggaagctgacaccactggagtggtctgccgtgctgctg cacctgatcaagcaccgcggctacctgagccagcggaagaacgagggagagacagcagacaaggagctgggcgccctgctgaagggagtggccaacaatgcccacgccctgca gaccggcgatttcaggacacctgccgagctggccctgaataagtttgagaaggagtccggccacatcagaaaccagaggggcgactatagccacaccttctcccgcaaggatctgc aggccgagctgatcctgctgttcgagaagcagaaggagtttggcaatccacacgtgagcggaggcctgaaggagggaatcgagaccctgctgatgacacagaggcctgccctgtc cggcgacgcagtgcagaagatgctgggacactgcaccttcgagcctgcagagccaaaggccgccaagaacacctacacagccgagcggtttatctggctgacaaagctgaacaat ctgagaatcctggagcagtctggcggttcaggtggatcaggcggtagctctgaggtggagttttcccacgagtactggatgagacatgccctgaccctggccaagagggcacgcga tgagagggaggtgcctgtgggagccgtgctggtgctgaacaatagagtgatcggcgagggctggaacagagccatcggcctgcacgacccaacagcccatgccgaaattatggcc ctgagacagggcggcctggtcatgcagaactacagactgattgacgccaccctgtacgtgacattcgagccttgcgtgatgtgcgccggcgccatgatccactctaggatcggccgc gtggtgtttggcgtgaggaacagcaaacggggcgccgcaggctccctgatgaacgtgctgaactaccccggcatgaatcaccgcgtcgaaattaccgagggaatcctggcagatg aatgtgccgccctgctgtgcgacttctaccggatgcctagacaggtgttcaatgctcagaagaaggcccagagctccatcaacgagacacctggcacaagcgagagcgcaacagg atccgagaggccactgaccgacacagagagggccaccctgatggatgagccttaccggaagtctaagctgacatatgcccaggccagaaagctgctgggcctggaggacaccgc cttctttaagggcctgagatacggcaaggataatgccgaggcctccacactgatggagatgaaggcctatcacgccatctctcgcgccctggagaaggagggcctgaaggacaag aagtcccccctgaacctgagctccgagctgcaggatgagatcggcaccgccttctctctgtttaagaccgacgaggatatcacaggccgcctgaaggacagggtgcagcctgagat cctggaggccctgctgaagcacatctctttcgataagtttgtgcagatcagcctgaaggccctgagaaggatcgtgccactgatggagcagggcaagcggtacgacgaggcctgcg ccgagatctacggcgatcactatggcaagaagaacacagaggagaagatctatctgccccctatccctgccgacgagatcagaaatcctgtggtgctgagggccctgtcccaggca agaaaagtgatcaacggagtggtgcgccggtacggatctccagcccggatccacatcgagaccgccagagaagtgggcaagagcttcaaggaccggaaggagatcgagaagag acaggaggagaatcgcaaggatcgggagaaggccgccgccaagtttagggagtacttccctaactttgtgggcgagccaaagtctaaggacatcctgaagctgcgcctgtacgag cagcagcacggcaagtgtctgtatagcggcaaggagatcaatctggtgcggctgaacgagaagggctatgtggagatcgatcacgccctgcctttctccagaacctgggacgattct tttaacaataaggtgctggtgctgggcagcgagaaccagaataagggcaatcagacaccatacgagtatttcaatggcaaggacaactccagggagtggcaggagttcaaggccc gcgtggagacctctagatttcccaggagcaagaagcagcggatcctgctgcagaagttcgacgaggatggctttaaggagtgcaacctgaatgacaccagatacgtgaaccggttc ctgtgccagtttgtggccgatcacatcctgctgaccggcaagggcaagagaagggtgttcgcctctaatggccagatcacaaacctgctgaggggattttggggactgaggaaggtg cgggcagagaatgacagacaccacgcactggatgcagtggtggtggcatgcagcaccgtggcaatgcagcagaagatcacaagattcgtgaggtataaggagatgaacgcctttg acggcaagaccatcgataaggagacaggcaaggtgctgcaccagaagacccacttcccccagccttgggagttctttgcccaggaagtgatgatccgggtgttcggcaagccaga cggcaagcctgagtttgaggaggccgataccccagagaagctgaggacactgctggcagagaagctgtctagcaggccagaggcagtgcacgagtacgtgaccccactgttcgtg tccagggcacccaatcggaagatgtctggcgcccacaagGGCacactgagaagcgccaagaggtttgtgaagcacaacgagaagatctccgtgaagagagtgtggctgaccAA GatcAGGctggccGCCctggagaacatggtgaattacaagaacggcagggagatcgagctgtatgaggccctgaaggcaaggctggaggcctacggaggaaatgccaagca ggccttcgacccaaagGGCaaccccttttataagaagggaggacagctggtgaaggccgtgcgggtggagAGGacccagAAGagcggcgtgctgctgaataagaagaacgc ctacacaatcgccgacaatggcgatatggtgagagtggacgtgttctgtaaggtggataagaagggcaagaatcagtactttatcgtgcctatctatgcctggcaggtggccgagaa catcctgccagacatcgattgcaagggctacagaatcgacgatagctatacattctgtttttccctgcacaagtatgacctgatcgccttccagaaggatgagaagtccaaggtggag tttgcctactatatcaattgcgactcctctaacggcaggttctacctggcctggcacgataagggcagcaaggagcagcagtttcgcatctccacccagaatctggtgctgatccaga agtatcaggtgaacgagctgggcaaggagatcaggccatgtcggctgaagaagcgcccacccgtgcgggaggataaaagaaccgccgacggcagcgaattcgagcccaagaag aagaggaaagtc

**Nme2^Smu^-ABE8e-i1_linker10 (WT)**: <u>BPSV40-NLS</u>, Nme2Cas9 – delta PID, <u>TadA8e</u>, <u>SmuCas9 PID</u>, Linkers

MKRTADGSEFESPKKKRKVEDMAAFKPNPINYILGLAIGIASVGWAMVEIDEEENPIRLIDLGVRVFERAEVPKTGDSLAMARRLARSVRRLTRR RAHRLLRARRLLKREGVLQAADFDENGLIKSLPNTPWQLRAAALDRKLTPLEWSAVLLHLIKHRGYLSQRKNEGETADKELGALLKGVANNAHAL QTGDFRTPAELALNKFEKESGHIRNQRGDYSHTFSRKDLQAELILLFEKQKEFGNPHVSGGLKEGIETLLMTQRPALSGDAVQKMLGHCTFEPAE PKAAKNTYTAERFIWLTKLNNLRILEQSGGSGGSGGSSEVEFSHEYWMRHALTLAKRARDEREVPVGAVLVLNNRVIGEGWNRAIGLHDPTAH AEIMALRQGGLVMQNYRLIDATLYVTFEPCVMCAGAMIHSRIGRVVFGVRNSKRGAAGSLMNVLNYPGMNHRVEITEGILADECAALLCDFYR MPRQVFNAQKKAQSSINETPGTSESATGSERPLTDTERATLMDEPYRKSKLTYAQARKLLGLEDTAFFKGLRYGKDNAEASTLMEMKAYHAISR ALEKEGLKDKKSPLNLSSELQDEIGTAFSLFKTDEDITGRLKDRVQPEILEALLKHISFDKFVQISLKALRRIVPLMEQGKRYDEACAEIYGDHYGKKN TEEKIYLPPIPADEIRNPVVLRALSQARKVINGVVRRYGSPARIHIETAREVGKSFKDRKEIEKRQEENRKDREKAAAKFREYFPNFVGEPKSKDILKL RLYEQQHGKCLYSGKEINLVRLNEKGYVEIDHALPFSRTWDDSFNNKVLVLGSENQNKGNQTPYEYFNGKDNSREWQEFKARVETSRFPRSKK QRILLQKFDEDGFKECNLNDTRYVNRFLCQFVADHILLTGKGKRRVFASNGQITNLLRGFWGLRKVRAENDRHHALDAVVVACSTVAMQQKIT RFVRYKEMNAFDGKTIDKETGKVLHQKTHFPQPWEFFAQEVMIRVFGKPDGKPEFEEADTPEKLRTLLAEKLSSRPEAVHEYVTPLFVSRAPNRK MSGAHKDTLRSAKRFVKHNEKISVKRVWLTEIKLADLENMVNYKNGREIELYEALKARLEAYGGNAKQAFDPKDNPFYKKGGQLVKAVRVEKT QESGVLLNKKNAYTIADNATMVRVDVYTKAGKNYLVPVYVWQVAQGILPNRAVTSGKSEADWDLIDESFEFKFSLSRGDLVEMISNKGRIFGYY NGLDRANGSIGIREHDLEKSKGKDGVHRVGVKTATAFNKYHVDPLGKEIHRCSSEPRPTLKIKSKKEDKRTADGSEFEPKKKRKV

atgaaacggacagccgacggaagcgagttcgagtcaccaaagaagaagcggaaagtcgaagatatggccgccttcaagcctaacccaatcaattacatcctgggactggccatcg gaatcgcatccgtgggatgggctatggtggagatcgacgaggaggagaatcctatccggctgatcgatctgggcgtgagagtgtttgagagggccgaggtgccaaagaccggcga ttctctggctatggcccggagactggcacggagcgtgaggcgcctgacacggagaagggcacacaggctgctgagggcacgccggctgctgaagagagagggcgtgctgcaggc agcagacttcgatgagaatggcctgatcaagagcctgccaaacaccccctggcagctgagagcagccgccctggacaggaagctgacaccactggagtggtctgccgtgctgctg cacctgatcaagcaccgcggctacctgagccagcggaagaacgagggagagacagcagacaaggagctgggcgccctgctgaagggagtggccaacaatgcccacgccctgca gaccggcgatttcaggacacctgccgagctggccctgaataagtttgagaaggagtccggccacatcagaaaccagaggggcgactatagccacaccttctcccgcaaggatctgc aggccgagctgatcctgctgttcgagaagcagaaggagtttggcaatccacacgtgagcggaggcctgaaggagggaatcgagaccctgctgatgacacagaggcctgccctgtc cggcgacgcagtgcagaagatgctgggacactgcaccttcgagcctgcagagccaaaggccgccaagaacacctacacagccgagcggtttatctggctgacaaagctgaacaat ctgagaatcctggagcagtctggcggttcaggtggatcaggcggtagctctgaggtggagttttcccacgagtactggatgagacatgccctgaccctggccaagagggcacgcga tgagagggaggtgcctgtgggagccgtgctggtgctgaacaatagagtgatcggcgagggctggaacagagccatcggcctgcacgacccaacagcccatgccgaaattatggcc ctgagacagggcggcctggtcatgcagaactacagactgattgacgccaccctgtacgtgacattcgagccttgcgtgatgtgcgccggcgccatgatccactctaggatcggccgc gtggtgtttggcgtgaggaacagcaaacggggcgccgcaggctccctgatgaacgtgctgaactaccccggcatgaatcaccgcgtcgaaattaccgagggaatcctggcagatg aatgtgccgccctgctgtgcgacttctaccggatgcctagacaggtgttcaatgctcagaagaaggcccagagctccatcaacgagacacctggcacaagcgagagcgcaacagg atccgagaggccactgaccgacacagagagggccaccctgatggatgagccttaccggaagtctaagctgacatatgcccaggccagaaagctgctgggcctggaggacaccgc cttctttaagggcctgagatacggcaaggataatgccgaggcctccacactgatggagatgaaggcctatcacgccatctctcgcgccctggagaaggagggcctgaaggacaag aagtcccccctgaacctgagctccgagctgcaggatgagatcggcaccgccttctctctgtttaagaccgacgaggatatcacaggccgcctgaaggacagggtgcagcctgagat cctggaggccctgctgaagcacatctctttcgataagtttgtgcagatcagcctgaaggccctgagaaggatcgtgccactgatggagcagggcaagcggtacgacgaggcctgcg ccgagatctacggcgatcactatggcaagaagaacacagaggagaagatctatctgccccctatccctgccgacgagatcagaaatcctgtggtgctgagggccctgtcccaggca agaaaagtgatcaacggagtggtgcgccggtacggatctccagcccggatccacatcgagaccgccagagaagtgggcaagagcttcaaggaccggaaggagatcgagaagag acaggaggagaatcgcaaggatcgggagaaggccgccgccaagtttagggagtacttccctaactttgtgggcgagccaaagtctaaggacatcctgaagctgcgcctgtacgag cagcagcacggcaagtgtctgtatagcggcaaggagatcaatctggtgcggctgaacgagaagggctatgtggagatcgatcacgccctgcctttctccagaacctgggacgattct tttaacaataaggtgctggtgctgggcagcgagaaccagaataagggcaatcagacaccatacgagtatttcaatggcaaggacaactccagggagtggcaggagttcaaggccc gcgtggagacctctagatttcccaggagcaagaagcagcggatcctgctgcagaagttcgacgaggatggctttaaggagtgcaacctgaatgacaccagatacgtgaaccggttc ctgtgccagtttgtggccgatcacatcctgctgaccggcaagggcaagagaagggtgttcgcctctaatggccagatcacaaacctgctgaggggattttggggactgaggaaggtg cgggcagagaatgacagacaccacgcactggatgcagtggtggtggcatgcagcaccgtggcaatgcagcagaagatcacaagattcgtgaggtataaggagatgaacgcctttg acggcaagaccatcgataaggagacaggcaaggtgctgcaccagaagacccacttcccccagccttgggagttctttgcccaggaagtgatgatccgggtgttcggcaagccaga cggcaagcctgagtttgaggaggccgataccccagagaagctgaggacactgctggcagagaagctgtctagcaggccagaggcagtgcacgagtacgtgaccccactgttcgtg tccagggcacccaatcggaagatgtctggcgcccacaaggacacactgagaagcgccaagaggtttgtgaagcacaacgagaagatctccgtgaagagagtgtggctgaccgag atcaagctggccgatctggagaacatggtgaattacaagaacggcagggagatcgagctgtatgaggccctgaaggcaaggctggaggcctacggaggaaatgccaagcaggcc ttcgacccaaaggataaccccttttataagaagggaggacagctggtgaaggccgtgcgggtggagaagacccaggagagcggcgtgctgctgaataagaagaacgcctacaca atcgccgacaacgccaccatggtgcgggtggacgtgtacaccaaggccggcaagaactacctggttcctgtgtacgtgtggcaggtggcccagggcatcttacccaaccgcgccgt gaccagcggcaagtccgaggctgactgggacctgatcgatgagagcttcgagttcaagttctctctgtcccggggagatctcgtggaaatgatctccaacaagggcagaatcttcgg ctactacaacggcctggacagagccaacggctctattggaattagagagcacgacctagagaagagcaagggcaaagacggcgtgcatagagtgggagtgaaaacagctacag catttaacaagtaccacgtggatcccctgggcaaagagatccacagatgcagcagcgaacccagacctacactgaaaatcaagtctaagaaggaggataaaagaaccgccgacg gcagcgaattcgagcccaagaagaagaggaaagtc

**Nme2^Smu^-ABE8e-i1_linker10 (E932R)**: <u>BPSV40-NLS</u>, Nme2Cas9 – delta PID, <u>TadA8e</u>, <u>SmuCas9 PID</u>, Linkers

MKRTADGSEFESPKKKRKVEDMAAFKPNPINYILGLAIGIASVGWAMVEIDEEENPIRLIDLGVRVFERAEVPKTGDSLAMARRLARSVRRLTRR RAHRLLRARRLLKREGVLQAADFDENGLIKSLPNTPWQLRAAALDRKLTPLEWSAVLLHLIKHRGYLSQRKNEGETADKELGALLKGVANNAHAL QTGDFRTPAELALNKFEKESGHIRNQRGDYSHTFSRKDLQAELILLFEKQKEFGNPHVSGGLKEGIETLLMTQRPALSGDAVQKMLGHCTFEPAE PKAAKNTYTAERFIWLTKLNNLRILEQSGGSGGSGGSSEVEFSHEYWMRHALTLAKRARDEREVPVGAVLVLNNRVIGEGWNRAIGLHDPTAH AEIMALRQGGLVMQNYRLIDATLYVTFEPCVMCAGAMIHSRIGRVVFGVRNSKRGAAGSLMNVLNYPGMNHRVEITEGILADECAALLCDFYR MPRQVFNAQKKAQSSINETPGTSESATGSERPLTDTERATLMDEPYRKSKLTYAQARKLLGLEDTAFFKGLRYGKDNAEASTLMEMKAYHAISR ALEKEGLKDKKSPLNLSSELQDEIGTAFSLFKTDEDITGRLKDRVQPEILEALLKHISFDKFVQISLKALRRIVPLMEQGKRYDEACAEIYGDHYGKKN TEEKIYLPPIPADEIRNPVVLRALSQARKVINGVVRRYGSPARIHIETAREVGKSFKDRKEIEKRQEENRKDREKAAAKFREYFPNFVGEPKSKDILKL RLYEQQHGKCLYSGKEINLVRLNEKGYVEIDHALPFSRTWDDSFNNKVLVLGSENQNKGNQTPYEYFNGKDNSREWQEFKARVETSRFPRSKK QRILLQKFDEDGFKECNLNDTRYVNRFLCQFVADHILLTGKGKRRVFASNGQITNLLRGFWGLRKVRAENDRHHALDAVVVACSTVAMQQKIT RFVRYKEMNAFDGKTIDKETGKVLHQKTHFPQPWEFFAQEVMIRVFGKPDGKPEFEEADTPEKLRTLLAEKLSSRPEAVHEYVTPLFVSRAPNRK MSGAHKDTLRSAKRFVKHNEKISVKRVWLTEIKLADLENMVNYKNGREIELYEALKARLEAYGGNAKQAFDPKDNPFYKKGGQLVKAVRVEKT QRSGVLLNKKNAYTIADNATMVRVDVYTKAGKNYLVPVYVWQVAQGILPNRAVTSGKSEADWDLIDESFEFKFSLSRGDLVEMISNKGRIFGYY NGLDRANGSIGIREHDLEKSKGKDGVHRVGVKTATAFNKYHVDPLGKEIHRCSSEPRPTLKIKSKKEDKRTADGSEFEPKKKRKV

atgaaacggacagccgacggaagcgagttcgagtcaccaaagaagaagcggaaagtcgaagatatggccgccttcaagcctaacccaatcaattacatcctgggactggccatcg gaatcgcatccgtgggatgggctatggtggagatcgacgaggaggagaatcctatccggctgatcgatctgggcgtgagagtgtttgagagggccgaggtgccaaagaccggcga ttctctggctatggcccggagactggcacggagcgtgaggcgcctgacacggagaagggcacacaggctgctgagggcacgccggctgctgaagagagagggcgtgctgcaggc agcagacttcgatgagaatggcctgatcaagagcctgccaaacaccccctggcagctgagagcagccgccctggacaggaagctgacaccactggagtggtctgccgtgctgctg cacctgatcaagcaccgcggctacctgagccagcggaagaacgagggagagacagcagacaaggagctgggcgccctgctgaagggagtggccaacaatgcccacgccctgca gaccggcgatttcaggacacctgccgagctggccctgaataagtttgagaaggagtccggccacatcagaaaccagaggggcgactatagccacaccttctcccgcaaggatctgc aggccgagctgatcctgctgttcgagaagcagaaggagtttggcaatccacacgtgagcggaggcctgaaggagggaatcgagaccctgctgatgacacagaggcctgccctgtc cggcgacgcagtgcagaagatgctgggacactgcaccttcgagcctgcagagccaaaggccgccaagaacacctacacagccgagcggtttatctggctgacaaagctgaacaat ctgagaatcctggagcagtctggcggttcaggtggatcaggcggtagctctgaggtggagttttcccacgagtactggatgagacatgccctgaccctggccaagagggcacgcga tgagagggaggtgcctgtgggagccgtgctggtgctgaacaatagagtgatcggcgagggctggaacagagccatcggcctgcacgacccaacagcccatgccgaaattatggcc ctgagacagggcggcctggtcatgcagaactacagactgattgacgccaccctgtacgtgacattcgagccttgcgtgatgtgcgccggcgccatgatccactctaggatcggccgc gtggtgtttggcgtgaggaacagcaaacggggcgccgcaggctccctgatgaacgtgctgaactaccccggcatgaatcaccgcgtcgaaattaccgagggaatcctggcagatg aatgtgccgccctgctgtgcgacttctaccggatgcctagacaggtgttcaatgctcagaagaaggcccagagctccatcaacgagacacctggcacaagcgagagcgcaacagg atccgagaggccactgaccgacacagagagggccaccctgatggatgagccttaccggaagtctaagctgacatatgcccaggccagaaagctgctgggcctggaggacaccgc cttctttaagggcctgagatacggcaaggataatgccgaggcctccacactgatggagatgaaggcctatcacgccatctctcgcgccctggagaaggagggcctgaaggacaag aagtcccccctgaacctgagctccgagctgcaggatgagatcggcaccgccttctctctgtttaagaccgacgaggatatcacaggccgcctgaaggacagggtgcagcctgagat cctggaggccctgctgaagcacatctctttcgataagtttgtgcagatcagcctgaaggccctgagaaggatcgtgccactgatggagcagggcaagcggtacgacgaggcctgcg ccgagatctacggcgatcactatggcaagaagaacacagaggagaagatctatctgccccctatccctgccgacgagatcagaaatcctgtggtgctgagggccctgtcccaggca agaaaagtgatcaacggagtggtgcgccggtacggatctccagcccggatccacatcgagaccgccagagaagtgggcaagagcttcaaggaccggaaggagatcgagaagag acaggaggagaatcgcaaggatcgggagaaggccgccgccaagtttagggagtacttccctaactttgtgggcgagccaaagtctaaggacatcctgaagctgcgcctgtacgag cagcagcacggcaagtgtctgtatagcggcaaggagatcaatctggtgcggctgaacgagaagggctatgtggagatcgatcacgccctgcctttctccagaacctgggacgattct tttaacaataaggtgctggtgctgggcagcgagaaccagaataagggcaatcagacaccatacgagtatttcaatggcaaggacaactccagggagtggcaggagttcaaggccc gcgtggagacctctagatttcccaggagcaagaagcagcggatcctgctgcagaagttcgacgaggatggctttaaggagtgcaacctgaatgacaccagatacgtgaaccggttc ctgtgccagtttgtggccgatcacatcctgctgaccggcaagggcaagagaagggtgttcgcctctaatggccagatcacaaacctgctgaggggattttggggactgaggaaggtg cgggcagagaatgacagacaccacgcactggatgcagtggtggtggcatgcagcaccgtggcaatgcagcagaagatcacaagattcgtgaggtataaggagatgaacgcctttg acggcaagaccatcgataaggagacaggcaaggtgctgcaccagaagacccacttcccccagccttgggagttctttgcccaggaagtgatgatccgggtgttcggcaagccaga cggcaagcctgagtttgaggaggccgataccccagagaagctgaggacactgctggcagagaagctgtctagcaggccagaggcagtgcacgagtacgtgaccccactgttcgtg tccagggcacccaatcggaagatgtctggcgcccacaaggacacactgagaagcgccaagaggtttgtgaagcacaacgagaagatctccgtgaagagagtgtggctgaccgag atcaagctggccgatctggagaacatggtgaattacaagaacggcagggagatcgagctgtatgaggccctgaaggcaaggctggaggcctacggaggaaatgccaagcaggcc ttcgacccaaaggataaccccttttataagaagggaggacagctggtgaaggccgtgcgggtggagaagacccagCGTagcggcgtgctgctgaataagaagaacgcctacaca atcgccgacaacgccaccatggtgcgggtggacgtgtacaccaaggccggcaagaactacctggttcctgtgtacgtgtggcaggtggcccagggcatcttacccaaccgcgccgt gaccagcggcaagtccgaggctgactgggacctgatcgatgagagcttcgagttcaagttctctctgtcccggggagatctcgtggaaatgatctccaacaagggcagaatcttcgg ctactacaacggcctggacagagccaacggctctattggaattagagagcacgacctagagaagagcaagggcaaagacggcgtgcatagagtgggagtgaaaacagctacag catttaacaagtaccacgtggatcccctgggcaaagagatccacagatgcagcagcgaacccagacctacactgaaaatcaagtctaagaaggaggataaaagaaccgccgacg gcagcgaattcgagcccaagaagaagaggaaagtc

**Nme2^Smu^-ABE8e-i1_linker10 (D56R)**: <u>BPSV40-NLS</u>, Nme2Cas9 – delta PID, <u>TadA8e</u>, <u>SmuCas9 PID</u>, Linkers

MKRTADGSEFESPKKKRKVEDMAAFKPNPINYILGLAIGIASVGWAMVEIDEEENPIRLIDLGVRVFERAEVPKTGRSLAMARRLARSVRRLTRR RAHRLLRARRLLKREGVLQAADFDENGLIKSLPNTPWQLRAAALDRKLTPLEWSAVLLHLIKHRGYLSQRKNEGETADKELGALLKGVANNAHAL QTGDFRTPAELALNKFEKESGHIRNQRGDYSHTFSRKDLQAELILLFEKQKEFGNPHVSGGLKEGIETLLMTQRPALSGDAVQKMLGHCTFEPAE PKAAKNTYTAERFIWLTKLNNLRILEQSGGSGGSGGSSEVEFSHEYWMRHALTLAKRARDEREVPVGAVLVLNNRVIGEGWNRAIGLHDPTAH AEIMALRQGGLVMQNYRLIDATLYVTFEPCVMCAGAMIHSRIGRVVFGVRNSKRGAAGSLMNVLNYPGMNHRVEITEGILADECAALLCDFYR MPRQVFNAQKKAQSSINETPGTSESATGSERPLTDTERATLMDEPYRKSKLTYAQARKLLGLEDTAFFKGLRYGKDNAEASTLMEMKAYHAISR ALEKEGLKDKKSPLNLSSELQDEIGTAFSLFKTDEDITGRLKDRVQPEILEALLKHISFDKFVQISLKALRRIVPLMEQGKRYDEACAEIYGDHYGKKN TEEKIYLPPIPADEIRNPVVLRALSQARKVINGVVRRYGSPARIHIETAREVGKSFKDRKEIEKRQEENRKDREKAAAKFREYFPNFVGEPKSKDILKL RLYEQQHGKCLYSGKEINLVRLNEKGYVEIDHALPFSRTWDDSFNNKVLVLGSENQNKGNQTPYEYFNGKDNSREWQEFKARVETSRFPRSKK QRILLQKFDEDGFKECNLNDTRYVNRFLCQFVADHILLTGKGKRRVFASNGQITNLLRGFWGLRKVRAENDRHHALDAVVVACSTVAMQQKIT RFVRYKEMNAFDGKTIDKETGKVLHQKTHFPQPWEFFAQEVMIRVFGKPDGKPEFEEADTPEKLRTLLAEKLSSRPEAVHEYVTPLFVSRAPNRK MSGAHKDTLRSAKRFVKHNEKISVKRVWLTEIKLADLENMVNYKNGREIELYEALKARLEAYGGNAKQAFDPKDNPFYKKGGQLVKAVRVEKT QESGVLLNKKNAYTIADNATMVRVDVYTKAGKNYLVPVYVWQVAQGILPNRAVTSGKSEADWDLIDESFEFKFSLSRGDLVEMISNKGRIFGYY NGLDRANGSIGIREHDLEKSKGKDGVHRVGVKTATAFNKYHVDPLGKEIHRCSSEPRPTLKIKSKKEDKRTADGSEFEPKKKRKV

atgaaacggacagccgacggaagcgagttcgagtcaccaaagaagaagcggaaagtcgaagatatggccgccttcaagcctaacccaatcaattacatcctgggactggccatcg gaatcgcatccgtgggatgggctatggtggagatcgacgaggaggagaatcctatccggctgatcgatctgggcgtgagagtgtttgagagggccgaggtgccaaagaccggcC GTtctctggctatggcccggagactggcacggagcgtgaggcgcctgacacggagaagggcacacaggctgctgagggcacgccggctgctgaagagagagggcgtgctgcag gcagcagacttcgatgagaatggcctgatcaagagcctgccaaacaccccctggcagctgagagcagccgccctggacaggaagctgacaccactggagtggtctgccgtgctgc tgcacctgatcaagcaccgcggctacctgagccagcggaagaacgagggagagacagcagacaaggagctgggcgccctgctgaagggagtggccaacaatgcccacgccctg cagaccggcgatttcaggacacctgccgagctggccctgaataagtttgagaaggagtccggccacatcagaaaccagaggggcgactatagccacaccttctcccgcaaggatct gcaggccgagctgatcctgctgttcgagaagcagaaggagtttggcaatccacacgtgagcggaggcctgaaggagggaatcgagaccctgctgatgacacagaggcctgccctg tccggcgacgcagtgcagaagatgctgggacactgcaccttcgagcctgcagagccaaaggccgccaagaacacctacacagccgagcggtttatctggctgacaaagctgaaca atctgagaatcctggagcagtctggcggttcaggtggatcaggcggtagctctgaggtggagttttcccacgagtactggatgagacatgccctgaccctggccaagagggcacgc gatgagagggaggtgcctgtgggagccgtgctggtgctgaacaatagagtgatcggcgagggctggaacagagccatcggcctgcacgacccaacagcccatgccgaaattatgg ccctgagacagggcggcctggtcatgcagaactacagactgattgacgccaccctgtacgtgacattcgagccttgcgtgatgtgcgccggcgccatgatccactctaggatcggcc gcgtggtgtttggcgtgaggaacagcaaacggggcgccgcaggctccctgatgaacgtgctgaactaccccggcatgaatcaccgcgtcgaaattaccgagggaatcctggcagat gaatgtgccgccctgctgtgcgacttctaccggatgcctagacaggtgttcaatgctcagaagaaggcccagagctccatcaacgagacacctggcacaagcgagagcgcaacag gatccgagaggccactgaccgacacagagagggccaccctgatggatgagccttaccggaagtctaagctgacatatgcccaggccagaaagctgctgggcctggaggacaccg ccttctttaagggcctgagatacggcaaggataatgccgaggcctccacactgatggagatgaaggcctatcacgccatctctcgcgccctggagaaggagggcctgaaggacaag aagtcccccctgaacctgagctccgagctgcaggatgagatcggcaccgccttctctctgtttaagaccgacgaggatatcacaggccgcctgaaggacagggtgcagcctgagat cctggaggccctgctgaagcacatctctttcgataagtttgtgcagatcagcctgaaggccctgagaaggatcgtgccactgatggagcagggcaagcggtacgacgaggcctgcg ccgagatctacggcgatcactatggcaagaagaacacagaggagaagatctatctgccccctatccctgccgacgagatcagaaatcctgtggtgctgagggccctgtcccaggca agaaaagtgatcaacggagtggtgcgccggtacggatctccagcccggatccacatcgagaccgccagagaagtgggcaagagcttcaaggaccggaaggagatcgagaagag acaggaggagaatcgcaaggatcgggagaaggccgccgccaagtttagggagtacttccctaactttgtgggcgagccaaagtctaaggacatcctgaagctgcgcctgtacgag cagcagcacggcaagtgtctgtatagcggcaaggagatcaatctggtgcggctgaacgagaagggctatgtggagatcgatcacgccctgcctttctccagaacctgggacgattct tttaacaataaggtgctggtgctgggcagcgagaaccagaataagggcaatcagacaccatacgagtatttcaatggcaaggacaactccagggagtggcaggagttcaaggccc gcgtggagacctctagatttcccaggagcaagaagcagcggatcctgctgcagaagttcgacgaggatggctttaaggagtgcaacctgaatgacaccagatacgtgaaccggttc ctgtgccagtttgtggccgatcacatcctgctgaccggcaagggcaagagaagggtgttcgcctctaatggccagatcacaaacctgctgaggggattttggggactgaggaaggtg cgggcagagaatgacagacaccacgcactggatgcagtggtggtggcatgcagcaccgtggcaatgcagcagaagatcacaagattcgtgaggtataaggagatgaacgcctttg acggcaagaccatcgataaggagacaggcaaggtgctgcaccagaagacccacttcccccagccttgggagttctttgcccaggaagtgatgatccgggtgttcggcaagccaga cggcaagcctgagtttgaggaggccgataccccagagaagctgaggacactgctggcagagaagctgtctagcaggccagaggcagtgcacgagtacgtgaccccactgttcgtg tccagggcacccaatcggaagatgtctggcgcccacaaggacacactgagaagcgccaagaggtttgtgaagcacaacgagaagatctccgtgaagagagtgtggctgaccgag atcaagctggccgatctggagaacatggtgaattacaagaacggcagggagatcgagctgtatgaggccctgaaggcaaggctggaggcctacggaggaaatgccaagcaggcc ttcgacccaaaggataaccccttttataagaagggaggacagctggtgaaggccgtgcgggtggagaagacccaggagagcggcgtgctgctgaataagaagaacgcctacaca atcgccgacaacgccaccatggtgcgggtggacgtgtacaccaaggccggcaagaactacctggttcctgtgtacgtgtggcaggtggcccagggcatcttacccaaccgcgccgt gaccagcggcaagtccgaggctgactgggacctgatcgatgagagcttcgagttcaagttctctctgtcccggggagatctcgtggaaatgatctccaacaagggcagaatcttcgg ctactacaacggcctggacagagccaacggctctattggaattagagagcacgacctagagaagagcaagggcaaagacggcgtgcatagagtgggagtgaaaacagctacag catttaacaagtaccacgtggatcccctgggcaaagagatccacagatgcagcagcgaacccagacctacactgaaaatcaagtctaagaaggaggataaaagaaccgccgacg gcagcgaattcgagcccaagaagaagaggaaagtc

**iNme2^Smu^-ABE8e-i1_linker10**: BPSV40-NLS, iNme2Cas9 – delta PID, TadA8e, SmuCas9 PID, Linkers Jenifer Dounda Lab, iNme2Cas9 (D16A) variant with SmuCas9 PID swap in domain-inlaid-i1 format

MKRTADGSEFESPKKKRKVEDMAAFKPNPINYILGLAIGIASVGWAMVEIDEEENPIRLIDLGVRVFERAEVPKTGDSLAMARRLARSVRRLTRR RAHRLLRARRLLKREGVLQAADFDENGLIKSLPNTPWQLRAAALDRKLTPLEWSAVLLHLIKHRGYLSQRKNEGETADKELGALLKGVANNAHAL QTGDFRTPAELALNKFEKESGHIRNQRGDYSHTFSRKDLQAELILLFEKQKEFGNPHVSGGLKEGIETLLMTQRPALSGDAVQKMLGHCTFEPAE PKAAKNTYTAERFIWLTKLNNLRILEQSGGSGGSGGSSEVEFSHEYWMRHALTLAKRARDEREVPVGAVLVLNNRVIGEGWNRAIGLHDPTAH AEIMALRQGGLVMQNYRLIDATLYVTFEPCVMCAGAMIHSRIGRVVFGVRNSKRGAAGSLMNVLNYPGMNHRVEITEGILADECAALLCDFYR MPRQVFNAQKKAQSSINETPGTSESATGSERPLTDTERATLMDEPYRKSKLTYAQARKLLGLEDTAFFKGLRYGKDNAEASTLMEMKAYHAISR ALEKEGLKDKKSPLNLSSELQDEIGTAFSLFKTDEDITGRLKDRVQPEILEALLKHISFDKFVQISLKALRRIVPLMEQGKRYDEACAEIYGDHYGKKN TEEKIYLPPIPADEIRNPVVLRALSQARKVINGVVRRYGSPARIHIETAREVGKSFKDRKEIEKRQEENRKDREKAAAKFREYFPNFVGEPKSKDILKL RLYEQQHGKCLYSGKEINLVRLNEKGYVEIDHALPFSRTWDDSFNNKVLVLGSENQNKGNQTPYEYFNGKDNSREWQEFKARVETSRFPRSKK QRILLQKFDEDGFKECNLNDTRYVNRFLCQFVADHILLTGKGKRRVFASNGQITNLLRGFWGLRKVRAENDRHHALDAVVVACSTVAMQQKIT RFVRYKEMNAFDGKTIDKETGKVLHQKTHFPQPWEFFAQEVMIRVFGKPDGKPEFEEADTPEKLRTLLAEKLSSRPEAVHEYVTPLFVSRAPNRK MSGAHKGTLRSAKRFVKHNEKISVKRVWLTKIRLAALENMVNYKNGREIELYEALKARLEAYGGNAKQAFDPKGNPFYKKGGQLVKAVRVERT QKSGVLLNKKNAYTIADNATMVRVDVYTKAGKNYLVPVYVWQVAQGILPNRAVTSGKSEADWDLIDESFEFKFSLSRGDLVEMISNKGRIFGYY NGLDRANGSIGIREHDLEKSKGKDGVHRVGVKTATAFNKYHVDPLGKEIHRCSSEPRPTLKIKSKKEDKRTADGSEFEPKKKRKV

atgaaacggacagccgacggaagcgagttcgagtcaccaaagaagaagcggaaagtcgaagatatggccgccttcaagcctaacccaatcaattacatcctgggactggccatcg gaatcgcatccgtgggatgggctatggtggagatcgacgaggaggagaatcctatccggctgatcgatctgggcgtgagagtgtttgagagggccgaggtgccaaagaccggcga ttctctggctatggcccggagactggcacggagcgtgaggcgcctgacacggagaagggcacacaggctgctgagggcacgccggctgctgaagagagagggcgtgctgcaggc agcagacttcgatgagaatggcctgatcaagagcctgccaaacaccccctggcagctgagagcagccgccctggacaggaagctgacaccactggagtggtctgccgtgctgctg cacctgatcaagcaccgcggctacctgagccagcggaagaacgagggagagacagcagacaaggagctgggcgccctgctgaagggagtggccaacaatgcccacgccctgca gaccggcgatttcaggacacctgccgagctggccctgaataagtttgagaaggagtccggccacatcagaaaccagaggggcgactatagccacaccttctcccgcaaggatctgc aggccgagctgatcctgctgttcgagaagcagaaggagtttggcaatccacacgtgagcggaggcctgaaggagggaatcgagaccctgctgatgacacagaggcctgccctgtc cggcgacgcagtgcagaagatgctgggacactgcaccttcgagcctgcagagccaaaggccgccaagaacacctacacagccgagcggtttatctggctgacaaagctgaacaat ctgagaatcctggagcagtctggcggttcaggtggatcaggcggtagctctgaggtggagttttcccacgagtactggatgagacatgccctgaccctggccaagagggcacgcga tgagagggaggtgcctgtgggagccgtgctggtgctgaacaatagagtgatcggcgagggctggaacagagccatcggcctgcacgacccaacagcccatgccgaaattatggcc ctgagacagggcggcctggtcatgcagaactacagactgattgacgccaccctgtacgtgacattcgagccttgcgtgatgtgcgccggcgccatgatccactctaggatcggccgc gtggtgtttggcgtgaggaacagcaaacggggcgccgcaggctccctgatgaacgtgctgaactaccccggcatgaatcaccgcgtcgaaattaccgagggaatcctggcagatg aatgtgccgccctgctgtgcgacttctaccggatgcctagacaggtgttcaatgctcagaagaaggcccagagctccatcaacgagacacctggcacaagcgagagcgcaacagg atccgagaggccactgaccgacacagagagggccaccctgatggatgagccttaccggaagtctaagctgacatatgcccaggccagaaagctgctgggcctggaggacaccgc cttctttaagggcctgagatacggcaaggataatgccgaggcctccacactgatggagatgaaggcctatcacgccatctctcgcgccctggagaaggagggcctgaaggacaag aagtcccccctgaacctgagctccgagctgcaggatgagatcggcaccgccttctctctgtttaagaccgacgaggatatcacaggccgcctgaaggacagggtgcagcctgagat cctggaggccctgctgaagcacatctctttcgataagtttgtgcagatcagcctgaaggccctgagaaggatcgtgccactgatggagcagggcaagcggtacgacgaggcctgcg ccgagatctacggcgatcactatggcaagaagaacacagaggagaagatctatctgccccctatccctgccgacgagatcagaaatcctgtggtgctgagggccctgtcccaggca agaaaagtgatcaacggagtggtgcgccggtacggatctccagcccggatccacatcgagaccgccagagaagtgggcaagagcttcaaggaccggaaggagatcgagaagag acaggaggagaatcgcaaggatcgggagaaggccgccgccaagtttagggagtacttccctaactttgtgggcgagccaaagtctaaggacatcctgaagctgcgcctgtacgag cagcagcacggcaagtgtctgtatagcggcaaggagatcaatctggtgcggctgaacgagaagggctatgtggagatcgatcacgccctgcctttctccagaacctgggacgattct tttaacaataaggtgctggtgctgggcagcgagaaccagaataagggcaatcagacaccatacgagtatttcaatggcaaggacaactccagggagtggcaggagttcaaggccc gcgtggagacctctagatttcccaggagcaagaagcagcggatcctgctgcagaagttcgacgaggatggctttaaggagtgcaacctgaatgacaccagatacgtgaaccggttc ctgtgccagtttgtggccgatcacatcctgctgaccggcaagggcaagagaagggtgttcgcctctaatggccagatcacaaacctgctgaggggattttggggactgaggaaggtg cgggcagagaatgacagacaccacgcactggatgcagtggtggtggcatgcagcaccgtggcaatgcagcagaagatcacaagattcgtgaggtataaggagatgaacgcctttg acggcaagaccatcgataaggagacaggcaaggtgctgcaccagaagacccacttcccccagccttgggagttctttgcccaggaagtgatgatccgggtgttcggcaagccaga cggcaagcctgagtttgaggaggccgataccccagagaagctgaggacactgctggcagagaagctgtctagcaggccagaggcagtgcacgagtacgtgaccccactgttcgtg tccagggcacccaatcggaagatgtctggcgcccacaagGGCacactgagaagcgccaagaggtttgtgaagcacaacgagaagatctccgtgaagagagtgtggctgaccAA GatcAGGctggccGCCctggagaacatggtgaattacaagaacggcagggagatcgagctgtatgaggccctgaaggcaaggctggaggcctacggaggaaatgccaagca ggccttcgacccaaagGGCaaccccttttataagaagggaggacagctggtgaaggccgtgcgggtggagAGGacccagAAGagcggcgtgctgctgaataagaagaacgc ctacacaatcgccgacaacgccaccatggtgcgggtggacgtgtacaccaaggccggcaagaactacctggttcctgtgtacgtgtggcaggtggcccagggcatcttacccaacc gcgccgtgaccagcggcaagtccgaggctgactgggacctgatcgatgagagcttcgagttcaagttctctctgtcccggggagatctcgtggaaatgatctccaacaagggcaga atcttcggctactacaacggcctggacagagccaacggctctattggaattagagagcacgacctagagaagagcaagggcaaagacggcgtgcatagagtgggagtgaaaaca gctacagcatttaacaagtaccacgtggatcccctgggcaaagagatccacagatgcagcagcgaacccagacctacactgaaaatcaagtctaagaaggaggataaaagaacc gccgacggcagcgaattcgagcccaagaagaagaggaaagtc

**eNme2-C-ABE8e-i1_linker10**: BPSV40-NLS, nNme2Cas9, TadA8e, Linkers David Liu Lab, eNme2-C ABE8e variant in domain-inlaid-i1 format

MKRTADGSEFESPKKKRKVEDAAFKSNPINYILGLAIGIASVGWAMVEIDEEGNPIRLIDLGVRVFERAEVPKTGDSLAMARRLARSVRRLTRRRA HRLLRARRLLKREGVLQAADFDENGLITSLPNTPWQLRAAALDRKLTPLEWSAVLLHLIKHRGYLSQRKNEGETAAKELGALLKGVANNAHALQT GDFRTPAELALNKFEKESGHIRNQRGDYSHTFSRKDLQAELILLFEKQKEFGNPHVSGGLKEGIETLLMTQRPALSGDAVQKMLGHCTLEPTEPK AAKNTYTAERFIWLTKLNNLRILEQSGGSGGSGGSSEVEFSHEYWMRHALTLAKRARDEREVPVGAVLVLNNRVIGEGWNRAIGLHDPTAHAEI MALRQGGLVMQNYRLIDATLYVTFEPCVMCAGAMIHSRIGRVVFGVRNSKRGAAGSLMNVLNYPGMNHRVEITEGILADECAALLCDFYRMP RQVFNAQKKAQSSINETPGTSESATGSERPLTDTERSTLMDEPYRKSKLTYAQARKLLGLEDTAFFKGLRYGKDNAEASTLMEMKAYHAISRALE KEGLKDKKSPLNLSSELQDEIGTAFSLFKTDEDITGRLKDRVQPEILEALLKHISFDKFVQISLKALRRIVPLMEQGKRYDEACAEIYGVHYGKKNTEEKIYLPPIPADEIRNPVVLRALSQARKVINGVVRRYGSPARIHIETAREVGKSFKDRKEIAKRQEENRKDREKAAAKFREYFPNFVGEPKSKDILKLRLYEQQHGKCLYSGKEINLVRLNEKGYVEIDHALPFSRTWDDSFNNKVLVLGSENQNKGNQTPYEYFNGKDNSREWQEFKARVETSRFPSSKKQRILLQKFDEDGFKECNLNDTRYVNRFLCQFVADHILLTGKGKRRVVASNGQITNLLRGFWRLRKVRAENDRHHALDAVVVACSTVAMQQKITRFVRYKEMNAFDGKTVDKETGKVLYQKTHFPQPWEFFAQEVMIRVFGKPDGKPEFEEADTPEKLRTLLAEKLSSRPEAVHEYVTPLFVSRAPNRKMSGAHKDTLRSAKRFVKHNEKISVKRVWLTEIKLADLENMVNYKNGREIELYEALKARLEAYGGNAKQAFDPKDNPFYKKGGQLVKAVRVEKTQKSGVLLNKKNAYTIADNGDMVRVDVFCKVDKKGKNQYFIVPIYAWQVAENILPDIDCKGYRIDDSYTFCFSLHKYDLIAFQKDEKSKVEFAYYINCDSSSGGFYLAWHDKGSREQRFRISTQNLALIQKYQVNELGKEIRPCRLKKRPPVRSGGSKRTADGSEFEPKKKRKV

atgaaacggacagccgacggaagcgagttcgagtcaccaaagaagaagcggaaagtcgaagatgcagcattcaagtcaaacccaatcaattacatcctgggactggcaatcggaatcgcatccgtgggatgggctatggtggagatcgacgaggaggggaatcctatccggctgatcgatctgggcgtgagagtgtttgagagggccgaggtgccaaagaccggcgattctctggctatggcccggagactggcacggagcgtgaggcgcctgacacggagaagggcacacaggctgctgagggcacgccggctgctgaagagagagggcgtgctgcaggcagcagacttcgatgagaatggcctgatcacgagcttgccaaacaccccctggcagctgagagcagccgccctggacaggaagctgacaccactggagtggtctgccgtgctgctgcacctgatcaagcaccgcggctacctgagccagcggaagaacgagggagagacagcagccaaggagctgggcgccctgctgaagggagtggccaacaatgcccacgccctgcagaccggcgatttcaggacacctgccgagctggccctgaataagtttgagaaggagtccggccacatcagaaaccagaggggcgactatagccacaccttctcccgcaaggatctgcaggccgagctgatcctgctgttcgagaagcagaaggagtttggcaatccacacgtgagcggaggcctgaaggagggaatcgagaccctgctgatgacacagaggcctgccctgtccggcgacgcagtgcagaagatgctggggcactgcaccctcgagcctacagagccaaaggccgccaagaacacctacacagccgagcggtttatctggctgacaaagctgaacaatctgagaatcctggagcagtctggcggttcaggtggatcaggcggtagctctgaggtggagttttcccacgagtactggatgagacatgccctgaccctggccaagagggcacgcgatgagagggaggtgcctgtgggagccgtgctggtgctgaacaatagagtgatcggcgagggctggaacagagccatcggcctgcacgacccaacagcccatgccgaaattatggccctgagacagggcggcctggtcatgcagaactacagactgattgacgccaccctgtacgtgacattcgagccttgcgtgatgtgcgccggcgccatgatccactctaggatcggccgcgtggtgtttggcgtgaggaacagcaaacggggcgccgcaggctccctgatgaacgtgctgaactaccccggcatgaatcaccgcgtcgaaattaccgagggaatcctggcagatgaatgtgccgccctgctgtgcgacttctaccggatgcctagacaggtgttcaatgctcagaagaaggcccagagctccatcaacgagacacctggcacaagcgagagcgcaacaggatccgagaggccactgaccgacacagagaggtccaccctgatggatgagccttaccggaagtctaaactgacatatgcccaggccagaaagctgctgggcctggaggacaccgccttctttaagggcctgagatacggcaaggataatgccgaggcctccacactgatggagatgaaggcctatcacgccatctctcgcgccctggagaaggagggcctgaaggacaagaagtcccccctgaacctgagctccgagctgcaggatgagatcggcaccgccttctctctgtttaagaccgacgaggatatcacaggccgcctgaaggacagggtgcagcctgagatcctggaggccctgctgaagcacatctctttcgataagtttgtgcagatcagcctgaaggccctgagaaggatcgtgccactgatggagcagggcaagcggtacgacgaggcctgcgccgagatctacggcgttcactatggcaagaagaacacagaggagaagatctatctgccccctatccctgccgacgagatcagaaatcctgtggtgctgagggccctgtcccaggcaagaaaagtgatcaacggagtggtgcgccggtacggatctccagcccggatccacatcgagaccgccagagaagtgggcaagagcttcaaggaccggaaggagatcgcgaagagacaggaggagaatcgcaaggatcgggagaaggccgccgccaagtttagggagtacttccctaactttgtgggcgagccaaagtctaaggacatcctgaagctgcgcctgtacgagcagcagcacggcaagtgtctgtatagcggcaaagagatcaatctggtgcggctgaacgagaagggctatgtggagatcgatcacgccctgcctttctccagaacctgggacgattcttttaacaataaggtgctggtgctgggcagcgagaaccagaataagggcaatcagacaccatacgagtatttcaatggcaaggacaactccagggagtggcaggagttcaaggcccgcgtggagacctctagatttcccagtagcaagaagcagcggatcctgctgcagaagttcgacgaggatggctttaaggagtgcaacctgaatgacaccagatacgtgaaccggttcctgtgccagtttgtggccgatcacatcctgctgaccggcaagggcaagagaagggtggtcgcctctaatggccagatcacaaacctgctgagggggttttggagactgaggaaggtgcgggcagagaatgacagacaccacgcactggatgcagtggtggtggcatgcagcaccgtggcaatgcagcagaagatcacaagattcgtgaggtataaggagatgaacgcctttgacggcaagaccgtcgataaggagacaggcaaggtgctgtaccagaagacccacttcccccagccttgggagttctttgcccaggaagttatgatccgggtgttcggcaagccagacggcaagcctgagtttgaggaggccgataccccagagaagctgaggacactgctggcagagaagctgtctagcaggccagaggcagtgcacgagtacgtgaccccgctgttcgtgtccagggcacccaatcggaagatgtctggcgcccacaaggacacactgagaagcgccaagaggtttgtgaagcacaacgagaagatctccgtgaagagagtgtggctgaccgagatcaagctggccgatctggagaacatggtgaattacaagaacggcagggagatcgagctgtatgaggccctgaaggcaaggctggaggcctacggaggaaatgccaagcaggccttcg acccaaaggataaccccttttataagaagggaggacagctggtgaaggccgtgcgggtggagaagacccagaagagcggcgtgctgctgaataagaagaacgcctacacaatcg ccgacaatggtgatatggtgagagtggacgtgttctgtaaggtggataagaagggcaagaatcagtactttatcgtgcctatctatgcctggcaggtggccgagaacatcctgccag acatcgattgcaagggctacagaatcgacgatagctatacattctgtttttccctgcacaagtatgacctgatcgccttccagaaggatgagaagtccaaggtggagtttgcctactat atcaattgcgactcctctagcggcgggttctacctggcctggcacgataagggcagcagggagcagcggtttcgcatctccacccagaatctggcgctgatccagaagtatcaggtg aacgagctgggcaaggagatcaggccatgtcggctgaagaagcgcccacccgtgcggtctggcggctcaaaaagaaccgccgacggcagcgaattcgagcccaagaagaagag gaaagtc

**Nme2-ABE8e-i8_linker10 (WT)**: <u>BPSV40-NLS</u>, Nme2Cas9 – delta PID, <u>TadA8e</u>, <u>SmuCas9 PID</u>, Linkers

MKRTADGSEFESPKKKRKVEDMAAFKPNPINYILGLAIGIASVGWAMVEIDEEENPIRLIDLGVRVFERAEVPKTGDSLAMARRLARSVRRLTRR RAHRLLRARRLLKREGVLQAADFDENGLIKSLPNTPWQLRAAALDRKLTPLEWSAVLLHLIKHRGYLSQRKNEGETADKELGALLKGVANNAHAL QTGDFRTPAELALNKFEKESGHIRNQRGDYSHTFSRKDLQAELILLFEKQKEFGNPHVSGGLKEGIETLLMTQRPALSGDAVQKMLGHCTFEPAE PKAAKNTYTAERFIWLTKLNNLRILEQGSERPLTDTERATLMDEPYRKSKLTYAQARKLLGLEDTAFFKGLRYGKDNAEASTLMEMKAYHAISRAL EKEGLKDKKSPLNLSSELQDEIGTAFSLFKTDEDITGRLKDRVQPEILEALLKHISFDKFVQISLKALRRIVPLMEQGKRYDEACAEIYGDHYGKKNTE EKIYLPPIPADEIRNPVVLRALSQARKVINGVVRRYGSPARIHIETAREVGKSFKDRKEIEKRQEENRKDREKAAAKFREYFPNFVGEPKSKDILKLRL YEQQHGKCLYSGKEINLVRLNEKGYVEIDHALPFSRTWDDSFNNKVLVLGSENQNKGNQTPYEYFNGKDNSREWQEFKARVETSRFPRSKKQRI LLQKFDEDGFKECNLNDTRYVNRFLCQFVADHILLTGKGKRRVFASNGQITNLLRGFWGLRKVRAENDRHHALDAVVVACSTVAMQQKITRFV RYKEMNAFDGKTIDKETGKVLHQKTHFPQPWEFFAQEVMIRVFGKPDGKPSGGSGGSGGSSEVEFSHEYWMRHALTLAKRARDEREVPVGA VLVLNNRVIGEGWNRAIGLHDPTAHAEIMALRQGGLVMQNYRLIDATLYVTFEPCVMCAGAMIHSRIGRVVFGVRNSKRGAAGSLMNVLNYP GMNHRVEITEGILADECAALLCDFYRMPRQVFNAQKKAQSSINETPGTSESATEFEEADTPEKLRTLLAEKLSSRPEAVHEYVTPLFVSRAPNRKM SGAHKDTLRSAKRFVKHNEKISVKRVWLTEIKLADLENMVNYKNGREIELYEALKARLEAYGGNAKQAFDPKDNPFYKKGGQLVKAVRVEKTQE SGVLLNKKNAYTIADNGDMVRVDVFCKVDKKGKNQYFIVPIYAWQVAENILPDIDCKGYRIDDSYTFCFSLHKYDLIAFQKDEKSKVEFAYYINCD SSNGRFYLAWHDKGSKEQQFRISTQNLVLIQKYQVNELGKEIRPCRLKKRPPVREDKRTADGSEFEPKKKRKV

atgaaacggacagccgacggaagcgagttcgagtcaccaaagaagaagcggaaagtcgaagatatggccgccttcaagcctaacccaatcaattacatcctgggactggCcatc ggaatcgcatccgtgggatgggctatggtggagatcgacgaggaggagaatcctatccggctgatcgatctgggcgtgagagtgtttgagagggccgaggtgccaaagaccggcg attctctggctatggcccggagactggcacggagcgtgaggcgcctgacacggagaagggcacacaggctgctgagggcacgccggctgctgaagagagagggcgtgctgcagg cagcagacttcgatgagaatggcctgatcaagagcctgccaaacaccccctggcagctgagagcagccgccctggacaggaagctgacaccactggagtggtctgccgtgctgct gcacctgatcaagcaccgcggctacctgagccagcggaagaacgagggagagacagcagacaaggagctgggcgccctgctgaagggagtggccaacaatgcccacgccctgc agaccggcgatttcaggacacctgccgagctggccctgaataagtttgagaaggagtccggccacatcagaaaccagaggggcgactatagccacaccttctcccgcaaggatctg caggccgagctgatcctgctgttcgagaagcagaaggagtttggcaatccacacgtgagcggaggcctgaaggagggaatcgagaccctgctgatgacacagaggcctgccctgt ccggcgacgcagtgcagaagatgctgggacactgcaccttcgagcctgcagagccaaaggccgccaagaacacctacacagccgagcggtttatctggctgacaaagctgaaca atctgagaatcctggagcagggatccgagaggccactgaccgacacagagagggccaccctgatggatgagccttaccggaagtctaagctgacatatgcccaggccagaaagct gctgggcctggaggacaccgccttctttaagggcctgagatacggcaaggataatgccgaggcctccacactgatggagatgaaggcctatcacgccatctctcgcgccctggaga aggagggcctgaaggacaagaagtcccccctgaacctgagctccgagctgcaggatgagatcggcaccgccttctctctgtttaagaccgacgaggatatcacaggccgcctgaag gacagggtgcagcctgagatcctggaggccctgctgaagcacatctctttcgataagtttgtgcagatcagcctgaaggccctgagaaggatcgtgccactgatggagcagggcaa gcggtacgacgaggcctgcgccgagatctacggcgatcactatggcaagaagaacacagaggagaagatctatctgccccctatccctgccgacgagatcagaaatcctgtggtg ctgagggccctgtcccaggcaagaaaagtgatcaacggagtggtgcgccggtacggatctccagcccggatccacatcgagaccgccagagaagtgggcaagagcttcaaggac cggaaggagatcgagaagagacaggaggagaatcgcaaggatcgggagaaggccgccgccaagtttagggagtacttccctaactttgtgggcgagccaaagtctaaggacatc ctgaagctgcgcctgtacgagcagcagcacggcaagtgtctgtatagcggcaaggagatcaatctggtgcggctgaacgagaagggctatgtggagatcgatcacgccctgccttt ctccagaacctgggacgattcttttaacaataaggtgctggtgctgggcagcgagaaccagaataagggcaatcagacaccatacgagtatttcaatggcaaggacaactccaggg agtggcaggagttcaaggcccgcgtggagacctctagatttcccaggagcaagaagcagcggatcctgctgcagaagttcgacgaggatggctttaaggagtgcaacctgaatga caccagatacgtgaaccggttcctgtgccagtttgtggccgatcacatcctgctgaccggcaagggcaagagaagggtgttcgcctctaatggccagatcacaaacctgctgagggg attttggggactgaggaaggtgcgggcagagaatgacagacaccacgcactggatgcagtggtggtggcatgcagcaccgtggcaatgcagcagaagatcacaagattcgtgag gtataaggagatgaacgcctttgacggcaagaccatcgataaggagacaggcaaggtgctgcaccagaagacccacttcccccagccttgggagttctttgcccaggaagtgatga tccgggtgttcggcaagccagacggcaagccttctggcggttcaggtggatcaggcggtagctctgaggtggagttttcccacgagtactggatgagacatgccctgaccctggcca agagggcacgcgatgagagggaggtgcctgtgggagccgtgctggtgctgaacaatagagtgatcggcgagggctggaacagagccatcggcctgcacgacccaacagcccatg ccgaaattatggccctgagacagggcggcctggtcatgcagaactacagactgattgacgccaccctgtacgtgacattcgagccttgcgtgatgtgcgccggcgccatgatccact ctaggatcggccgcgtggtgtttggcgtgaggaacagcaaacggggcgccgcaggctccctgatgaacgtgctgaactaccccggcatgaatcaccgcgtcgaaattaccgaggg aatcctggcagatgaatgtgccgccctgctgtgcgacttctaccggatgcctagacaggtgttcaatgctcagaagaaggcccagagctccatcaacgagacacctggcacaagcg agagcgcaacagagtttgaggaggccgataccccagagaagctgaggacactgctggcagagaagctgtctagcaggccagaggcagtgcacgagtacgtgaccccactgttcg tgtccagggcacccaatcggaagatgtctggcgcccacaaggacacactgagaagcgccaagaggtttgtgaagcacaacgagaagatctccgtgaagagagtgtggctgaccg agatcaagctggccgatctggagaacatggtgaattacaagaacggcagggagatcgagctgtatgaggccctgaaggcaaggctggaggcctacggaggaaatgccaagcag gccttcgacccaaaggataaccccttttataagaagggaggacagctggtgaaggccgtgcgggtggagaagacccaggagagcggcgtgctgctgaataagaagaacgcctac acaatcgccgacaatggcgatatggtgagagtggacgtgttctgtaaggtggataagaagggcaagaatcagtactttatcgtgcctatctatgcctggcaggtggccgagaacatc ctgccagacatcgattgcaagggctacagaatcgacgatagctatacattctgtttttccctgcacaagtatgacctgatcgccttccagaaggatgagaagtccaaggtggagtttg cctactatatcaattgcgactcctctaacggcaggttctacctggcctggcacgataagggcagcaaggagcagcagtttcgcatctccacccagaatctggtgctgatccagaagta tcaggtgaacgagctgggcaaggagatcaggccatgtcggctgaagaagcgcccacccgtgcgggaggataaaagaaccgccgacggcagcgaattcgagcccaagaagaaga ggaaagtc

**Nme2-ABE8e-i8_linker10 (E932R)**: <u>BPSV40-NLS</u>, Nme2Cas9 – delta PID, <u>TadA8e</u>, <u>SmuCas9 PID</u>, Linkers

MKRTADGSEFESPKKKRKVEDMAAFKPNPINYILGLAIGIASVGWAMVEIDEEENPIRLIDLGVRVFERAEVPKTGDSLAMARRLARSVRRLTRR RAHRLLRARRLLKREGVLQAADFDENGLIKSLPNTPWQLRAAALDRKLTPLEWSAVLLHLIKHRGYLSQRKNEGETADKELGALLKGVANNAHAL QTGDFRTPAELALNKFEKESGHIRNQRGDYSHTFSRKDLQAELILLFEKQKEFGNPHVSGGLKEGIETLLMTQRPALSGDAVQKMLGHCTFEPAE PKAAKNTYTAERFIWLTKLNNLRILEQGSERPLTDTERATLMDEPYRKSKLTYAQARKLLGLEDTAFFKGLRYGKDNAEASTLMEMKAYHAISRAL EKEGLKDKKSPLNLSSELQDEIGTAFSLFKTDEDITGRLKDRVQPEILEALLKHISFDKFVQISLKALRRIVPLMEQGKRYDEACAEIYGDHYGKKNTE EKIYLPPIPADEIRNPVVLRALSQARKVINGVVRRYGSPARIHIETAREVGKSFKDRKEIEKRQEENRKDREKAAAKFREYFPNFVGEPKSKDILKLRL YEQQHGKCLYSGKEINLVRLNEKGYVEIDHALPFSRTWDDSFNNKVLVLGSENQNKGNQTPYEYFNGKDNSREWQEFKARVETSRFPRSKKQRI LLQKFDEDGFKECNLNDTRYVNRFLCQFVADHILLTGKGKRRVFASNGQITNLLRGFWGLRKVRAENDRHHALDAVVVACSTVAMQQKITRFV RYKEMNAFDGKTIDKETGKVLHQKTHFPQPWEFFAQEVMIRVFGKPDGKPSGGSGGSGGSSEVEFSHEYWMRHALTLAKRARDEREVPVGA VLVLNNRVIGEGWNRAIGLHDPTAHAEIMALRQGGLVMQNYRLIDATLYVTFEPCVMCAGAMIHSRIGRVVFGVRNSKRGAAGSLMNVLNYP GMNHRVEITEGILADECAALLCDFYRMPRQVFNAQKKAQSSINETPGTSESATEFEEADTPEKLRTLLAEKLSSRPEAVHEYVTPLFVSRAPNRKM SGAHKDTLRSAKRFVKHNEKISVKRVWLTEIKLADLENMVNYKNGREIELYEALKARLEAYGGNAKQAFDPKDNPFYKKGGQLVKAVRVEKTRE SGVLLNKKNAYTIADNGDMVRVDVFCKVDKKGKNQYFIVPIYAWQVAENILPDIDCKGYRIDDSYTFCFSLHKYDLIAFQKDEKSKVEFAYYINCD SSNGRFYLAWHDKGSKEQQFRISTQNLVLIQKYQVNELGKEIRPCRLKKRPPVREDKRTADGSEFEPKKKRKV

atgaaacggacagccgacggaagcgagttcgagtcaccaaagaagaagcggaaagtcgaagatatggccgccttcaagcctaacccaatcaattacatcctgggactggCcatc ggaatcgcatccgtgggatgggctatggtggagatcgacgaggaggagaatcctatccggctgatcgatctgggcgtgagagtgtttgagagggccgaggtgccaaagaccggcg attctctggctatggcccggagactggcacggagcgtgaggcgcctgacacggagaagggcacacaggctgctgagggcacgccggctgctgaagagagagggcgtgctgcagg cagcagacttcgatgagaatggcctgatcaagagcctgccaaacaccccctggcagctgagagcagccgccctggacaggaagctgacaccactggagtggtctgccgtgctgct gcacctgatcaagcaccgcggctacctgagccagcggaagaacgagggagagacagcagacaaggagctgggcgccctgctgaagggagtggccaacaatgcccacgccctgc agaccggcgatttcaggacacctgccgagctggccctgaataagtttgagaaggagtccggccacatcagaaaccagaggggcgactatagccacaccttctcccgcaaggatctg caggccgagctgatcctgctgttcgagaagcagaaggagtttggcaatccacacgtgagcggaggcctgaaggagggaatcgagaccctgctgatgacacagaggcctgccctgt ccggcgacgcagtgcagaagatgctgggacactgcaccttcgagcctgcagagccaaaggccgccaagaacacctacacagccgagcggtttatctggctgacaaagctgaaca atctgagaatcctggagcagggatccgagaggccactgaccgacacagagagggccaccctgatggatgagccttaccggaagtctaagctgacatatgcccaggccagaaagct gctgggcctggaggacaccgccttctttaagggcctgagatacggcaaggataatgccgaggcctccacactgatggagatgaaggcctatcacgccatctctcgcgccctggaga aggagggcctgaaggacaagaagtcccccctgaacctgagctccgagctgcaggatgagatcggcaccgccttctctctgtttaagaccgacgaggatatcacaggccgcctgaag gacagggtgcagcctgagatcctggaggccctgctgaagcacatctctttcgataagtttgtgcagatcagcctgaaggccctgagaaggatcgtgccactgatggagcagggcaa gcggtacgacgaggcctgcgccgagatctacggcgatcactatggcaagaagaacacagaggagaagatctatctgccccctatccctgccgacgagatcagaaatcctgtggtg ctgagggccctgtcccaggcaagaaaagtgatcaacggagtggtgcgccggtacggatctccagcccggatccacatcgagaccgccagagaagtgggcaagagcttcaaggac cggaaggagatcgagaagagacaggaggagaatcgcaaggatcgggagaaggccgccgccaagtttagggagtacttccctaactttgtgggcgagccaaagtctaaggacatc ctgaagctgcgcctgtacgagcagcagcacggcaagtgtctgtatagcggcaaggagatcaatctggtgcggctgaacgagaagggctatgtggagatcgatcacgccctgccttt ctccagaacctgggacgattcttttaacaataaggtgctggtgctgggcagcgagaaccagaataagggcaatcagacaccatacgagtatttcaatggcaaggacaactccaggg agtggcaggagttcaaggcccgcgtggagacctctagatttcccaggagcaagaagcagcggatcctgctgcagaagttcgacgaggatggctttaaggagtgcaacctgaatga caccagatacgtgaaccggttcctgtgccagtttgtggccgatcacatcctgctgaccggcaagggcaagagaagggtgttcgcctctaatggccagatcacaaacctgctgagggg attttggggactgaggaaggtgcgggcagagaatgacagacaccacgcactggatgcagtggtggtggcatgcagcaccgtggcaatgcagcagaagatcacaagattcgtgag gtataaggagatgaacgcctttgacggcaagaccatcgataaggagacaggcaaggtgctgcaccagaagacccacttcccccagccttgggagttctttgcccaggaagtgatga tccgggtgttcggcaagccagacggcaagccttctggcggttcaggtggatcaggcggtagctctgaggtggagttttcccacgagtactggatgagacatgccctgaccctggcca agagggcacgcgatgagagggaggtgcctgtgggagccgtgctggtgctgaacaatagagtgatcggcgagggctggaacagagccatcggcctgcacgacccaacagcccatg ccgaaattatggccctgagacagggcggcctggtcatgcagaactacagactgattgacgccaccctgtacgtgacattcgagccttgcgtgatgtgcgccggcgccatgatccact ctaggatcggccgcgtggtgtttggcgtgaggaacagcaaacggggcgccgcaggctccctgatgaacgtgctgaactaccccggcatgaatcaccgcgtcgaaattaccgaggg aatcctggcagatgaatgtgccgccctgctgtgcgacttctaccggatgcctagacaggtgttcaatgctcagaagaaggcccagagctccatcaacgagacacctggcacaagcg agagcgcaacagagtttgaggaggccgataccccagagaagctgaggacactgctggcagagaagctgtctagcaggccagaggcagtgcacgagtacgtgaccccactgttcg tgtccagggcacccaatcggaagatgtctggcgcccacaaggacacactgagaagcgccaagaggtttgtgaagcacaacgagaagatctccgtgaagagagtgtggctgaccg agatcaagctggccgatctggagaacatggtgaattacaagaacggcagggagatcgagctgtatgaggccctgaaggcaaggctggaggcctacggaggaaatgccaagcag gccttcgacccaaaggataaccccttttataagaagggaggacagctggtgaaggccgtgcgggtggagaagacccagCGTagcggcgtgctgctgaataagaagaacgcctac acaatcgccgacaatggcgatatggtgagagtggacgtgttctgtaaggtggataagaagggcaagaatcagtactttatcgtgcctatctatgcctggcaggtggccgagaacatc ctgccagacatcgattgcaagggctacagaatcgacgatagctatacattctgtttttccctgcacaagtatgacctgatcgccttccagaaggatgagaagtccaaggtggagtttg cctactatatcaattgcgactcctctaacggcaggttctacctggcctggcacgataagggcagcaaggagcagcagtttcgcatctccacccagaatctggtgctgatccagaagta tcaggtgaacgagctgggcaaggagatcaggccatgtcggctgaagaagcgcccacccgtgcgggaggataaaagaaccgccgacggcagcgaattcgagcccaagaagaaga ggaaagtc

**Nme2-ABE8e-i8_linker10 (D56R)**: <u>BPSV40-NLS</u>, Nme2Cas9 – delta PID, <u>TadA8e</u>, <u>SmuCas9 PID</u>, Linkers

MKRTADGSEFESPKKKRKVEDMAAFKPNPINYILGLAIGIASVGWAMVEIDEEENPIRLIDLGVRVFERAEVPKTGRSLAMARRLARSVRRLTRR RAHRLLRARRLLKREGVLQAADFDENGLIKSLPNTPWQLRAAALDRKLTPLEWSAVLLHLIKHRGYLSQRKNEGETADKELGALLKGVANNAHAL QTGDFRTPAELALNKFEKESGHIRNQRGDYSHTFSRKDLQAELILLFEKQKEFGNPHVSGGLKEGIETLLMTQRPALSGDAVQKMLGHCTFEPAE PKAAKNTYTAERFIWLTKLNNLRILEQGSERPLTDTERATLMDEPYRKSKLTYAQARKLLGLEDTAFFKGLRYGKDNAEASTLMEMKAYHAISRAL EKEGLKDKKSPLNLSSELQDEIGTAFSLFKTDEDITGRLKDRVQPEILEALLKHISFDKFVQISLKALRRIVPLMEQGKRYDEACAEIYGDHYGKKNTE EKIYLPPIPADEIRNPVVLRALSQARKVINGVVRRYGSPARIHIETAREVGKSFKDRKEIEKRQEENRKDREKAAAKFREYFPNFVGEPKSKDILKLRL YEQQHGKCLYSGKEINLVRLNEKGYVEIDHALPFSRTWDDSFNNKVLVLGSENQNKGNQTPYEYFNGKDNSREWQEFKARVETSRFPRSKKQRI LLQKFDEDGFKECNLNDTRYVNRFLCQFVADHILLTGKGKRRVFASNGQITNLLRGFWGLRKVRAENDRHHALDAVVVACSTVAMQQKITRFV RYKEMNAFDGKTIDKETGKVLHQKTHFPQPWEFFAQEVMIRVFGKPDGKPSGGSGGSGGSSEVEFSHEYWMRHALTLAKRARDEREVPVGA VLVLNNRVIGEGWNRAIGLHDPTAHAEIMALRQGGLVMQNYRLIDATLYVTFEPCVMCAGAMIHSRIGRVVFGVRNSKRGAAGSLMNVLNYP GMNHRVEITEGILADECAALLCDFYRMPRQVFNAQKKAQSSINETPGTSESATEFEEADTPEKLRTLLAEKLSSRPEAVHEYVTPLFVSRAPNRKM SGAHKDTLRSAKRFVKHNEKISVKRVWLTEIKLADLENMVNYKNGREIELYEALKARLEAYGGNAKQAFDPKDNPFYKKGGQLVKAVRVEKTQE SGVLLNKKNAYTIADNGDMVRVDVFCKVDKKGKNQYFIVPIYAWQVAENILPDIDCKGYRIDDSYTFCFSLHKYDLIAFQKDEKSKVEFAYYINCD SSNGRFYLAWHDKGSKEQQFRISTQNLVLIQKYQVNELGKEIRPCRLKKRPPVREDKRTADGSEFEPKKKRKV

atgaaacggacagccgacggaagcgagttcgagtcaccaaagaagaagcggaaagtcgaagatatggccgccttcaagcctaacccaatcaattacatcctgggactggCcatc ggaatcgcatccgtgggatgggctatggtggagatcgacgaggaggagaatcctatccggctgatcgatctgggcgtgagagtgtttgagagggccgaggtgccaaagaccggcC GTtctctggctatggcccggagactggcacggagcgtgaggcgcctgacacggagaagggcacacaggctgctgagggcacgccggctgctgaagagagagggcgtgctgcag gcagcagacttcgatgagaatggcctgatcaagagcctgccaaacaccccctggcagctgagagcagccgccctggacaggaagctgacaccactggagtggtctgccgtgctgc tgcacctgatcaagcaccgcggctacctgagccagcggaagaacgagggagagacagcagacaaggagctgggcgccctgctgaagggagtggccaacaatgcccacgccctg cagaccggcgatttcaggacacctgccgagctggccctgaataagtttgagaaggagtccggccacatcagaaaccagaggggcgactatagccacaccttctcccgcaaggatct gcaggccgagctgatcctgctgttcgagaagcagaaggagtttggcaatccacacgtgagcggaggcctgaaggagggaatcgagaccctgctgatgacacagaggcctgccctg tccggcgacgcagtgcagaagatgctgggacactgcaccttcgagcctgcagagccaaaggccgccaagaacacctacacagccgagcggtttatctggctgacaaagctgaaca atctgagaatcctggagcagggatccgagaggccactgaccgacacagagagggccaccctgatggatgagccttaccggaagtctaagctgacatatgcccaggccagaaagct gctgggcctggaggacaccgccttctttaagggcctgagatacggcaaggataatgccgaggcctccacactgatggagatgaaggcctatcacgccatctctcgcgccctggaga aggagggcctgaaggacaagaagtcccccctgaacctgagctccgagctgcaggatgagatcggcaccgccttctctctgtttaagaccgacgaggatatcacaggccgcctgaag gacagggtgcagcctgagatcctggaggccctgctgaagcacatctctttcgataagtttgtgcagatcagcctgaaggccctgagaaggatcgtgccactgatggagcagggcaa gcggtacgacgaggcctgcgccgagatctacggcgatcactatggcaagaagaacacagaggagaagatctatctgccccctatccctgccgacgagatcagaaatcctgtggtg ctgagggccctgtcccaggcaagaaaagtgatcaacggagtggtgcgccggtacggatctccagcccggatccacatcgagaccgccagagaagtgggcaagagcttcaaggac cggaaggagatcgagaagagacaggaggagaatcgcaaggatcgggagaaggccgccgccaagtttagggagtacttccctaactttgtgggcgagccaaagtctaaggacatc ctgaagctgcgcctgtacgagcagcagcacggcaagtgtctgtatagcggcaaggagatcaatctggtgcggctgaacgagaagggctatgtggagatcgatcacgccctgccttt ctccagaacctgggacgattcttttaacaataaggtgctggtgctgggcagcgagaaccagaataagggcaatcagacaccatacgagtatttcaatggcaaggacaactccaggg agtggcaggagttcaaggcccgcgtggagacctctagatttcccaggagcaagaagcagcggatcctgctgcagaagttcgacgaggatggctttaaggagtgcaacctgaatga caccagatacgtgaaccggttcctgtgccagtttgtggccgatcacatcctgctgaccggcaagggcaagagaagggtgttcgcctctaatggccagatcacaaacctgctgagggg attttggggactgaggaaggtgcgggcagagaatgacagacaccacgcactggatgcagtggtggtggcatgcagcaccgtggcaatgcagcagaagatcacaagattcgtgag gtataaggagatgaacgcctttgacggcaagaccatcgataaggagacaggcaaggtgctgcaccagaagacccacttcccccagccttgggagttctttgcccaggaagtgatga tccgggtgttcggcaagccagacggcaagccttctggcggttcaggtggatcaggcggtagctctgaggtggagttttcccacgagtactggatgagacatgccctgaccctggcca agagggcacgcgatgagagggaggtgcctgtgggagccgtgctggtgctgaacaatagagtgatcggcgagggctggaacagagccatcggcctgcacgacccaacagcccatg ccgaaattatggccctgagacagggcggcctggtcatgcagaactacagactgattgacgccaccctgtacgtgacattcgagccttgcgtgatgtgcgccggcgccatgatccact ctaggatcggccgcgtggtgtttggcgtgaggaacagcaaacggggcgccgcaggctccctgatgaacgtgctgaactaccccggcatgaatcaccgcgtcgaaattaccgaggg aatcctggcagatgaatgtgccgccctgctgtgcgacttctaccggatgcctagacaggtgttcaatgctcagaagaaggcccagagctccatcaacgagacacctggcacaagcg agagcgcaacagagtttgaggaggccgataccccagagaagctgaggacactgctggcagagaagctgtctagcaggccagaggcagtgcacgagtacgtgaccccactgttcg tgtccagggcacccaatcggaagatgtctggcgcccacaaggacacactgagaagcgccaagaggtttgtgaagcacaacgagaagatctccgtgaagagagtgtggctgaccg agatcaagctggccgatctggagaacatggtgaattacaagaacggcagggagatcgagctgtatgaggccctgaaggcaaggctggaggcctacggaggaaatgccaagcag gccttcgacccaaaggataaccccttttataagaagggaggacagctggtgaaggccgtgcgggtggagaagacccaggagagcggcgtgctgctgaataagaagaacgcctac acaatcgccgacaatggcgatatggtgagagtggacgtgttctgtaaggtggataagaagggcaagaatcagtactttatcgtgcctatctatgcctggcaggtggccgagaacatc ctgccagacatcgattgcaagggctacagaatcgacgatagctatacattctgtttttccctgcacaagtatgacctgatcgccttccagaaggatgagaagtccaaggtggagtttg cctactatatcaattgcgactcctctaacggcaggttctacctggcctggcacgataagggcagcaaggagcagcagtttcgcatctccacccagaatctggtgctgatccagaagta tcaggtgaacgagctgggcaaggagatcaggccatgtcggctgaagaagcgcccacccgtgcgggaggataaaagaaccgccgacggcagcgaattcgagcccaagaagaaga ggaaagtc

**Nme2^Smu^-ABE8e-i8_linker10 (WT)**: <u>BPSV40-NLS</u>, Nme2Cas9 – delta PID, <u>TadA8e</u>, <u>SmuCas9 PID</u>, Linkers

MKRTADGSEFESPKKKRKVEDMAAFKPNPINYILGLAIGIASVGWAMVEIDEEENPIRLIDLGVRVFERAEVPKTGDSLAMARRLARSVRRLTRR RAHRLLRARRLLKREGVLQAADFDENGLIKSLPNTPWQLRAAALDRKLTPLEWSAVLLHLIKHRGYLSQRKNEGETADKELGALLKGVANNAHAL QTGDFRTPAELALNKFEKESGHIRNQRGDYSHTFSRKDLQAELILLFEKQKEFGNPHVSGGLKEGIETLLMTQRPALSGDAVQKMLGHCTFEPAE PKAAKNTYTAERFIWLTKLNNLRILEQGSERPLTDTERATLMDEPYRKSKLTYAQARKLLGLEDTAFFKGLRYGKDNAEASTLMEMKAYHAISRAL EKEGLKDKKSPLNLSSELQDEIGTAFSLFKTDEDITGRLKDRVQPEILEALLKHISFDKFVQISLKALRRIVPLMEQGKRYDEACAEIYGDHYGKKNTE EKIYLPPIPADEIRNPVVLRALSQARKVINGVVRRYGSPARIHIETAREVGKSFKDRKEIEKRQEENRKDREKAAAKFREYFPNFVGEPKSKDILKLRL YEQQHGKCLYSGKEINLVRLNEKGYVEIDHALPFSRTWDDSFNNKVLVLGSENQNKGNQTPYEYFNGKDNSREWQEFKARVETSRFPRSKKQRI LLQKFDEDGFKECNLNDTRYVNRFLCQFVADHILLTGKGKRRVFASNGQITNLLRGFWGLRKVRAENDRHHALDAVVVACSTVAMQQKITRFV RYKEMNAFDGKTIDKETGKVLHQKTHFPQPWEFFAQEVMIRVFGKPDGKPSGGSGGSGGSSEVEFSHEYWMRHALTLAKRARDEREVPVGA VLVLNNRVIGEGWNRAIGLHDPTAHAEIMALRQGGLVMQNYRLIDATLYVTFEPCVMCAGAMIHSRIGRVVFGVRNSKRGAAGSLMNVLNYP GMNHRVEITEGILADECAALLCDFYRMPRQVFNAQKKAQSSINETPGTSESATEFEEADTPEKLRTLLAEKLSSRPEAVHEYVTPLFVSRAPNRKM SGAHKDTLRSAKRFVKHNEKISVKRVWLTEIKLADLENMVNYKNGREIELYEALKARLEAYGGNAKQAFDPKDNPFYKKGGQLVKAVRVEKTQE SGVLLNKKNAYTIADNATMVRVDVYTKAGKNYLVPVYVWQVAQGILPNRAVTSGKSEADWDLIDESFEFKFSLSRGDLVEMISNKGRIFGYYNG LDRANGSIGIREHDLEKSKGKDGVHRVGVKTATAFNKYHVDPLGKEIHRCSSEPRPTLKIKSKKEDKRTADGSEFEPKKKRKV

atgaaacggacagccgacggaagcgagttcgagtcaccaaagaagaagcggaaagtcgaagatatggccgccttcaagcctaacccaatcaattacatcctgggactggCcatc ggaatcgcatccgtgggatgggctatggtggagatcgacgaggaggagaatcctatccggctgatcgatctgggcgtgagagtgtttgagagggccgaggtgccaaagaccggcg attctctggctatggcccggagactggcacggagcgtgaggcgcctgacacggagaagggcacacaggctgctgagggcacgccggctgctgaagagagagggcgtgctgcagg cagcagacttcgatgagaatggcctgatcaagagcctgccaaacaccccctggcagctgagagcagccgccctggacaggaagctgacaccactggagtggtctgccgtgctgct gcacctgatcaagcaccgcggctacctgagccagcggaagaacgagggagagacagcagacaaggagctgggcgccctgctgaagggagtggccaacaatgcccacgccctgc agaccggcgatttcaggacacctgccgagctggccctgaataagtttgagaaggagtccggccacatcagaaaccagaggggcgactatagccacaccttctcccgcaaggatctg caggccgagctgatcctgctgttcgagaagcagaaggagtttggcaatccacacgtgagcggaggcctgaaggagggaatcgagaccctgctgatgacacagaggcctgccctgt ccggcgacgcagtgcagaagatgctgggacactgcaccttcgagcctgcagagccaaaggccgccaagaacacctacacagccgagcggtttatctggctgacaaagctgaaca atctgagaatcctggagcagggatccgagaggccactgaccgacacagagagggccaccctgatggatgagccttaccggaagtctaagctgacatatgcccaggccagaaagct gctgggcctggaggacaccgccttctttaagggcctgagatacggcaaggataatgccgaggcctccacactgatggagatgaaggcctatcacgccatctctcgcgccctggaga aggagggcctgaaggacaagaagtcccccctgaacctgagctccgagctgcaggatgagatcggcaccgccttctctctgtttaagaccgacgaggatatcacaggccgcctgaag gacagggtgcagcctgagatcctggaggccctgctgaagcacatctctttcgataagtttgtgcagatcagcctgaaggccctgagaaggatcgtgccactgatggagcagggcaa gcggtacgacgaggcctgcgccgagatctacggcgatcactatggcaagaagaacacagaggagaagatctatctgccccctatccctgccgacgagatcagaaatcctgtggtg ctgagggccctgtcccaggcaagaaaagtgatcaacggagtggtgcgccggtacggatctccagcccggatccacatcgagaccgccagagaagtgggcaagagcttcaaggac cggaaggagatcgagaagagacaggaggagaatcgcaaggatcgggagaaggccgccgccaagtttagggagtacttccctaactttgtgggcgagccaaagtctaaggacatc ctgaagctgcgcctgtacgagcagcagcacggcaagtgtctgtatagcggcaaggagatcaatctggtgcggctgaacgagaagggctatgtggagatcgatcacgccctgccttt ctccagaacctgggacgattcttttaacaataaggtgctggtgctgggcagcgagaaccagaataagggcaatcagacaccatacgagtatttcaatggcaaggacaactccaggg agtggcaggagttcaaggcccgcgtggagacctctagatttcccaggagcaagaagcagcggatcctgctgcagaagttcgacgaggatggctttaaggagtgcaacctgaatga caccagatacgtgaaccggttcctgtgccagtttgtggccgatcacatcctgctgaccggcaagggcaagagaagggtgttcgcctctaatggccagatcacaaacctgctgagggg attttggggactgaggaaggtgcgggcagagaatgacagacaccacgcactggatgcagtggtggtggcatgcagcaccgtggcaatgcagcagaagatcacaagattcgtgag gtataaggagatgaacgcctttgacggcaagaccatcgataaggagacaggcaaggtgctgcaccagaagacccacttcccccagccttgggagttctttgcccaggaagtgatga tccgggtgttcggcaagccagacggcaagccttctggcggttcaggtggatcaggcggtagctctgaggtggagttttcccacgagtactggatgagacatgccctgaccctggcca agagggcacgcgatgagagggaggtgcctgtgggagccgtgctggtgctgaacaatagagtgatcggcgagggctggaacagagccatcggcctgcacgacccaacagcccatg ccgaaattatggccctgagacagggcggcctggtcatgcagaactacagactgattgacgccaccctgtacgtgacattcgagccttgcgtgatgtgcgccggcgccatgatccact ctaggatcggccgcgtggtgtttggcgtgaggaacagcaaacggggcgccgcaggctccctgatgaacgtgctgaactaccccggcatgaatcaccgcgtcgaaattaccgaggg aatcctggcagatgaatgtgccgccctgctgtgcgacttctaccggatgcctagacaggtgttcaatgctcagaagaaggcccagagctccatcaacgagacacctggcacaagcg agagcgcaacagagtttgaggaggccgataccccagagaagctgaggacactgctggcagagaagctgtctagcaggccagaggcagtgcacgagtacgtgaccccactgttcg tgtccagggcacccaatcggaagatgtctggcgcccacaaggacacactgagaagcgccaagaggtttgtgaagcacaacgagaagatctccgtgaagagagtgtggctgaccg agatcaagctggccgatctggagaacatggtgaattacaagaacggcagggagatcgagctgtatgaggccctgaaggcaaggctggaggcctacggaggaaatgccaagcag gccttcgacccaaaggataaccccttttataagaagggaggacagctggtgaaggccgtgcgggtggagaagacccaggagagcggcgtgctgctgaataagaagaacgcctac acaatcgccgacaacgccaccatggtgcgggtggacgtgtacaccaaggccggcaagaactacctggttcctgtgtacgtgtggcaggtggcccagggcatcttacccaaccgcgc cgtgaccagcggcaagtccgaggctgactgggacctgatcgatgagagcttcgagttcaagttctctctgtcccggggagatctcgtggaaatgatctccaacaagggcagaatctt cggctactacaacggcctggacagagccaacggctctattggaattagagagcacgacctagagaagagcaagggcaaagacggcgtgcatagagtgggagtgaaaacagcta cagcatttaacaagtaccacgtggatcccctgggcaaagagatccacagatgcagcagcgaacccagacctacactgaaaatcaagtctaagaaggaggataaaagaaccgccg acggcagcgaattcgagcccaagaagaagaggaaagtc

**Nme2^Smu^-ABE8e-i8_linker10 (E932R)**: <u>BPSV40-NLS</u>, Nme2Cas9 – delta PID, <u>TadA8e</u>, <u>SmuCas9 PID</u>, Linkers

MKRTADGSEFESPKKKRKVEDMAAFKPNPINYILGLAIGIASVGWAMVEIDEEENPIRLIDLGVRVFERAEVPKTGDSLAMARRLARSVRRLTRR RAHRLLRARRLLKREGVLQAADFDENGLIKSLPNTPWQLRAAALDRKLTPLEWSAVLLHLIKHRGYLSQRKNEGETADKELGALLKGVANNAHAL QTGDFRTPAELALNKFEKESGHIRNQRGDYSHTFSRKDLQAELILLFEKQKEFGNPHVSGGLKEGIETLLMTQRPALSGDAVQKMLGHCTFEPAE PKAAKNTYTAERFIWLTKLNNLRILEQGSERPLTDTERATLMDEPYRKSKLTYAQARKLLGLEDTAFFKGLRYGKDNAEASTLMEMKAYHAISRAL EKEGLKDKKSPLNLSSELQDEIGTAFSLFKTDEDITGRLKDRVQPEILEALLKHISFDKFVQISLKALRRIVPLMEQGKRYDEACAEIYGDHYGKKNTE EKIYLPPIPADEIRNPVVLRALSQARKVINGVVRRYGSPARIHIETAREVGKSFKDRKEIEKRQEENRKDREKAAAKFREYFPNFVGEPKSKDILKLRL YEQQHGKCLYSGKEINLVRLNEKGYVEIDHALPFSRTWDDSFNNKVLVLGSENQNKGNQTPYEYFNGKDNSREWQEFKARVETSRFPRSKKQRI LLQKFDEDGFKECNLNDTRYVNRFLCQFVADHILLTGKGKRRVFASNGQITNLLRGFWGLRKVRAENDRHHALDAVVVACSTVAMQQKITRFV RYKEMNAFDGKTIDKETGKVLHQKTHFPQPWEFFAQEVMIRVFGKPDGKPSGGSGGSGGSSEVEFSHEYWMRHALTLAKRARDEREVPVGA VLVLNNRVIGEGWNRAIGLHDPTAHAEIMALRQGGLVMQNYRLIDATLYVTFEPCVMCAGAMIHSRIGRVVFGVRNSKRGAAGSLMNVLNYP GMNHRVEITEGILADECAALLCDFYRMPRQVFNAQKKAQSSINETPGTSESATEFEEADTPEKLRTLLAEKLSSRPEAVHEYVTPLFVSRAPNRKM SGAHKDTLRSAKRFVKHNEKISVKRVWLTEIKLADLENMVNYKNGREIELYEALKARLEAYGGNAKQAFDPKDNPFYKKGGQLVKAVRVEKTQR SGVLLNKKNAYTIADNATMVRVDVYTKAGKNYLVPVYVWQVAQGILPNRAVTSGKSEADWDLIDESFEFKFSLSRGDLVEMISNKGRIFGYYNG LDRANGSIGIREHDLEKSKGKDGVHRVGVKTATAFNKYHVDPLGKEIHRCSSEPRPTLKIKSKKEDKRTADGSEFEPKKKRKV

atgaaacggacagccgacggaagcgagttcgagtcaccaaagaagaagcggaaagtcgaagatatggccgccttcaagcctaacccaatcaattacatcctgggactggCcatc ggaatcgcatccgtgggatgggctatggtggagatcgacgaggaggagaatcctatccggctgatcgatctgggcgtgagagtgtttgagagggccgaggtgccaaagaccggcg attctctggctatggcccggagactggcacggagcgtgaggcgcctgacacggagaagggcacacaggctgctgagggcacgccggctgctgaagagagagggcgtgctgcagg cagcagacttcgatgagaatggcctgatcaagagcctgccaaacaccccctggcagctgagagcagccgccctggacaggaagctgacaccactggagtggtctgccgtgctgct gcacctgatcaagcaccgcggctacctgagccagcggaagaacgagggagagacagcagacaaggagctgggcgccctgctgaagggagtggccaacaatgcccacgccctgc agaccggcgatttcaggacacctgccgagctggccctgaataagtttgagaaggagtccggccacatcagaaaccagaggggcgactatagccacaccttctcccgcaaggatctg caggccgagctgatcctgctgttcgagaagcagaaggagtttggcaatccacacgtgagcggaggcctgaaggagggaatcgagaccctgctgatgacacagaggcctgccctgt ccggcgacgcagtgcagaagatgctgggacactgcaccttcgagcctgcagagccaaaggccgccaagaacacctacacagccgagcggtttatctggctgacaaagctgaaca atctgagaatcctggagcagggatccgagaggccactgaccgacacagagagggccaccctgatggatgagccttaccggaagtctaagctgacatatgcccaggccagaaagct gctgggcctggaggacaccgccttctttaagggcctgagatacggcaaggataatgccgaggcctccacactgatggagatgaaggcctatcacgccatctctcgcgccctggaga aggagggcctgaaggacaagaagtcccccctgaacctgagctccgagctgcaggatgagatcggcaccgccttctctctgtttaagaccgacgaggatatcacaggccgcctgaag gacagggtgcagcctgagatcctggaggccctgctgaagcacatctctttcgataagtttgtgcagatcagcctgaaggccctgagaaggatcgtgccactgatggagcagggcaa gcggtacgacgaggcctgcgccgagatctacggcgatcactatggcaagaagaacacagaggagaagatctatctgccccctatccctgccgacgagatcagaaatcctgtggtg ctgagggccctgtcccaggcaagaaaagtgatcaacggagtggtgcgccggtacggatctccagcccggatccacatcgagaccgccagagaagtgggcaagagcttcaaggac cggaaggagatcgagaagagacaggaggagaatcgcaaggatcgggagaaggccgccgccaagtttagggagtacttccctaactttgtgggcgagccaaagtctaaggacatc ctgaagctgcgcctgtacgagcagcagcacggcaagtgtctgtatagcggcaaggagatcaatctggtgcggctgaacgagaagggctatgtggagatcgatcacgccctgccttt ctccagaacctgggacgattcttttaacaataaggtgctggtgctgggcagcgagaaccagaataagggcaatcagacaccatacgagtatttcaatggcaaggacaactccaggg agtggcaggagttcaaggcccgcgtggagacctctagatttcccaggagcaagaagcagcggatcctgctgcagaagttcgacgaggatggctttaaggagtgcaacctgaatga caccagatacgtgaaccggttcctgtgccagtttgtggccgatcacatcctgctgaccggcaagggcaagagaagggtgttcgcctctaatggccagatcacaaacctgctgagggg attttggggactgaggaaggtgcgggcagagaatgacagacaccacgcactggatgcagtggtggtggcatgcagcaccgtggcaatgcagcagaagatcacaagattcgtgag gtataaggagatgaacgcctttgacggcaagaccatcgataaggagacaggcaaggtgctgcaccagaagacccacttcccccagccttgggagttctttgcccaggaagtgatga tccgggtgttcggcaagccagacggcaagccttctggcggttcaggtggatcaggcggtagctctgaggtggagttttcccacgagtactggatgagacatgccctgaccctggcca agagggcacgcgatgagagggaggtgcctgtgggagccgtgctggtgctgaacaatagagtgatcggcgagggctggaacagagccatcggcctgcacgacccaacagcccatg ccgaaattatggccctgagacagggcggcctggtcatgcagaactacagactgattgacgccaccctgtacgtgacattcgagccttgcgtgatgtgcgccggcgccatgatccact ctaggatcggccgcgtggtgtttggcgtgaggaacagcaaacggggcgccgcaggctccctgatgaacgtgctgaactaccccggcatgaatcaccgcgtcgaaattaccgaggg aatcctggcagatgaatgtgccgccctgctgtgcgacttctaccggatgcctagacaggtgttcaatgctcagaagaaggcccagagctccatcaacgagacacctggcacaagcg agagcgcaacagagtttgaggaggccgataccccagagaagctgaggacactgctggcagagaagctgtctagcaggccagaggcagtgcacgagtacgtgaccccactgttcg tgtccagggcacccaatcggaagatgtctggcgcccacaaggacacactgagaagcgccaagaggtttgtgaagcacaacgagaagatctccgtgaagagagtgtggctgaccg agatcaagctggccgatctggagaacatggtgaattacaagaacggcagggagatcgagctgtatgaggccctgaaggcaaggctggaggcctacggaggaaatgccaagcag gccttcgacccaaaggataaccccttttataagaagggaggacagctggtgaaggccgtgcgggtggagaagacccagCGTagcggcgtgctgctgaataagaagaacgcctac acaatcgccgacaacgccaccatggtgcgggtggacgtgtacaccaaggccggcaagaactacctggttcctgtgtacgtgtggcaggtggcccagggcatcttacccaaccgcgc cgtgaccagcggcaagtccgaggctgactgggacctgatcgatgagagcttcgagttcaagttctctctgtcccggggagatctcgtggaaatgatctccaacaagggcagaatctt cggctactacaacggcctggacagagccaacggctctattggaattagagagcacgacctagagaagagcaagggcaaagacggcgtgcatagagtgggagtgaaaacagcta cagcatttaacaagtaccacgtggatcccctgggcaaagagatccacagatgcagcagcgaacccagacctacactgaaaatcaagtctaagaaggaggataaaagaaccgccg acggcagcgaattcgagcccaagaagaagaggaaagtc

**Nme2^Smu^-ABE8e-i8_linker10 (D56R)**: <u>BPSV40-NLS</u>, Nme2Cas9 – delta PID, <u>TadA8e</u>, <u>SmuCas9 PID</u>, Linkers

MKRTADGSEFESPKKKRKVEDMAAFKPNPINYILGLAIGIASVGWAMVEIDEEENPIRLIDLGVRVFERAEVPKTGRSLAMARRLARSVRRLTRR RAHRLLRARRLLKREGVLQAADFDENGLIKSLPNTPWQLRAAALDRKLTPLEWSAVLLHLIKHRGYLSQRKNEGETADKELGALLKGVANNAHAL QTGDFRTPAELALNKFEKESGHIRNQRGDYSHTFSRKDLQAELILLFEKQKEFGNPHVSGGLKEGIETLLMTQRPALSGDAVQKMLGHCTFEPAE PKAAKNTYTAERFIWLTKLNNLRILEQGSERPLTDTERATLMDEPYRKSKLTYAQARKLLGLEDTAFFKGLRYGKDNAEASTLMEMKAYHAISRAL EKEGLKDKKSPLNLSSELQDEIGTAFSLFKTDEDITGRLKDRVQPEILEALLKHISFDKFVQISLKALRRIVPLMEQGKRYDEACAEIYGDHYGKKNTE EKIYLPPIPADEIRNPVVLRALSQARKVINGVVRRYGSPARIHIETAREVGKSFKDRKEIEKRQEENRKDREKAAAKFREYFPNFVGEPKSKDILKLRL YEQQHGKCLYSGKEINLVRLNEKGYVEIDHALPFSRTWDDSFNNKVLVLGSENQNKGNQTPYEYFNGKDNSREWQEFKARVETSRFPRSKKQRI LLQKFDEDGFKECNLNDTRYVNRFLCQFVADHILLTGKGKRRVFASNGQITNLLRGFWGLRKVRAENDRHHALDAVVVACSTVAMQQKITRFV RYKEMNAFDGKTIDKETGKVLHQKTHFPQPWEFFAQEVMIRVFGKPDGKPSGGSGGSGGSSEVEFSHEYWMRHALTLAKRARDEREVPVGA VLVLNNRVIGEGWNRAIGLHDPTAHAEIMALRQGGLVMQNYRLIDATLYVTFEPCVMCAGAMIHSRIGRVVFGVRNSKRGAAGSLMNVLNYP GMNHRVEITEGILADECAALLCDFYRMPRQVFNAQKKAQSSINETPGTSESATEFEEADTPEKLRTLLAEKLSSRPEAVHEYVTPLFVSRAPNRKM SGAHKDTLRSAKRFVKHNEKISVKRVWLTEIKLADLENMVNYKNGREIELYEALKARLEAYGGNAKQAFDPKDNPFYKKGGQLVKAVRVEKTQE SGVLLNKKNAYTIADNATMVRVDVYTKAGKNYLVPVYVWQVAQGILPNRAVTSGKSEADWDLIDESFEFKFSLSRGDLVEMISNKGRIFGYYNG LDRANGSIGIREHDLEKSKGKDGVHRVGVKTATAFNKYHVDPLGKEIHRCSSEPRPTLKIKSKKEDKRTADGSEFEPKKKRKV

atgaaacggacagccgacggaagcgagttcgagtcaccaaagaagaagcggaaagtcgaagatatggccgccttcaagcctaacccaatcaattacatcctgggactggCcatc ggaatcgcatccgtgggatgggctatggtggagatcgacgaggaggagaatcctatccggctgatcgatctgggcgtgagagtgtttgagagggccgaggtgccaaagaccggcC GTtctctggctatggcccggagactggcacggagcgtgaggcgcctgacacggagaagggcacacaggctgctgagggcacgccggctgctgaagagagagggcgtgctgcag gcagcagacttcgatgagaatggcctgatcaagagcctgccaaacaccccctggcagctgagagcagccgccctggacaggaagctgacaccactggagtggtctgccgtgctgc tgcacctgatcaagcaccgcggctacctgagccagcggaagaacgagggagagacagcagacaaggagctgggcgccctgctgaagggagtggccaacaatgcccacgccctg cagaccggcgatttcaggacacctgccgagctggccctgaataagtttgagaaggagtccggccacatcagaaaccagaggggcgactatagccacaccttctcccgcaaggatct gcaggccgagctgatcctgctgttcgagaagcagaaggagtttggcaatccacacgtgagcggaggcctgaaggagggaatcgagaccctgctgatgacacagaggcctgccctg tccggcgacgcagtgcagaagatgctgggacactgcaccttcgagcctgcagagccaaaggccgccaagaacacctacacagccgagcggtttatctggctgacaaagctgaaca atctgagaatcctggagcagggatccgagaggccactgaccgacacagagagggccaccctgatggatgagccttaccggaagtctaagctgacatatgcccaggccagaaagct gctgggcctggaggacaccgccttctttaagggcctgagatacggcaaggataatgccgaggcctccacactgatggagatgaaggcctatcacgccatctctcgcgccctggaga aggagggcctgaaggacaagaagtcccccctgaacctgagctccgagctgcaggatgagatcggcaccgccttctctctgtttaagaccgacgaggatatcacaggccgcctgaag gacagggtgcagcctgagatcctggaggccctgctgaagcacatctctttcgataagtttgtgcagatcagcctgaaggccctgagaaggatcgtgccactgatggagcagggcaa gcggtacgacgaggcctgcgccgagatctacggcgatcactatggcaagaagaacacagaggagaagatctatctgccccctatccctgccgacgagatcagaaatcctgtggtg ctgagggccctgtcccaggcaagaaaagtgatcaacggagtggtgcgccggtacggatctccagcccggatccacatcgagaccgccagagaagtgggcaagagcttcaaggac cggaaggagatcgagaagagacaggaggagaatcgcaaggatcgggagaaggccgccgccaagtttagggagtacttccctaactttgtgggcgagccaaagtctaaggacatc ctgaagctgcgcctgtacgagcagcagcacggcaagtgtctgtatagcggcaaggagatcaatctggtgcggctgaacgagaagggctatgtggagatcgatcacgccctgccttt ctccagaacctgggacgattcttttaacaataaggtgctggtgctgggcagcgagaaccagaataagggcaatcagacaccatacgagtatttcaatggcaaggacaactccaggg agtggcaggagttcaaggcccgcgtggagacctctagatttcccaggagcaagaagcagcggatcctgctgcagaagttcgacgaggatggctttaaggagtgcaacctgaatga caccagatacgtgaaccggttcctgtgccagtttgtggccgatcacatcctgctgaccggcaagggcaagagaagggtgttcgcctctaatggccagatcacaaacctgctgagggg attttggggactgaggaaggtgcgggcagagaatgacagacaccacgcactggatgcagtggtggtggcatgcagcaccgtggcaatgcagcagaagatcacaagattcgtgag gtataaggagatgaacgcctttgacggcaagaccatcgataaggagacaggcaaggtgctgcaccagaagacccacttcccccagccttgggagttctttgcccaggaagtgatga tccgggtgttcggcaagccagacggcaagccttctggcggttcaggtggatcaggcggtagctctgaggtggagttttcccacgagtactggatgagacatgccctgaccctggcca agagggcacgcgatgagagggaggtgcctgtgggagccgtgctggtgctgaacaatagagtgatcggcgagggctggaacagagccatcggcctgcacgacccaacagcccatg ccgaaattatggccctgagacagggcggcctggtcatgcagaactacagactgattgacgccaccctgtacgtgacattcgagccttgcgtgatgtgcgccggcgccatgatccact ctaggatcggccgcgtggtgtttggcgtgaggaacagcaaacggggcgccgcaggctccctgatgaacgtgctgaactaccccggcatgaatcaccgcgtcgaaattaccgaggg aatcctggcagatgaatgtgccgccctgctgtgcgacttctaccggatgcctagacaggtgttcaatgctcagaagaaggcccagagctccatcaacgagacacctggcacaagcg agagcgcaacagagtttgaggaggccgataccccagagaagctgaggacactgctggcagagaagctgtctagcaggccagaggcagtgcacgagtacgtgaccccactgttcg tgtccagggcacccaatcggaagatgtctggcgcccacaaggacacactgagaagcgccaagaggtttgtgaagcacaacgagaagatctccgtgaagagagtgtggctgaccg agatcaagctggccgatctggagaacatggtgaattacaagaacggcagggagatcgagctgtatgaggccctgaaggcaaggctggaggcctacggaggaaatgccaagcag gccttcgacccaaaggataaccccttttataagaagggaggacagctggtgaaggccgtgcgggtggagaagacccaggagagcggcgtgctgctgaataagaagaacgcctac acaatcgccgacaacgccaccatggtgcgggtggacgtgtacaccaaggccggcaagaactacctggttcctgtgtacgtgtggcaggtggcccagggcatcttacccaaccgcgc cgtgaccagcggcaagtccgaggctgactgggacctgatcgatgagagcttcgagttcaagttctctctgtcccggggagatctcgtggaaatgatctccaacaagggcagaatctt cggctactacaacggcctggacagagccaacggctctattggaattagagagcacgacctagagaagagcaagggcaaagacggcgtgcatagagtgggagtgaaaacagcta cagcatttaacaagtaccacgtggatcccctgggcaaagagatccacagatgcagcagcgaacccagacctacactgaaaatcaagtctaagaaggaggataaaagaaccgccg acggcagcgaattcgagcccaagaagaagaggaaagtc

**iNme2-ABE8e-nt**: <u>BPSV40-NLS</u>, nNme2Cas9, <u>TadA8e</u>, Linkers

Jenifer Dounda Lab, iNme2Cas9 (D16A) variant in n-terminally fused ABE8e format.

MKRTADGSEFESPKKKRKVGGSGGGSGGGSGSEVEFSHEYWMRHALTLAKRARDEREVPVGAVLVLNNRVIGEGWNRAIGLHDPTAHAEIM ALRQGGLVMQNYRLIDATLYVTFEPCVMCAGAMIHSRIGRVVFGVRNSKRGAAGSLMNVLNYPGMNHRVEITEGILADECAALLCDFYRMPR QVFNAQKKAQSSINSGGSSGGSSGSETPGTSESATPESSGGSSGGSMAAFKPNPINYILGLAIGIASVGWAMVEIDEEENPIRLIDLGVRVFERAE VPKTGDSLAMARRLARSVRRLTRRRAHRLLRARRLLKREGVLQAADFDENGLIKSLPNTPWQLRAAALDRKLTPLEWSAVLLHLIKHRGYLSQRK NEGETADKELGALLKGVANNAHALQTGDFRTPAELALNKFEKESGHIRNQRGDYSHTFSRKDLQAELILLFEKQKEFGNPHVSGGLKEGIETLLM TQRPALSGDAVQKMLGHCTFEPAEPKAAKNTYTAERFIWLTKLNNLRILEQGSERPLTDTERATLMDEPYRKSKLTYAQARKLLGLEDTAFFKGL RYGKDNAEASTLMEMKAYHAISRALEKEGLKDKKSPLNLSSELQDEIGTAFSLFKTDEDITGRLKDRVQPEILEALLKHISFDKFVQISLKALRRIVPL MEQGKRYDEACAEIYGDHYGKKNTEEKIYLPPIPADEIRNPVVLRALSQARKVINGVVRRYGSPARIHIETAREVGKSFKDRKEIEKRQEENRKDRE KAAAKFREYFPNFVGEPKSKDILKLRLYEQQHGKCLYSGKEINLVRLNEKGYVEIDHALPFSRTWDDSFNNKVLVLGSENQNKGNQTPYEYFNGK DNSREWQEFKARVETSRFPRSKKQRILLQKFDEDGFKECNLNDTRYVNRFLCQFVADHILLTGKGKRRVFASNGQITNLLRGFWGLRKVRAEND RHHALDAVVVACSTVAMQQKITRFVRYKEMNAFDGKTIDKETGKVLHQKTHFPQPWEFFAQEVMIRVFGKPDGKPEFEEADTPEKLRTLLAEK LSSRPEAVHEYVTPLFVSRAPNRKMSGAHKGTLRSAKRFVKHNEKISVKRVWLTKIRLAALENMVNYKNGREIELYEALKARLEAYGGNAKQAFD PKGNPFYKKGGQLVKAVRVERTQKSGVLLNKKNAYTIADNGDMVRVDVFCKVDKKGKNQYFIVPIYAWQVAENILPDIDCKGYRIDDSYTFCFS LHKYDLIAFQKDEKSKVEFAYYINCDSSNGRFYLAWHDKGSKEQQFRISTQNLVLIQKYQVNELGKEIRPCRLKKRPPVREDKRTADGSEFEPKKK RKV

atgaaacggacagccgacggaagcgagttcgagtcaccaaagaagaagcggaaagtcggcggtagcggcggaggcagcggtggcggcagcggctctgaggtggagttttccca cgagtactggatgagacatgccctgaccctggccaagagggcacgcgatgagagggaggtgcctgtgggagccgtgctggtgctgaacaatagagtgatcggcgagggctggaa cagagccatcggcctgcacgacccaacagcccatgccgaaattatggccctgagacagggcggcctggtcatgcagaactacagactgattgacgccaccctgtacgtgacattcg agccttgcgtgatgtgcgccggcgccatgatccactctaggatcggccgcgtggtgtttggcgtgaggaacagcaaacggggcgccgcaggctccctgatgaacgtgctgaactac cccggcatgaatcaccgcgtcgaaattaccgagggaatcctggcagatgaatgtgccgccctgctgtgcgacttctaccggatgcctagacaggtgttcaatgctcagaagaaggcc cagagctccatcaactccggaggatctagcggaggctcctctggctctgagacacctggcacaagcgagagcgcaacacctgaaagcagcgggggcagcagcggggggtcaatg gccgccttcaagcctaacccaatcaattacatcctgggactggccatcggaatcgcatccgtgggatgggctatggtggagatcgacgaggaggagaatcctatccggctgatcgat ctgggcgtgagagtgtttgagagggccgaggtgccaaagaccggcgattctctggctatggcccggagactggcacggagcgtgaggcgcctgacacggagaagggcacacagg ctgctgagggcacgccggctgctgaagagagagggcgtgctgcaggcagcagacttcgatgagaatggcctgatcaagagcctgccaaacaccccctggcagctgagagcagcc gccctggacaggaagctgacaccactggagtggtctgccgtgctgctgcacctgatcaagcaccgcggctacctgagccagcggaagaacgagggagagacagcagacaaggag ctgggcgccctgctgaagggagtggccaacaatgcccacgccctgcagaccggcgatttcaggacacctgccgagctggccctgaataagtttgagaaggagtccggccacatca gaaaccagaggggcgactatagccacaccttctcccgcaaggatctgcaggccgagctgatcctgctgttcgagaagcagaaggagtttggcaatccacacgtgagcggaggcct gaaggagggaatcgagaccctgctgatgacacagaggcctgccctgtccggcgacgcagtgcagaagatgctgggacactgcaccttcgagcctgcagagccaaaggccgccaa gaacacctacacagccgagcggtttatctggctgacaaagctgaacaatctgagaatcctggagcagggatccgagaggccactgaccgacacagagagggccaccctgatggat gagccttaccggaagtctaagctgacatatgcccaggccagaaagctgctgggcctggaggacaccgccttctttaagggcctgagatacggcaaggataatgccgaggcctccac actgatggagatgaaggcctatcacgccatctctcgcgccctggagaaggagggcctgaaggacaagaagtcccccctgaacctgagctccgagctgcaggatgagatcggcacc gccttctctctgtttaagaccgacgaggatatcacaggccgcctgaaggacagggtgcagcctgagatcctggaggccctgctgaagcacatctctttcgataagtttgtgcagatca gcctgaaggccctgagaaggatcgtgccactgatggagcagggcaagcggtacgacgaggcctgcgccgagatctacggcgatcactatggcaagaagaacacagaggagaag atctatctgccccctatccctgccgacgagatcagaaatcctgtggtgctgagggccctgtcccaggcaagaaaagtgatcaacggagtggtgcgccggtacggatctccagcccgg atccacatcgagaccgccagagaagtgggcaagagcttcaaggaccggaaggagatcgagaagagacaggaggagaatcgcaaggatcgggagaaggccgccgccaagttta gggagtacttccctaactttgtgggcgagccaaagtctaaggacatcctgaagctgcgcctgtacgagcagcagcacggcaagtgtctgtatagcggcaaggagatcaatctggtg cggctgaacgagaagggctatgtggagatcgatcacgccctgcctttctccagaacctgggacgattcttttaacaataaggtgctggtgctgggcagcgagaaccagaataaggg caatcagacaccatacgagtatttcaatggcaaggacaactccagggagtggcaggagttcaaggcccgcgtggagacctctagatttcccaggagcaagaagcagcggatcctg ctgcagaagttcgacgaggatggctttaaggagtgcaacctgaatgacaccagatacgtgaaccggttcctgtgccagtttgtggccgatcacatcctgctgaccggcaagggcaag agaagggtgttcgcctctaatggccagatcacaaacctgctgaggggattttggggactgaggaaggtgcgggcagagaatgacagacaccacgcactggatgcagtggtggtgg catgcagcaccgtggcaatgcagcagaagatcacaagattcgtgaggtataaggagatgaacgcctttgacggcaagaccatcgataaggagacaggcaaggtgctgcaccaga agacccacttcccccagccttgggagttctttgcccaggaagtgatgatccgggtgttcggcaagccagacggcaagcctgagtttgaggaggccgataccccagagaagctgagg acactgctggcagagaagctgtctagcaggccagaggcagtgcacgagtacgtgaccccactgttcgtgtccagggcacccaatcggaagatgtctggcgcccacaagggcacac tgagaagcgccaagaggtttgtgaagcacaacgagaagatctccgtgaagagagtgtggctgaccaagatcaggctggccgccctggagaacatggtgaattacaagaacggca gggagatcgagctgtatgaggccctgaaggcaaggctggaggcctacggaggaaatgccaagcaggccttcgacccaaagggcaaccccttttataagaagggaggacagctgg tgaaggccgtgcgggtggagaggacccagaagagcggcgtgctgctgaataagaagaacgcctacacaatcgccgacaatggcgatatggtgagagtggacgtgttctgtaaggt ggataagaagggcaagaatcagtactttatcgtgcctatctatgcctggcaggtggccgagaacatcctgccagacatcgattgcaagggctacagaatcgacgatagctatacatt ctgtttttccctgcacaagtatgacctgatcgccttccagaaggatgagaagtccaaggtggagtttgcctactatatcaattgcgactcctctaacggcaggttctacctggcctggca cgataagggcagcaaggagcagcagtttcgcatctccacccagaatctggtgctgatccagaagtatcaggtgaacgagctgggcaaggagatcaggccatgtcggctgaagaag cgcccacccgtgcgggaggataaaagaaccgccgacggcagcgaattcgagcccaagaagaagaggaaagtc

**iNme2^Smu^-ABE8e-nt**: <u>BPSV40-NLS</u>, Nme2Cas9 – delta PID, <u>TadA8e</u>, <u>SmuCas9 PID</u>, Linkers

Jenifer Dounda Lab, iNme2Cas9 (D16A) variant with SmuCas9 PID swap in n-terminally fused ABE8e format

MKRTADGSEFESPKKKRKVGGSGGGSGGGSGSEVEFSHEYWMRHALTLAKRARDEREVPVGAVLVLNNRVIGEGWNRAIGLHDPTAHAEIM ALRQGGLVMQNYRLIDATLYVTFEPCVMCAGAMIHSRIGRVVFGVRNSKRGAAGSLMNVLNYPGMNHRVEITEGILADECAALLCDFYRMPR QVFNAQKKAQSSINSGGSSGGSSGSETPGTSESATPESSGGSSGGSMAAFKPNPINYILGLAIGIASVGWAMVEIDEEENPIRLIDLGVRVFERAE VPKTGDSLAMARRLARSVRRLTRRRAHRLLRARRLLKREGVLQAADFDENGLIKSLPNTPWQLRAAALDRKLTPLEWSAVLLHLIKHRGYLSQRK NEGETADKELGALLKGVANNAHALQTGDFRTPAELALNKFEKESGHIRNQRGDYSHTFSRKDLQAELILLFEKQKEFGNPHVSGGLKEGIETLLM TQRPALSGDAVQKMLGHCTFEPAEPKAAKNTYTAERFIWLTKLNNLRILEQGSERPLTDTERATLMDEPYRKSKLTYAQARKLLGLEDTAFFKGL RYGKDNAEASTLMEMKAYHAISRALEKEGLKDKKSPLNLSSELQDEIGTAFSLFKTDEDITGRLKDRVQPEILEALLKHISFDKFVQISLKALRRIVPL MEQGKRYDEACAEIYGDHYGKKNTEEKIYLPPIPADEIRNPVVLRALSQARKVINGVVRRYGSPARIHIETAREVGKSFKDRKEIEKRQEENRKDRE KAAAKFREYFPNFVGEPKSKDILKLRLYEQQHGKCLYSGKEINLVRLNEKGYVEIDHALPFSRTWDDSFNNKVLVLGSENQNKGNQTPYEYFNGK DNSREWQEFKARVETSRFPRSKKQRILLQKFDEDGFKECNLNDTRYVNRFLCQFVADHILLTGKGKRRVFASNGQITNLLRGFWGLRKVRAEND RHHALDAVVVACSTVAMQQKITRFVRYKEMNAFDGKTIDKETGKVLHQKTHFPQPWEFFAQEVMIRVFGKPDGKPEFEEADTPEKLRTLLAEK LSSRPEAVHEYVTPLFVSRAPNRKMSGAHKGTLRSAKRFVKHNEKISVKRVWLTKIRLAALENMVNYKNGREIELYEALKARLEAYGGNAKQAFD PKGNPFYKKGGQLVKAVRVERTQKSGVLLNKKNAYTIADNATMVRVDVYTKAGKNYLVPVYVWQVAQGILPNRAVTSGKSEADWDLIDESFE FKFSLSRGDLVEMISNKGRIFGYYNGLDRANGSIGIREHDLEKSKGKDGVHRVGVKTATAFNKYHVDPLGKEIHRCSSEPRPTLKIKSKKEDKRTA DGSEFEPKKKRKV

atgaaacggacagccgacggaagcgagttcgagtcaccaaagaagaagcggaaagtcggcggtagcggcggaggcagcggtggcggcagcggctctgaggtggagttttccca cgagtactggatgagacatgccctgaccctggccaagagggcacgcgatgagagggaggtgcctgtgggagccgtgctggtgctgaacaatagagtgatcggcgagggctggaa cagagccatcggcctgcacgacccaacagcccatgccgaaattatggccctgagacagggcggcctggtcatgcagaactacagactgattgacgccaccctgtacgtgacattcg agccttgcgtgatgtgcgccggcgccatgatccactctaggatcggccgcgtggtgtttggcgtgaggaacagcaaacggggcgccgcaggctccctgatgaacgtgctgaactac cccggcatgaatcaccgcgtcgaaattaccgagggaatcctggcagatgaatgtgccgccctgctgtgcgacttctaccggatgcctagacaggtgttcaatgctcagaagaaggcc cagagctccatcaactccggaggatctagcggaggctcctctggctctgagacacctggcacaagcgagagcgcaacacctgaaagcagcgggggcagcagcggggggtcaatg gccgccttcaagcctaacccaatcaattacatcctgggactggccatcggaatcgcatccgtgggatgggctatggtggagatcgacgaggaggagaatcctatccggctgatcgat ctgggcgtgagagtgtttgagagggccgaggtgccaaagaccggcgattctctggctatggcccggagactggcacggagcgtgaggcgcctgacacggagaagggcacacagg ctgctgagggcacgccggctgctgaagagagagggcgtgctgcaggcagcagacttcgatgagaatggcctgatcaagagcctgccaaacaccccctggcagctgagagcagcc gccctggacaggaagctgacaccactggagtggtctgccgtgctgctgcacctgatcaagcaccgcggctacctgagccagcggaagaacgagggagagacagcagacaaggag ctgggcgccctgctgaagggagtggccaacaatgcccacgccctgcagaccggcgatttcaggacacctgccgagctggccctgaataagtttgagaaggagtccggccacatca gaaaccagaggggcgactatagccacaccttctcccgcaaggatctgcaggccgagctgatcctgctgttcgagaagcagaaggagtttggcaatccacacgtgagcggaggcct gaaggagggaatcgagaccctgctgatgacacagaggcctgccctgtccggcgacgcagtgcagaagatgctgggacactgcaccttcgagcctgcagagccaaaggccgccaa gaacacctacacagccgagcggtttatctggctgacaaagctgaacaatctgagaatcctggagcagggatccgagaggccactgaccgacacagagagggccaccctgatggat gagccttaccggaagtctaagctgacatatgcccaggccagaaagctgctgggcctggaggacaccgccttctttaagggcctgagatacggcaaggataatgccgaggcctccac actgatggagatgaaggcctatcacgccatctctcgcgccctggagaaggagggcctgaaggacaagaagtcccccctgaacctgagctccgagctgcaggatgagatcggcacc gccttctctctgtttaagaccgacgaggatatcacaggccgcctgaaggacagggtgcagcctgagatcctggaggccctgctgaagcacatctctttcgataagtttgtgcagatca gcctgaaggccctgagaaggatcgtgccactgatggagcagggcaagcggtacgacgaggcctgcgccgagatctacggcgatcactatggcaagaagaacacagaggagaag atctatctgccccctatccctgccgacgagatcagaaatcctgtggtgctgagggccctgtcccaggcaagaaaagtgatcaacggagtggtgcgccggtacggatctccagcccgg atccacatcgagaccgccagagaagtgggcaagagcttcaaggaccggaaggagatcgagaagagacaggaggagaatcgcaaggatcgggagaaggccgccgccaagttta gggagtacttccctaactttgtgggcgagccaaagtctaaggacatcctgaagctgcgcctgtacgagcagcagcacggcaagtgtctgtatagcggcaaggagatcaatctggtg cggctgaacgagaagggctatgtggagatcgatcacgccctgcctttctccagaacctgggacgattcttttaacaataaggtgctggtgctgggcagcgagaaccagaataaggg caatcagacaccatacgagtatttcaatggcaaggacaactccagggagtggcaggagttcaaggcccgcgtggagacctctagatttcccaggagcaagaagcagcggatcctg ctgcagaagttcgacgaggatggctttaaggagtgcaacctgaatgacaccagatacgtgaaccggttcctgtgccagtttgtggccgatcacatcctgctgaccggcaagggcaag agaagggtgttcgcctctaatggccagatcacaaacctgctgaggggattttggggactgaggaaggtgcgggcagagaatgacagacaccacgcactggatgcagtggtggtgg catgcagcaccgtggcaatgcagcagaagatcacaagattcgtgaggtataaggagatgaacgcctttgacggcaagaccatcgataaggagacaggcaaggtgctgcaccaga agacccacttcccccagccttgggagttctttgcccaggaagtgatgatccgggtgttcggcaagccagacggcaagcctgagtttgaggaggccgataccccagagaagctgagg acactgctggcagagaagctgtctagcaggccagaggcagtgcacgagtacgtgaccccactgttcgtgtccagggcacccaatcggaagatgtctggcgcccacaagggcacac tgagaagcgccaagaggtttgtgaagcacaacgagaagatctccgtgaagagagtgtggctgaccaagatcaggctggccgccctggagaacatggtgaattacaagaacggca gggagatcgagctgtatgaggccctgaaggcaaggctggaggcctacggaggaaatgccaagcaggccttcgacccaaagggcaaccccttttataagaagggaggacagctgg tgaaggccgtgcgggtggagaggacccagaagagcggcgtgctgctgaataagaagaacgcctacacaatcgccgacaacgccaccatggtgcgggtggacgtgtacaccaagg ccggcaagaactacctggttcctgtgtacgtgtggcaggtggcccagggcatcttacccaaccgcgccgtgaccagcggcaagtccgaggctgactgggacctgatcgatgagagc ttcgagttcaagttctctctgtcccggggagatctcgtggaaatgatctccaacaagggcagaatcttcggctactacaacggcctggacagagccaacggctctattggaattagag agcacgacctagagaagagcaagggcaaagacggcgtgcatagagtgggagtgaaaacagctacagcatttaacaagtaccacgtggatcccctgggcaaagagatccacaga tgcagcagcgaacccagacctacactgaaaatcaagtctaagaaggaggataaaagaaccgccgacggcagcgaattcgagcccaagaagaagaggaaagtc

Nucleotide and amino acid sequences of deaminase linkers used in this study and their orientation with *n*Cas9 domains.

**Nme2^Smu^-ABE8e-i1 (WT)**: <u>BPSV40-NLS</u>, Nme2Cas9 – delta PID, <u>TadA8e</u>, <u>SmuCas9 PID</u>, Deaminase Flanking Linkers

<u>MKRTADGSEFESPKKKRKV</u>ED<u>MAAFKPNPINYILGLAIGIASVGWAMVEIDEEENPIRLIDLGVRVFERAEVPKTGDSLAMARRLARSVRRLTRRRAHRLLRARRLLKREGVLQAADFD</u> <u>ENGLIKSLPNTPWQLRAAALDRKLTPLEWSAVLLHLIKHRGYLSQRKNEGETADKELGALLKGVANNAHALQTGDFRTPAELALNKFEKESGHIRNQRGDYSHTFSRKDLQAELILLFEK</u> <u>QKEFGNPHVSGGLKEGIETLLMTQRPALSGDAVQKMLGHCTFEPAEPKAAKNTYTAERFIWLTKLNNLRILEQ(N_Linker)SEVEFSHEYWMRHALTLAKRARDEREVPVGAVLVLN</u> <u>NRVIGEGWNRAIGLHDPTAHAEIMALRQGGLVMQNYRLIDATLYVTFEPCVMCAGAMIHSRIGRVVFGVRNSKRGAAGSLMNVLNYPGMNHRVEITEGILADECAALLCDFYRMP</u> <u>RQVFNAQKKAQSSIN</u>(C_Linker)<u>GSERPLTDTERATLMDEPYRKSKLTYAQARKLLGLEDTAFFKGLRYGKDNAEASTLMEMKAYHAISRALEKEGLKDKKSPLNLSSELQDEIGTAFSLF</u> <u>KTDEDITGRLKDRVQPEILEALLKHISFDKFVQISLKALRRIVPLMEQGKRYDEACAEIYGDHYGKKNTEEKIYLPPIPADEIRNPVVLRALSQARKVINGVVRRYGSPARIHIETAREVGKSF</u> <u>KDRKEIEKRQEENRKDREKAAAKFREYFPNFVGEPKSKDILKLRLYEQQHGKCLYSGKEINLVRLNEKGYVEIDHALPFSRTWDDSFNNKVLVLGSENQNKGNQTPYEYFNGKDNSREW</u> <u>QEFKARVETSRFPRSKKQRILLQKFDEDGFKECNLNDTRYVNRFLCQFVADHILLTGKGKRRVFASNGQITNLLRGFWGLRKVRAENDRHHALDAVVVACSTVAMQQKITRFVRYKE</u> <u>MNAFDGKTIDKETGKVLHQKTHFPQPWEFFAQEVMIRVFGKPDGKPEFEEADTPEKLRTLLAEKLSSRPEAVHEYVTPLFVSRAPNRKMSGAHKDTLRSAKRFVKHNEKISVKRVWLT</u> <u>EIKLADLENMVNYKNGREIELYEALKARLEAYGGNAKQAFDPKDNPFYKKGGQLVKAVRVEKTQESGVLLNKKNAYTIADNATMVRVDVYTKAGKNYLVPVYVWQVAQGILPNRAV</u> <u>TSGKSEADWDLIDESFEFKFSLSRGDLVEMISNKGRIFGYYNGLDRANGSIGIREHDLEKSKGKDGVHRVGVKTATAFNKYHVDPLGKEIHRCSSEPRPTLKIKSKK</u>ED<u>KRTADGSEFEPK</u> <u>KKRKV</u>

**Nme2^Smu^-ABE8e-i8 (WT)**: <u>BPSV40-NLS</u>, Nme2Cas9 – delta PID, <u>TadA8e</u>, <u>SmuCas9 PID</u>, Deaminase Flanking Linkers

<u>MKRTADGSEFESPKKKRKV</u>ED<u>MAAFKPNPINYILGLAIGIASVGWAMVEIDEEENPIRLIDLGVRVFERAEVPKTGDSLAMARRLARSVRRLTRRRAHRLLRARRLLKREGVLQAADFD</u> <u>ENGLIKSLPNTPWQLRAAALDRKLTPLEWSAVLLHLIKHRGYLSQRKNEGETADKELGALLKGVANNAHALQTGDFRTPAELALNKFEKESGHIRNQRGDYSHTFSRKDLQAELILLFEK</u> <u>QKEFGNPHVSGGLKEGIETLLMTQRPALSGDAVQKMLGHCTFEPAEPKAAKNTYTAERFIWLTKLNNLRILEQGSERPLTDTERATLMDEPYRKSKLTYAQARKLLGLEDTAFFKGLRY</u> <u>GKDNAEASTLMEMKAYHAISRALEKEGLKDKKSPLNLSSELQDEIGTAFSLFKTDEDITGRLKDRVQPEILEALLKHISFDKFVQISLKALRRIVPLMEQGKRYDEACAEIYGDHYGKKNTE</u> <u>EKIYLPPIPADEIRNPVVLRALSQARKVINGVVRRYGSPARIHIETAREVGKSFKDRKEIEKRQEENRKDREKAAAKFREYFPNFVGEPKSKDILKLRLYEQQHGKCLYSGKEINLVRLNEKG</u> <u>YVEIDHALPFSRTWDDSFNNKVLVLGSENQNKGNQTPYEYFNGKDNSREWQEFKARVETSRFPRSKKQRILLQKFDEDGFKECNLNDTRYVNRFLCQFVADHILLTGKGKRRVFASNG</u> <u>QITNLLRGFWGLRKVRAENDRHHALDAVVVACSTVAMQQKITRFVRYKEMNAFDGKTIDKETGKVLHQKTHFPQPWEFFAQEVMIRVFGKPDGKP</u>(N_Linker)<u>SEVEFSHEYWM</u> <u>RHALTLAKRARDEREVPVGAVLVLNNRVIGEGWNRAIGLHDPTAHAEIMALRQGGLVMQNYRLIDATLYVTFEPCVMCAGAMIHSRIGRVVFGVRNSKRGAAGSLMNVLNYPGM</u> <u>NHRVEITEGILADECAALLCDFYRMPRQVFNAQKKAQSSIN(C_Linker)EFEEADTPEKLRTLLAEKLSSRPEAVHEYVTPLFVSRAPNRKMSGAHKDTLRSAKRFVKHNEKISVKRVWL</u> <u>TEIKLADLENMVNYKNGREIELYEALKARLEAYGGNAKQAFDPKDNPFYKKGGQLVKAVRVEKTQESGVLLNKKNAYTIADNATMVRVDVYTKAGKNYLVPVYVWQVAQGILPNRA</u> <u>VTSGKSEADWDLIDESFEFKFSLSRGDLVEMISNKGRIFGYYNGLDRANGSIGIREHDLEKSKGKDGVHRVGVKTATAFNKYHVDPLGKEIHRCSSEPRPTLKIKSKK</u>ED<u>KRTADGSEFEP</u> <u>KKKRKV</u>

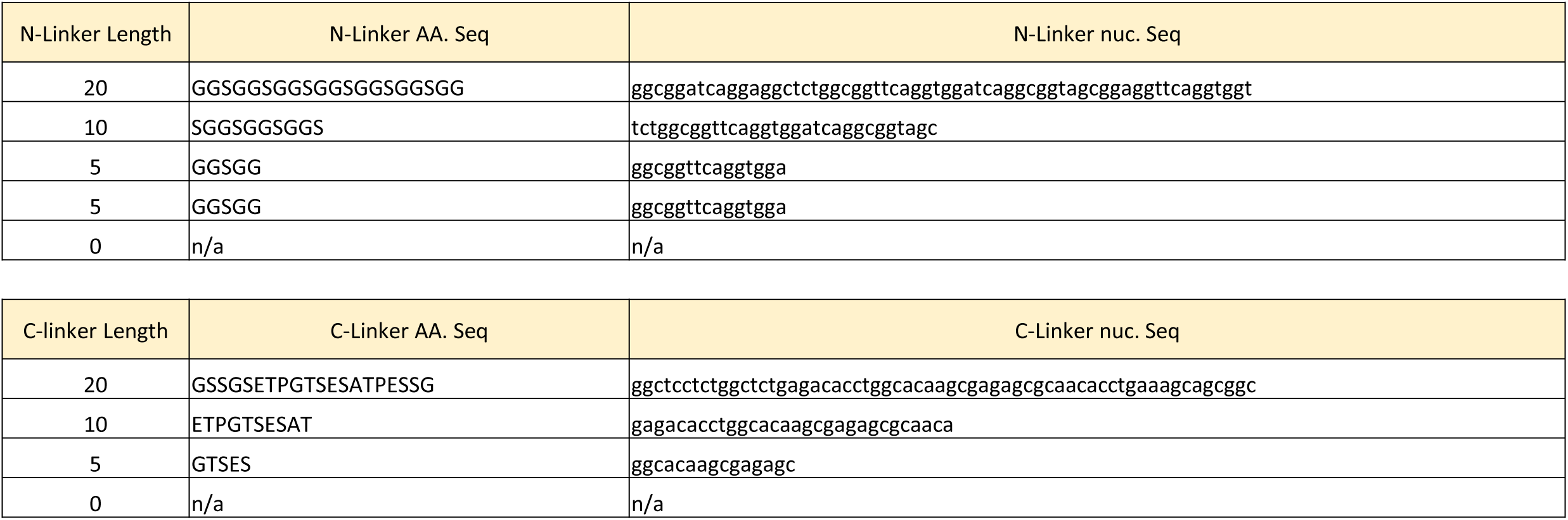

Narrow window TadA deaminase variants used in this study

**TAD9e: Tu et al. *Mol* Ther. 2022. DOI: 10.1016/j.ymthe.2022.07.010**

TCTGAGGTGGAGTTTTCCCACGAGTACTGGATGAGACATGCCCTGACCCTGGCCAAGAGGGCACGCGATGAGAGGGAGGTGCCTGTGGGAGCCGTGCTGGTGCTGAACAAT AGAGTGATCGGCGAGGGCTGGAACAGAGCCATCGGCCTGCACGACCCAACAGCCCATGCCGAAATTATGGCCCTGAGACAGGGCGGCCTGGTCATGCAGAACTACAGACTG ATTGACGCCACCCTGTACGTGACATTCGAGCCTTGCGTGATGTGCGCCGGCGCCATGATCCACTCTAGGATCGGCCGCGTGGTGTTTGGCGTGAGGAACAGCAAAACCGGCGC CGCAGGCTCCCTGATGAACGTGCTGAACTACCCCGGCATGAATAAGCACCGCGTCGAAATTACCGAGGGAATCCTGGCAGATGAATGTGCCGCCCTGCTGTGCGACTTCTACC GGATGCCTAGAAGACAGGTGTTCAATGCTCAGAAGAAGGCCCAGAGCTCCATCAAC

**TAD9: Chen et al. *Nat Chem Biol*. 2023. DOI: 10.1038/s41589-022-01163-8.**

TCTGAGGTGGAGTTTTCCCACGAGTACTGGATGAGACATGCCCTGACCCTGGCCAAGAGGGCACGGGATGAGAGGGAGGTGCCTGTGGGAGCCGTGCTGG TGCTGAACAATAGAGTGATCGGCGAGGGCTGGAACAGAGCCATCGGCCTGCACGACCCAACAGCCCATGCCGAAATTATGGCCCTGAGACAGGGCGGCCT GGTCATGCAGAACTACAGACTGATTGACGCCACCCTGTACGTGACATTCGAGCCTTGCGTGATGTGCGCCGGCGCCATGATCCACTCTAGGATCGGCCGCGT GGTGTTTGGCGTGAGGCAGTCAAAAAGAGGCGCCGCAGGCTCCCTGATGAACGTGCTGAACTACCCCGGCATGAATCACCGCGTCGAAATTACCGAGGGA ATCCTGGCAGATGAATGTGCCGCCCTGACCTGCGATTTCTATCGGATGCCTAGACAGGTGTTCAATGCTCAGAAGAAGGCCCAGAGCTCCATCAAC

---

**Supplementary Figure 1.**
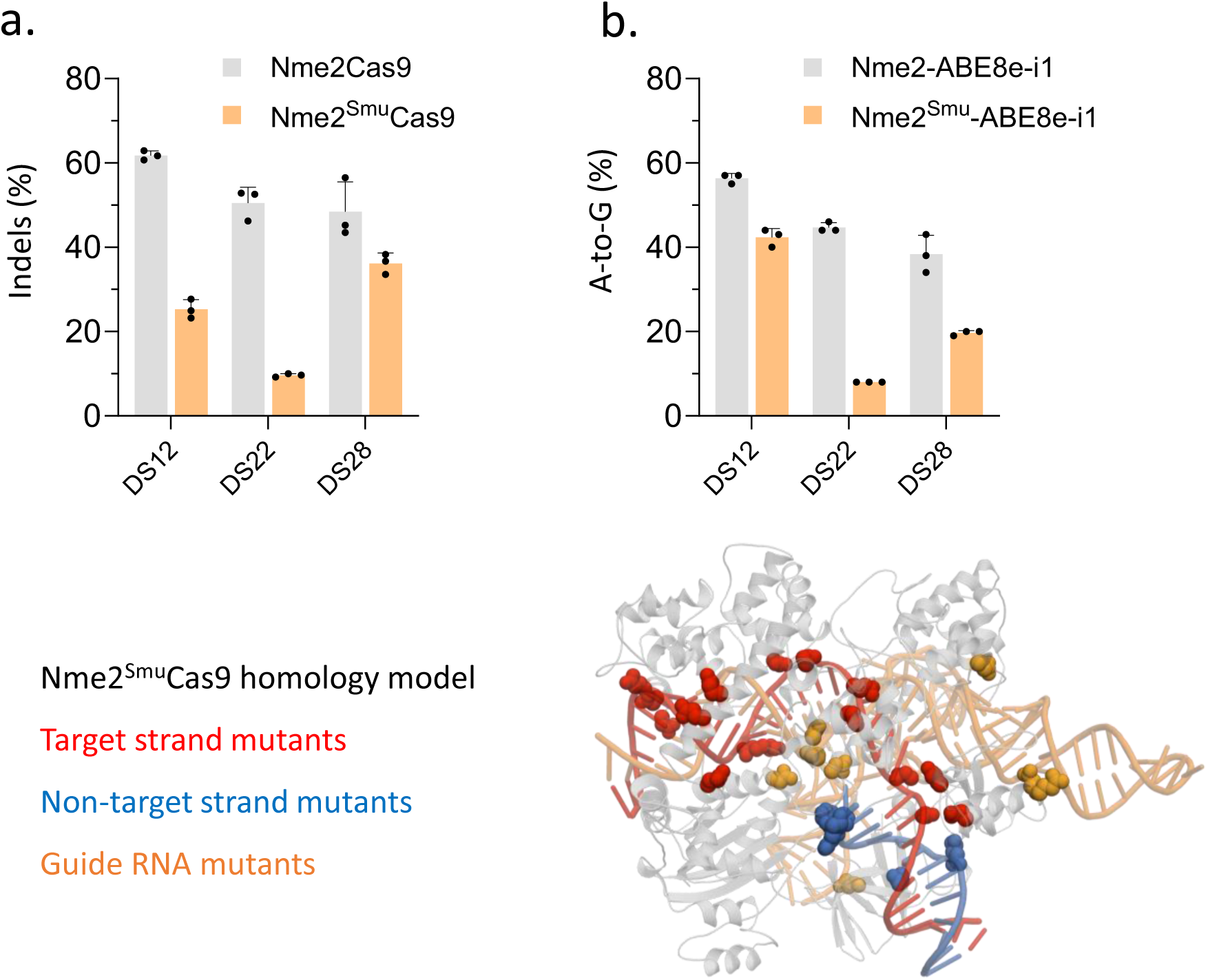
PID-chimeric Nme2^Smu^Cas9 nucleases and ABEs perform poorly at N_4_CC targets compared to WT PID Nme2Cas9 effectors. (**a**) Nuclease editing, following transfection of Nme2- or Nme2^Smu^Cas9 and associated sgRNA plasmids, at endogenous N_4_CC PAM genomic loci in HEK293T cells. Editing efficiencies were measured by amplicon deep sequencing (n = 3 biological replicates; data represent mean ± SD). (**b**) Editing with Nme2- or Nme2^Smu^-ABE8e-i1 plasmids at N_4_CC PAM target loci in HEK293T cells. The editing efficiency at the maximally edited adenine for each target was plotted. Editing efficiencies were measured by amplicon deep sequencing (n = 3 biological replicates; data represent mean ± SD).

**Supplementary Figure 2.**
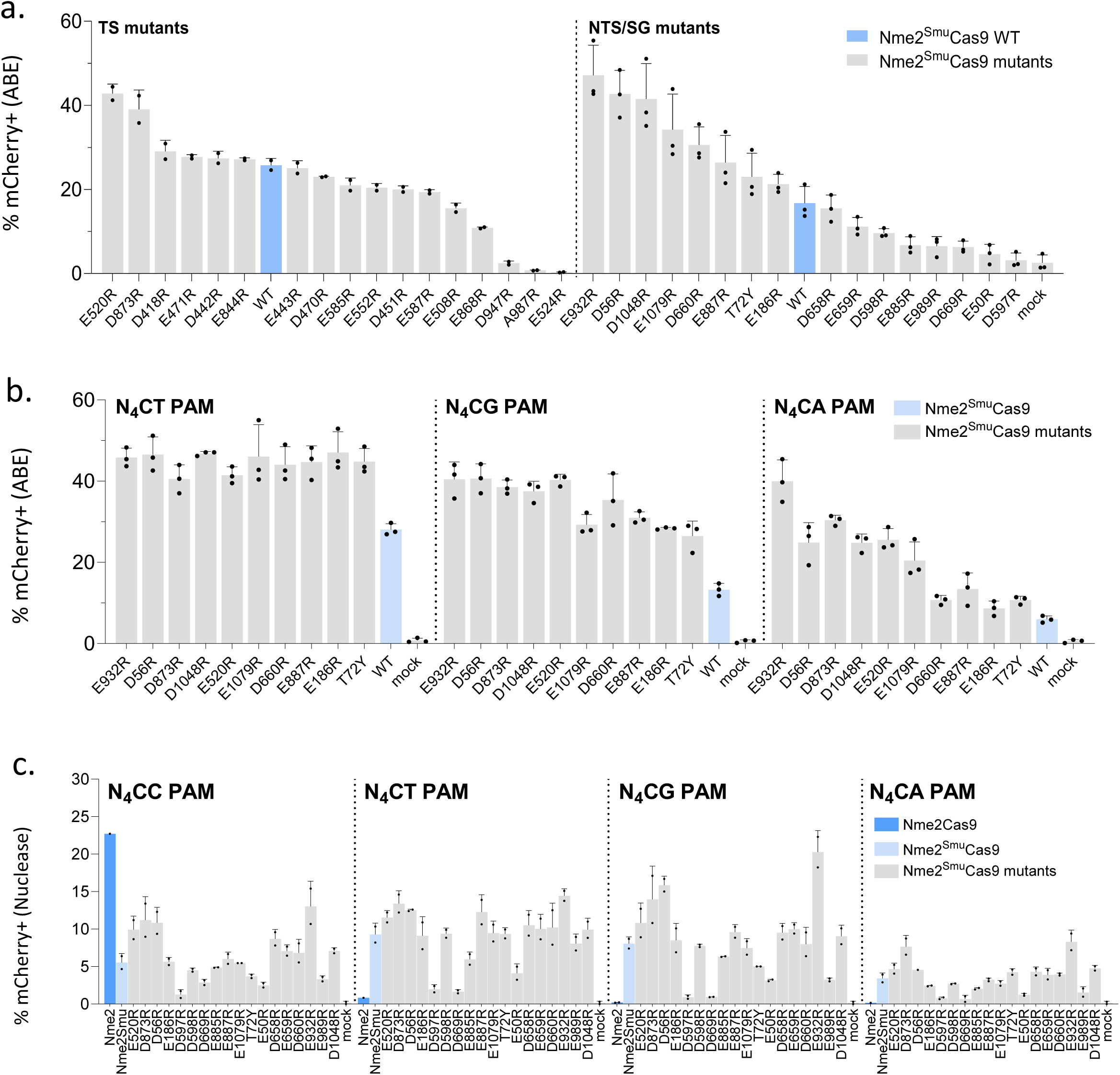
Arginine mutagenesis improves Nme2^Smu^Cas9 effector activity. (**a**) Activities of Nme2^Smu^-ABE8e-i1 (denoted as WT, blue bar) and nucleic acid-interacting arginine mutants [target DNA strand (TS), single guide RNA (SG), and non-target DNA strand (NTS)] denoted by amino acid substitution (grey bars) in the mCherry ABE reporter cell line (activated upon A-to-G editing). After plasmid transfection with an N_4_CC PAM-targeting sgRNA plasmid and an effector-expressing plasmid, editing activities were measured by flow cytometry (n = 2 or 3 biological replicates; data represent mean ± SD). (**b**) Activities of Nme2^Smu^-ABE8e-i1 and the top 10 performing arginine mutants in the mCherry ABE reporter cell line (activated upon A-to-G editing) at N_4_CD (D = not C) PAM targets. After plasmid transfection with the associated sgRNA plasmid and the effector-expressing plasmid, activities were measured by flow cytometry (n = 3 biological replicates; data represent mean ± SD). (**c**) Activities of Nme2Cas9 nuclease variants within the HEK293T TLR-MCV1 reporter at N_4_CC, N_4_CT, N_4_CG and N_4_CA PAM targets, comparing Nme2Cas9 (dark blue), Nme2^Smu^Cas9 (light blue), and Nme2^Smu^Cas9 arginine mutants (grey). After parallel plasmid transfection with the sgRNA and nuclease effector plasmids, activities were measured by flow cytometry (n = 2 biological replicates; data represent mean ± SD).

**Supplementary Figure 3.**
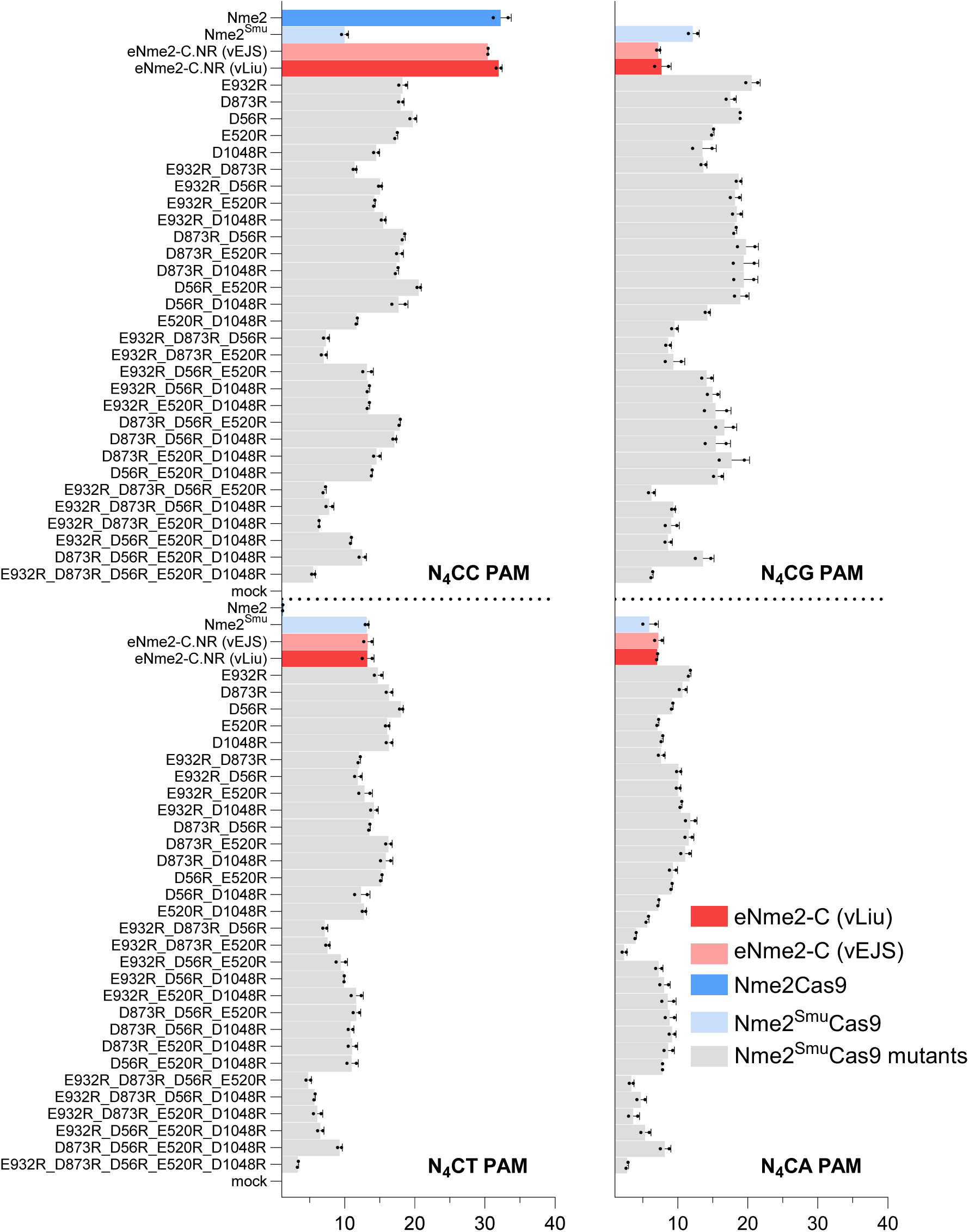
Arginine mutagenesis improves Nme2^Smu^Cas9 nuclease activity. Activities of Nme2Cas9 nuclease variants with the HEK293T TLR-MCV1 reporter at N_4_CC, N_4_CT, N_4_CG and N_4_CA PAM targets, comparing Nme2Cas9, eNme2-C.NR (vLiu), eNme2-C.NR (vEJS), Nme2^Smu^Cas9, and Nme2^Smu^Cas9 single and multiple arginine mutants. For eNme2-C.NR, vLiu is the original plasmid obtained from Addgene (#185672), whereas vEJS is the same effector re-cloned into the expression plasmid backbone used for all other effectors in this experiment. Activities were measured after parallel plasmid transfection with sgRNA and nuclease editor plasmids, followed by flow cytometry (n = 2 biological replicates; data represent mean ± SD).

**Supplementary Figure 4.**
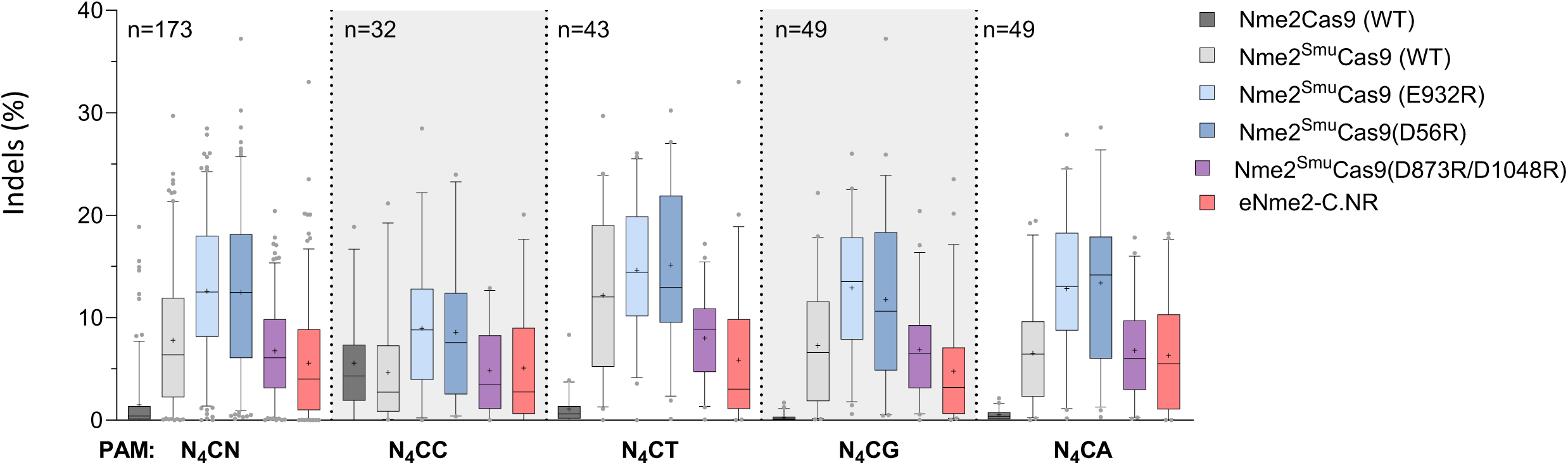
Activities of Nme2^Smu^Cas9 nuclease variants. Nuclease-induced indels in experimental panel 1 of the guide-target activity library following plasmid transfection of Nme2Cas9 (WT), Nme2^Smu^Cas9 (WT and E932R, D56R, and E520R/D873R variants) or eNme2-C.NR into HEK293T cells. The editing efficiencies for 173 target sites were plotted. Editing activities were measured by amplicon sequencing (n = 3 biological replicates; boxplots represent median and interquartile range; whiskers indicate 5th and 95th percentiles and the cross represents the mean).

**Supplementary Figure 5.**
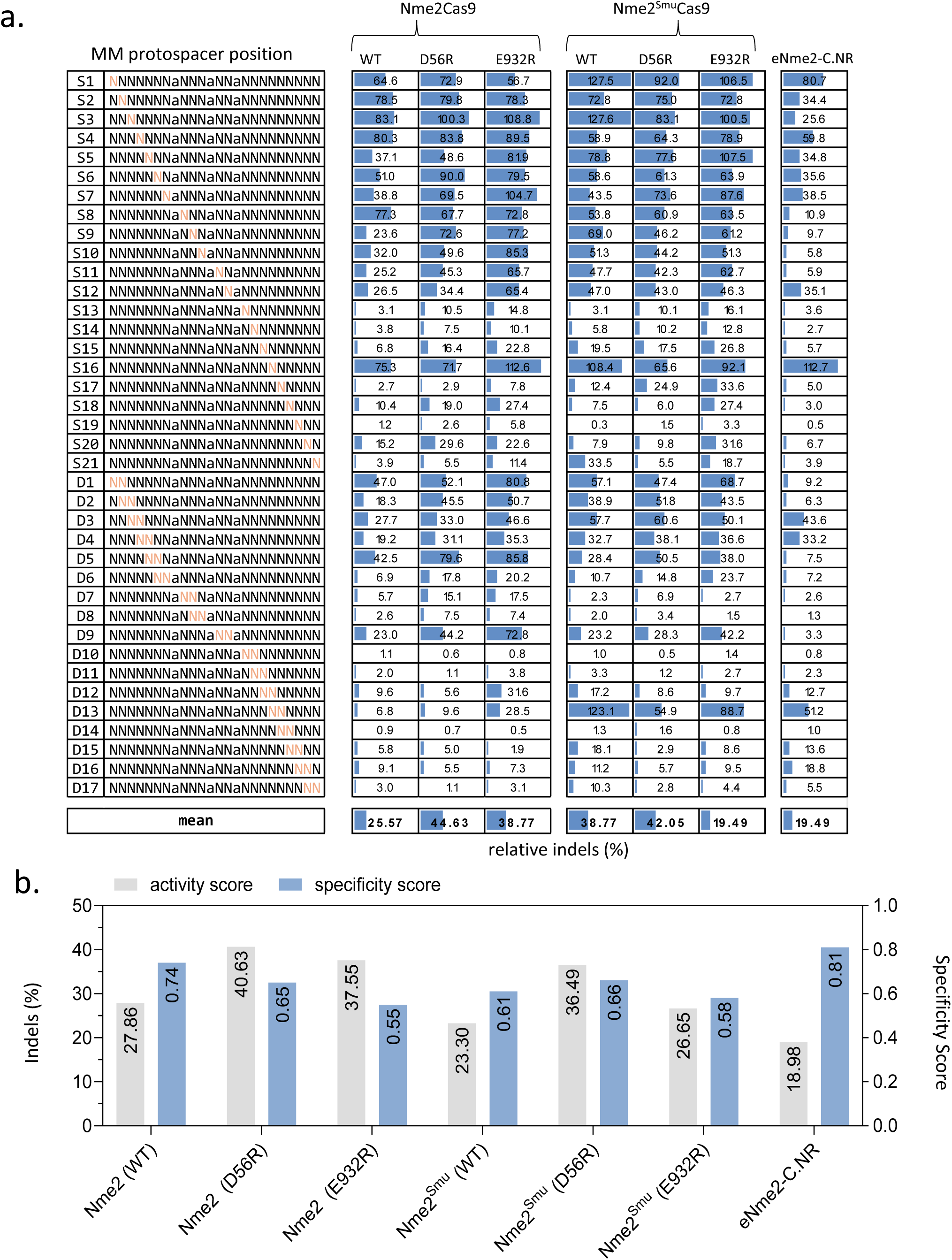
Specificities of Nme2- and Nme2^Smu^Cas9 nucleases at N_4_CC PAM targets. (a) Indel editing frequencies of Nme2Cas9, Nme2^Smu^Cas9 variants, and eNme2-C.NR across single-(S) or di-nucleotide (D) mismatched target sites within the guide-target mismatch library. Activities for each mismatched target were normalized to the mean efficiency of their respective perfectly-matched target site. Orange nucleotides represent protospacer positions of the transversion mutation(s) present within the mismatched target site. (**b**) indel activity vs. specificity score for nuclease variants from (**a**) across the mismatched guide-target library. Nuclease activity was compiled from editing data for three perfectly matched N_4_CC target sites (0 MM) (**Supplementary data file**). The specificity score was calculated as one minus the tiled mismatched editing mean from (**a**), normalized to a scale of one to 100. Editing activities were measured by amplicon sequencing (n = 3 biological replicates).

**Supplementary Figure 6.**
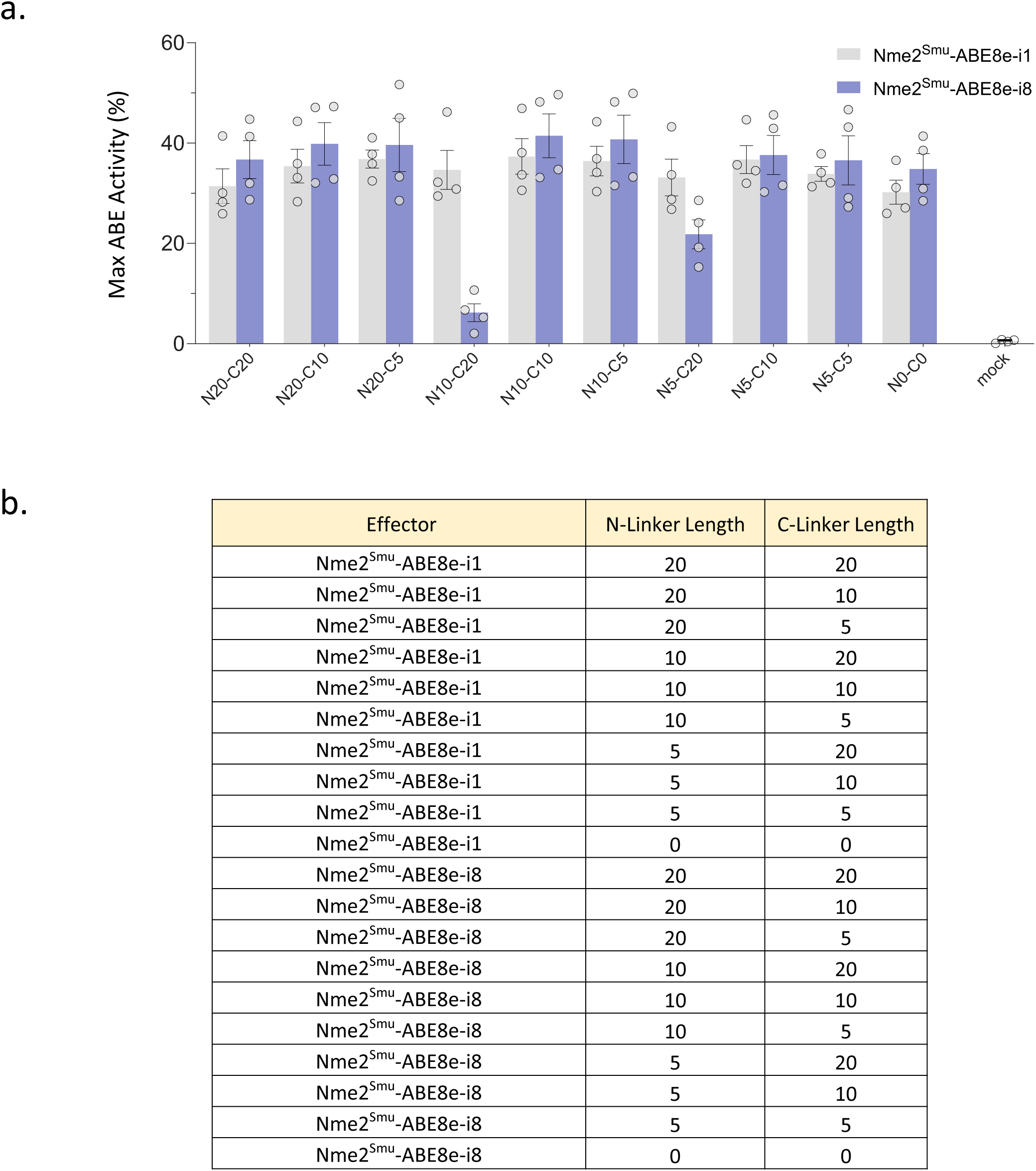
Domain-inlaid Nme2^Smu^-ABE8e linker length optimization. (**a**) A-to-G editing at four endogenous HEK293T genomic loci with Nme2^Smu^-ABE8e-i1 (grey bars) or Nme2^Smu^-ABE8e-i8 (blue bars) carrying N- or C-terminal (Nx-Cx) linker variants following plasmid transfection. The editing efficiencies at the maximally edited adenine for each target was plotted and aggregated. Data for individual target sites are in the **Supplementary data file**. Editing efficiencies were measured by amplicon deep sequencing (n = 3 biological replicates; data represent mean ± SD).

**Supplementary Figure 7.**
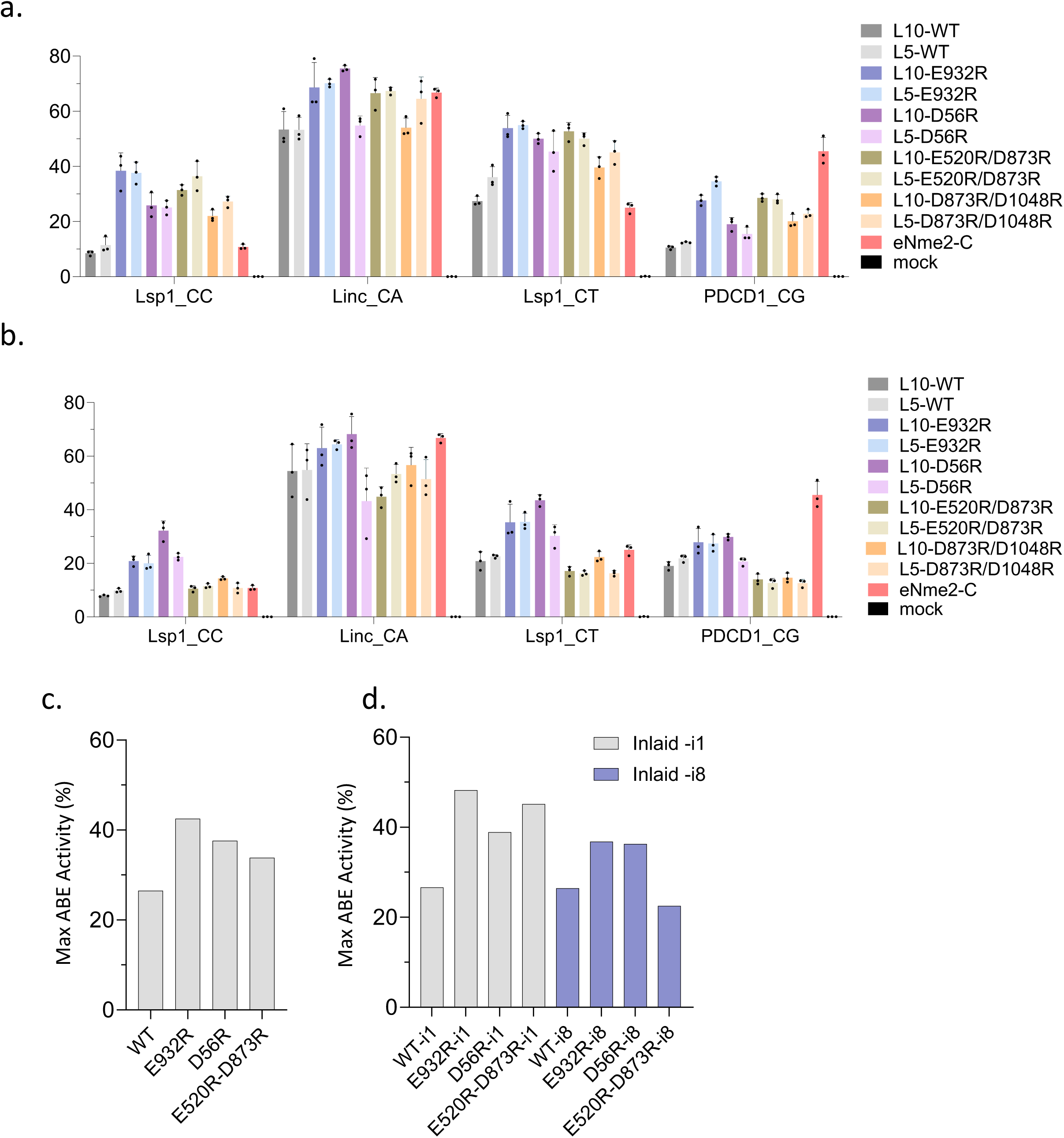
Domain-inlaid Nme2^Smu^-ABE8e linker length and activity optimization. (**a-b**) A-to-G editing at four endogenous HEK293T genomic loci with Nme2^Smu^-ABE8e-i1 (**a**) or Nme2^Smu^-ABE8e-i8 (**b**) arginine mutants (WT and E932R, D56R, E520R/D873R, D873R/D1048R) and linker (L10, L5) variants by plasmid transfection. The editing efficiency at the maximally edited adenine for each target was plotted. Editing efficiencies were measured by amplicon deep sequencing (n = 3 biological replicates; data represent mean ± SD). Data for individual target sites are in the **Supplementary data file**. (**c-d**) Summary data from endogenous maximum activity aggregated from (**a**) and (**b**). (**c**) Nme2^Smu^-ABE8e and arginine mutant activity independent of domain insertion site and linker length, or (**d**) separated by position of domain insertion (-i1 vs. –i8).

**Supplementary Figure 8.**
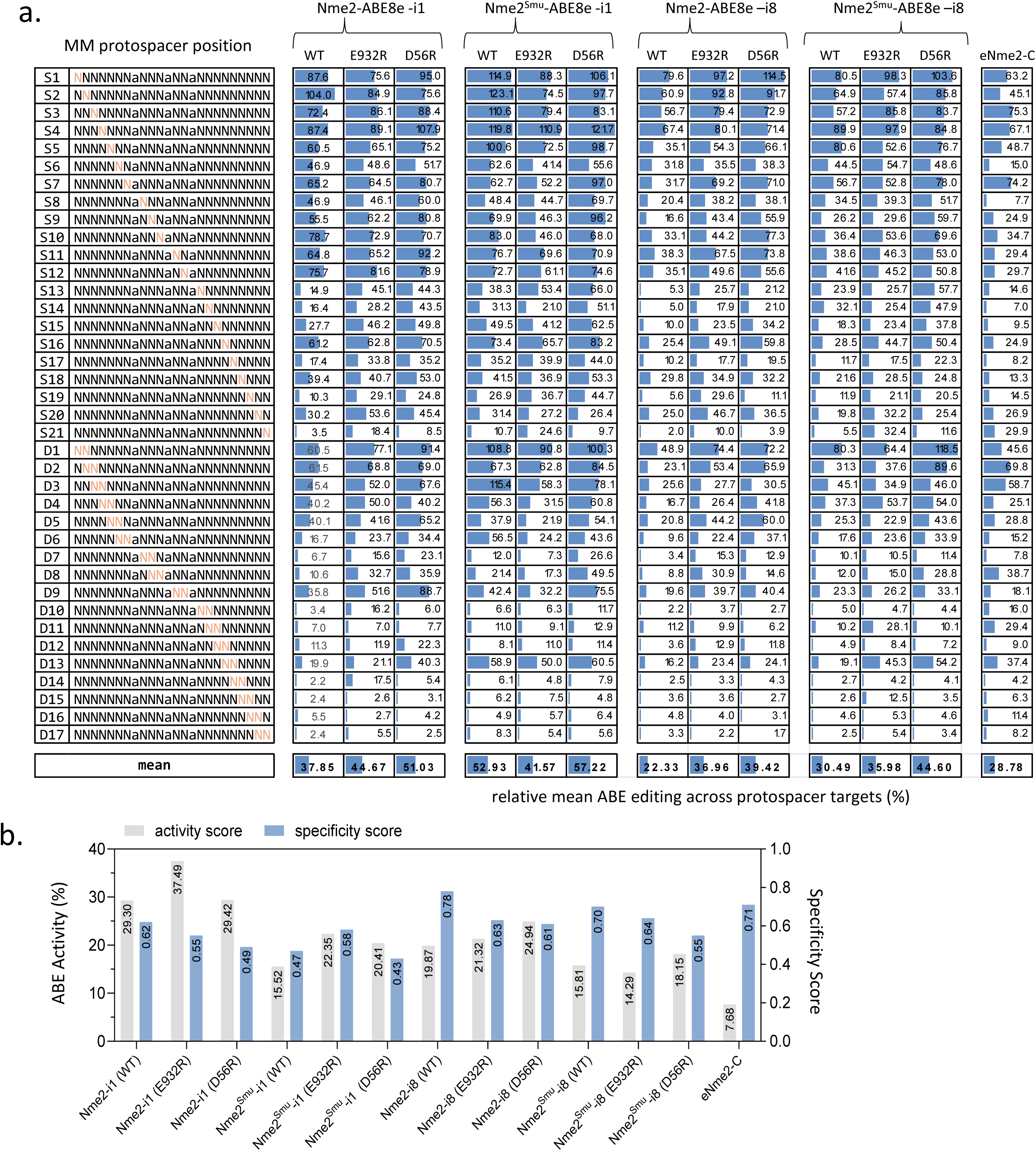
Specificities of domain-inlaid Nme2- and Nme2^Smu^-ABE8e variants at N_4_CC PAM targets. (**a**) Mean A-to-G editing frequencies of domain-inlaid Nme2^Smu^-ABE8e variants or eNme2-C across single-(S) or di-nucleotide (D) mismatched target sites within the guide-target mismatch library. Activities for each mismatched target were normalized to the mean efficiency of their respective perfectly matched target site. Orange nucleotides represent protospacer position of the transversion mutation(s) present within the mismatched target site. (**b**) ABE activity vs. specificity score for base editing variants in (**a**) across the mismatched guide-target library. ABE activity was compiled from editing data for three perfectly matched N_4_CC target sites (0 MM) (**Supplementary data file**). The specificity scores were calculated as one minus the tiled mismatched editing mean in (**a**) normalized to a scale of one to 100. Editing activities were measured by amplicon sequencing (n = 3 biological replicates).

**Supplementary Figure 9.**
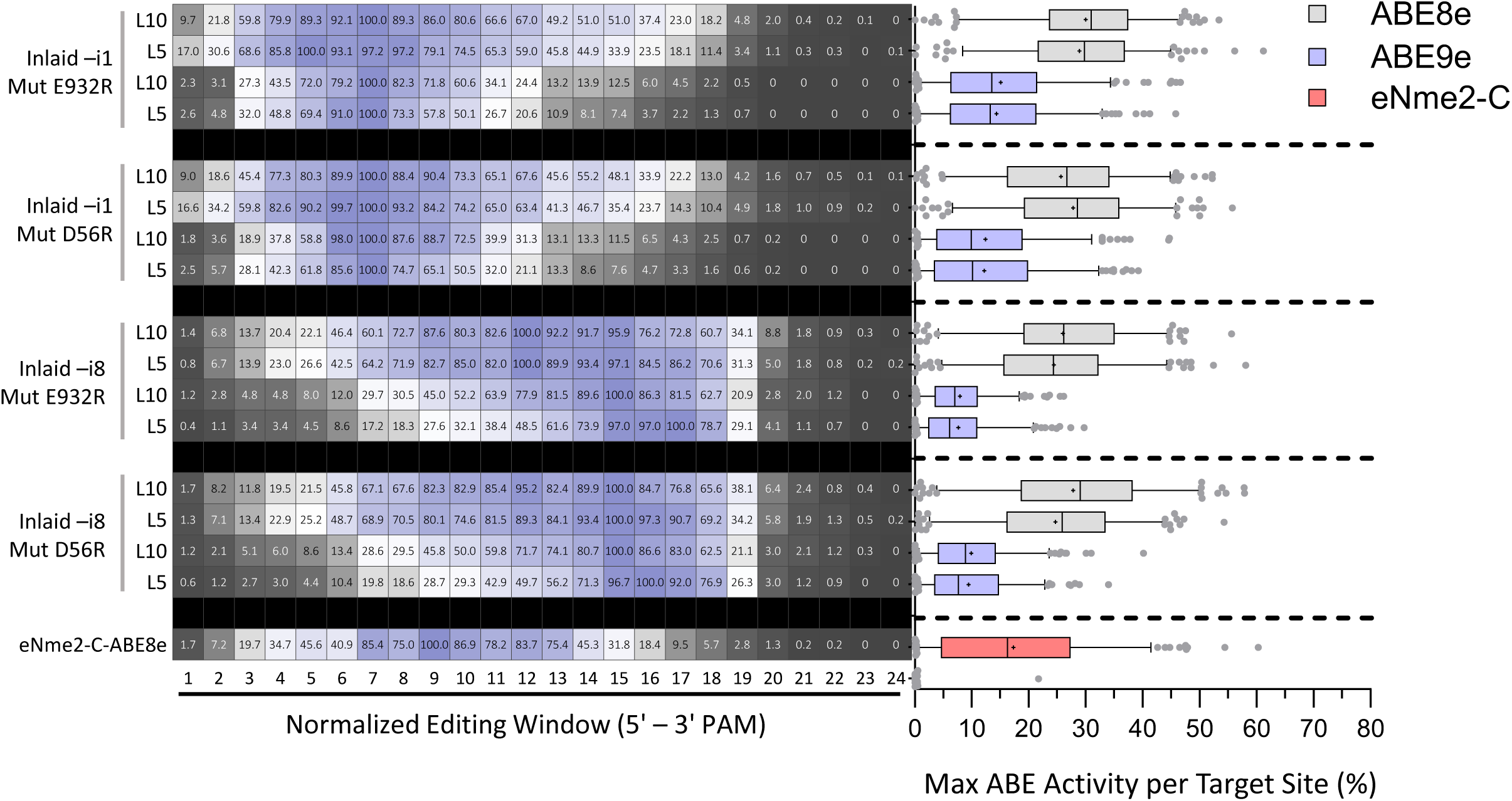
Editing windows of domain-inlaid Nme2^Smu^-ABEs with narrow-window adenine deaminases. Assessment of editing windows and activities from experimental panel 3 of the guide-target activity library (193 sites) for broad-or narrow-window deaminases (ABE8e or ABE9e, respectively). Test subjects include Nme2^Smu^-ABE –i1 or –i8 with E932R or D56R arginine mutants, in combination with deaminase linker lengths (L10 and L5). The eNme2-C variant was also included (bottom). Editing was assessed following plasmid transfection of the ABE variants into HEK293T cells with the integrated guide-target library. Left: average editing windows across the target sites, normalized on a scale of 0 – 100 (%) against adenine positions with the highest observed edited efficiencies within the window. Right: activity at the maximally edited adenine for each target was plotted. Editing activities were measured by amplicon sequencing (n = 3 biological replicates; boxplots represent median and interquartile ranges; whiskers indicate 5th and 95th percentiles; the cross represents the mean).

**Supplementary Figure 10.**
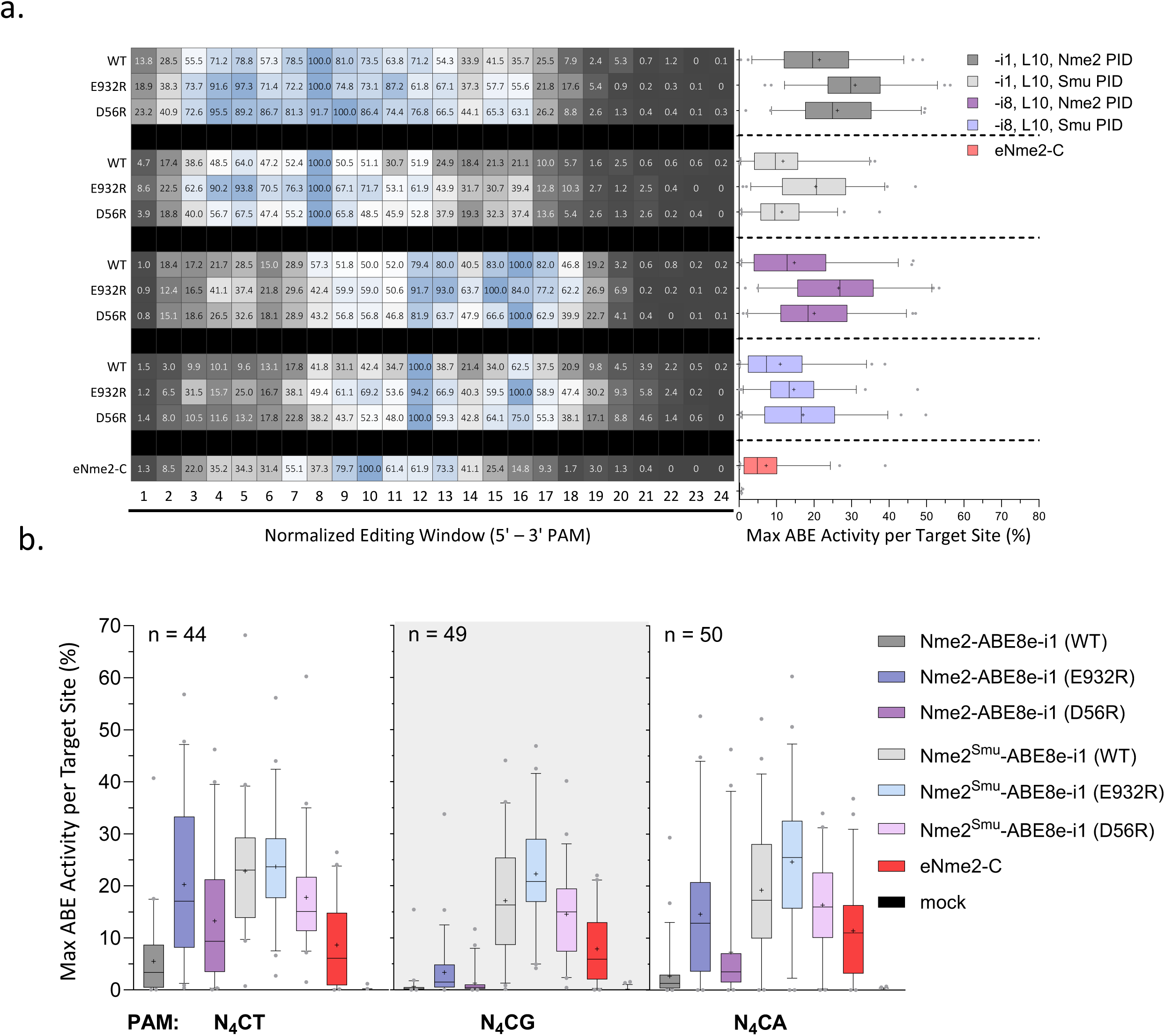
Activities and editing windows of domain-inlaid Nme2- and Nme2^Smu^-ABE8e variants at N_4_CC or N_4_CN PAM targets. Assessment of editing windows and activities from experimental panel 2 of the guide-target activity library (192 sites) for Nme2- and Nme2^Smu-^ABE8e –i1 or –i8, with and without the E932R or D56R arginine mutations, in combination with the L10 deaminase linker. The eNme2-C variant was also included (bottom). (**a**) Editing data for N_4_CC PAM targets only (49 sites). Following plasmid transfection of the ABE variants into HEK293T cells with the integrated guide-target library, editing activities were measured by amplicon sequencing. Left: average editing windows across target sites, normalized on a scale of 0 – 100 (%) against adenine positions with the highest observed edited efficiencies within the window. Right: editing efficiency at the maximally edited adenine for each target. (**b**) Nme2- or Nme2^Smu^-ABE8e-i1 editing data at N_4_CD PAM target sites. The maximally edited adenine for each target was plotted. *n*, the number of target sites with each PAM (n = 3 biological replicates; boxplots represent median and interquartile ranges; whiskers indicate 5th and 95th percentiles; the cross represents the mean).

**Supplementary Figure 11.**
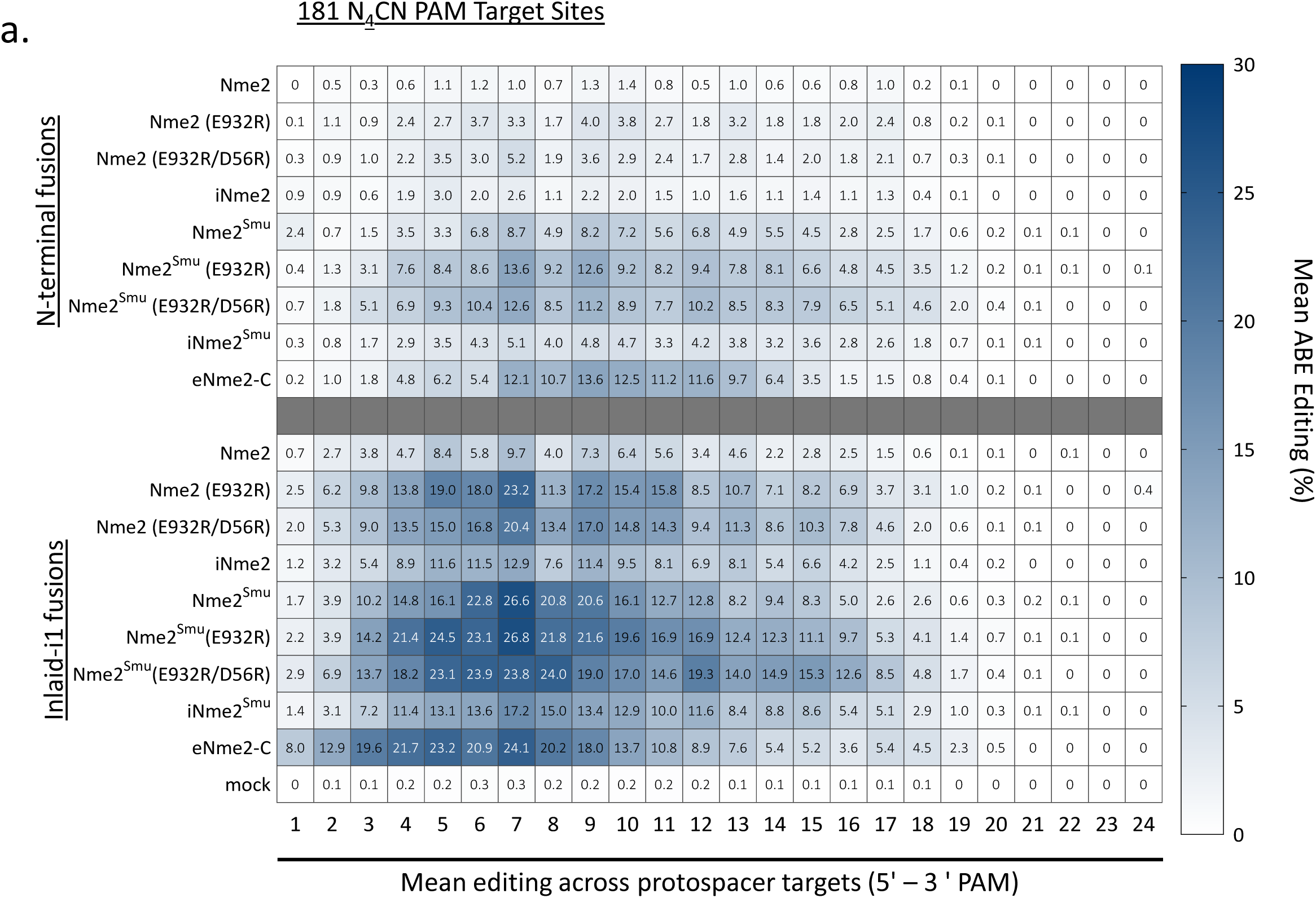
Activities and editing windows of engineered Nme2Cas9 ABE8e variants at N_4_CN PAM targets. Assessment of editing windows and activities from experimental panel 4 of the guide-target activity library (181 N_4_CN PAM sites) for Nme2-, Nme2^Smu^-, iNme2-, iNme2^Smu^- and eNme2-C variants in either the N-terminal or inlaid-i1 (linker 10) formats. Mean A-to-G editing activities and editing windows across protospacer positions in the activity guide-target library are shown for the engineered Nme2Cas9 ABE8e variants. Editing activities were measured by amplicon sequencing (n = 3 biological replicates).

**Supplementary Figure 12.**
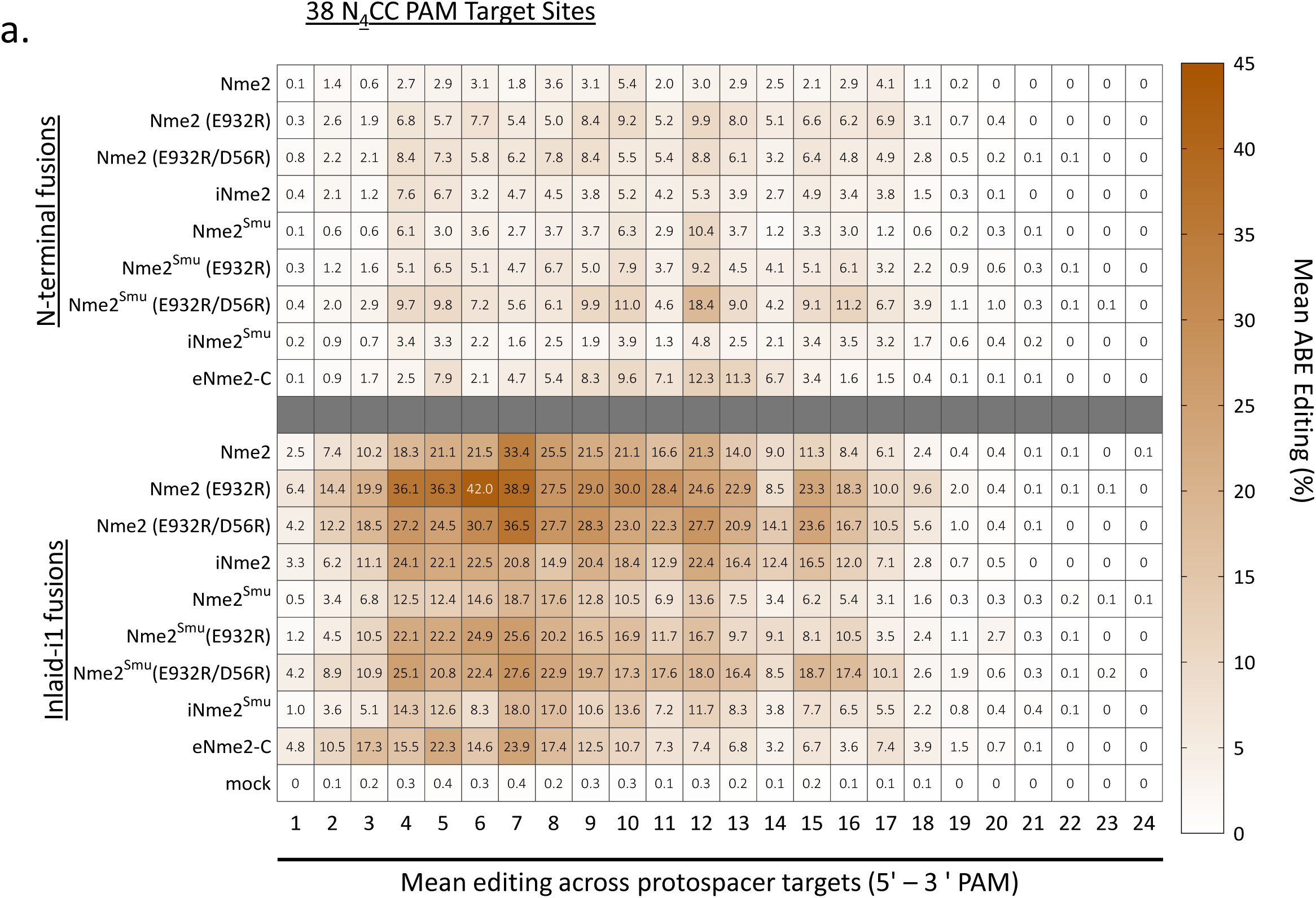
Activities and editing windows of engineered Nme2Cas9 ABE8e variants at N_4_CC PAM targets. Assessment of editing windows and activities from experimental panel 4 of the guide-target activity library (38 N_4_CC PAM sites) for Nme2-, Nme2^Smu^-, iNme2-, iNme2^Smu^- and eNme2-C variants in either the N-terminal or inlaid-i1 (linker 10) formats. Mean A-to-G editing activities and editing windows across protospacer positions in the activity guide-target library are shown for the engineered Nme2Cas9 ABE8e variants. Editing activities were measured by amplicon sequencing (n = 3 biological replicates).

**Supplementary Figure 13.**
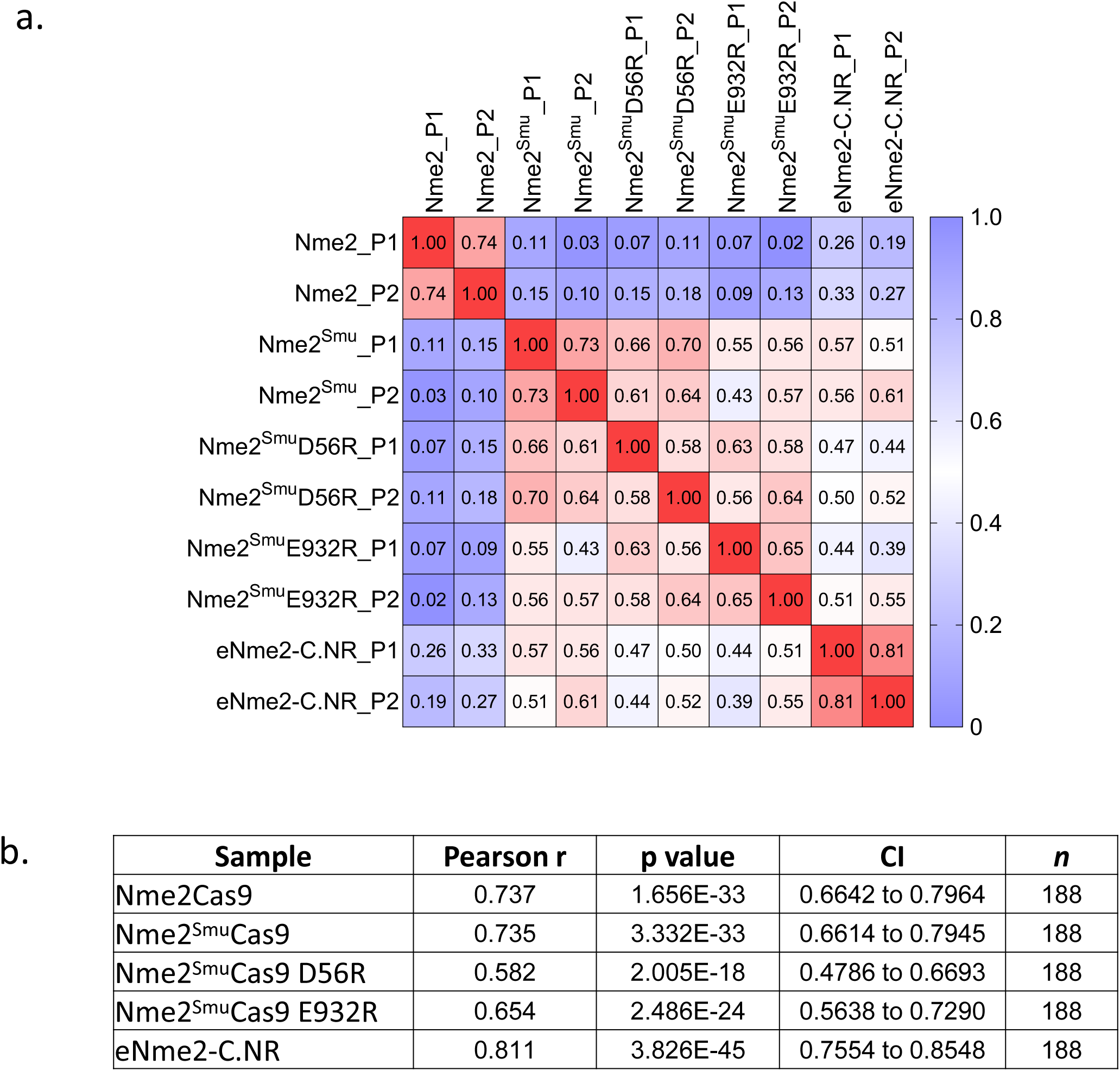
Guide-target library quality control for nuclease editing panels. (**a**) Pearson correlation between experimental panels of nuclease guide-target library experiments, comparing the mean editing efficiencies of library member target sites. Each experimental panel consisted of 3 biological replicates (Rep1, 2 or 3), compiled from **Supplementary** Figure 4 [panel 1, (P1)] and Figure 2a [panel 2, (P2)]. (**b**) Summary statistics from (**a**); CI indicates confidence interval, and *n* indicates number of comparisons used.

**Supplementary Figure 14.**
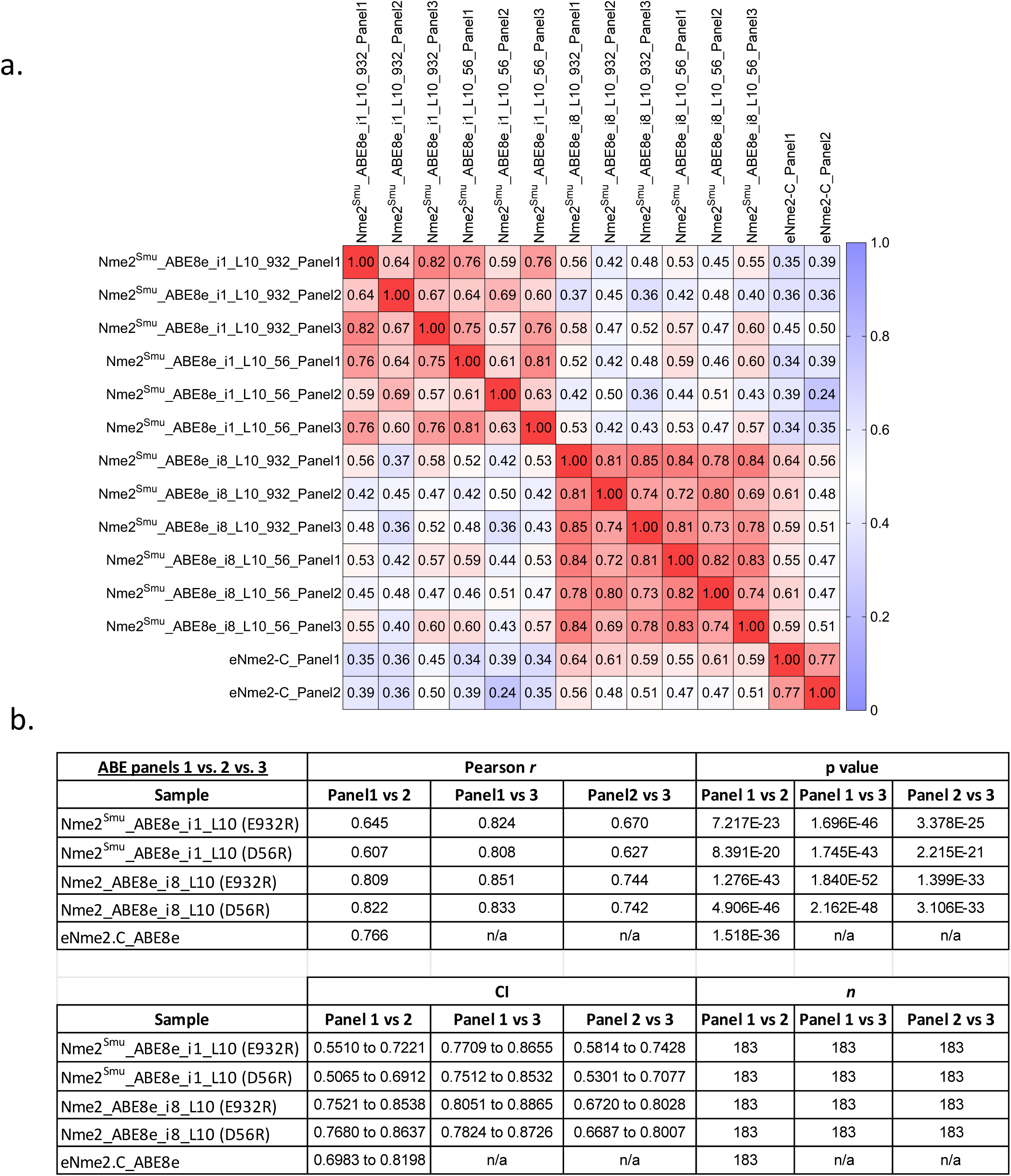
Guide-target quality control for ABE editing panels. (**a**) Pearson correlation between experimental panels of Nme2^Smu^-ABE8e and eNme2-C effectors for the guide-target library experiments, comparing the mean editing rates across the adenines of a library member target site. Each experimental panel consisted of 3 biological replicates (Rep1, 2 or 3) compiled from Figure 3c (panel 1)**, Supplementary** Figure 10 (panel 2), and **Supplementary** Figure 9 (panel 3). (**b**) Summary statistics for data in (**a**); CI indicates confidence interval, and *n* indicates number of comparisons.

**Supplementary Table 1.** Amino acid substitutions of Nme2Cas9 and Nme2^Smu^Cas9 arginine mutants.

| AA Mutation Location and Identity in nuclease | Codon Usage | Nucleic Acid Proximity in Homology Model | Domain | Note |
| --- | --- | --- | --- | --- |
| D56R | CGT | sgRNA | Nme2 - Bridge Helix |  |
| E186R | CGT | sgRNA | Nme2 - REC1 |  |
| D597R | CGT | sgRNA | Nme2 - HNH |  |
| D598R | CGT | sgRNA | Nme2 - HNH |  |
| D669R | CGT | sgRNA | Nme2 - RuvC III |  |
| E885R | CGT | sgRNA | Nme2 - WEDGE |  |
| E887R | CGT | sgRNA | Nme2 - WEDGE |  |
| E1079R | CGT | sgRNA | Nme2<sup>Smu</sup> - PID | Specific to Nme2<sup>Smu</sup> Effectors |
| T72Y | TAT | sgRNA | Nme2 - Bridge Helix |  |
| E50R | CGT | non-target DNA | Nme2 - RuvC I |  |
| D658R | CGT | non-target DNA | Nme2 - Linker II |  |
| E659R | CGT | non-target DNA | Nme2 - Linker II |  |
| D660R | CGT | non-target DNA | Nme2 - Linker II |  |
| E932R | CGT | non-target DNA | Nme2 - WEDGE |  |
| E989R | CGT | target & non-target DNA | Nme2<sup>Smu</sup> - WEDGE |  |
| D1048R | CGT | target & non-target DNA | Nme2<sup>Smu</sup> - PID | Specific to Nme2<sup>Smu</sup> Effectors |
| D418R | CGT | Target DNA | Nme2 - REC1 |  |
| D442R | CGT | Target DNA | Nme2 - REC1 |  |
| E443R | CGT | Target DNA | Nme2 - REC1 |  |
| D451R | CGT | Target DNA | Nme2 - REC1 |  |
| D470R | CGT | Target DNA | Nme2 - RuvC II |  |
| E471R | CGT | Target DNA | Nme2 - RuvC II |  |
| E508R | CGT | Target DNA | Nme2 - RuvC II |  |
| E520R | CGT | Target DNA | Nme2 - Linker I |  |
| E524R | CGT | Target DNA | Nme2 - Linker I |  |
| E552R | CGT | Target DNA | Nme2 - HNH |  |
| E585R | CGT | Target DNA | Nme2 - HNH |  |
| E587R | CGT | Target DNA | Nme2 - HNH |  |
| E844R | CGT | Target DNA | Nme2 - WEDGE |  |
| E868R | CGT | Target DNA | Nme2 - WEDGE |  |
| E873R | CGT | Target DNA | Nme2 - WEDGE |  |
| D947R | CGT | Target DNA | Nme2<sup>Smu</sup> - PID | Specific to Nme2<sup>Smu</sup> Effectors |
| A987R | CGT | Target DNA | Nme2<sup>Smu</sup> - PID | Equivalent to D987 in Nme2 |

**Supplementary Table 2.** Amino acid substitutions present within previously published PAM-relaxed Nme2Cas9 variants. Black-highlighted amino acid mutations are unique to specific Nme2Cas9 variants, while red-highlighted mutations indicate overlapping positions between previously characterized mutations and/or the Nme2Cas9 arginine variants described in this study.

| eNme2-C | eNme2-C_NR | iNme2 |
| --- | --- | --- |
| P6S |  | D844G |
| E33G |  | E868K |
| K104T | K104T | K870R |
| D152A | D152A | D873A |
| F260L | F260L | D911G |
| A263T | A263T | K929R |
| A303S | A303S | E932K |
| D451V | D451V |  |
| E520A |  |  |
| R646S |  |  |
| F696V |  |  |
| G711R |  |  |
| I758V |  |  |
| H767Y |  |  |
| E932K | E932K |  |
| N1031S | N1031S |  |
| R1033G | R1033G |  |
| K1044R | K1044R |  |
| Q1047R | Q1047R |  |
| V1056A | V1056A |  |

