## Supplementary material for "Rational Design of Enhanced Nme2Cas9 and Nme2^Smu^Cas9 Nucleases and Base Editors": Inventory of Supplementary Items

Supplementary Data 1. Dataset for nuclease panel1 of the activity guide-target.  
Supplementary Data 2.  Dataset for nuclease panel2 of the activity guide-target.
Supplementary Data 3.  Dataset for ABE panel1 of the activity guide-target.
Supplementary Data 4. Dataset for ABE panel2 of the activity guide-target.
Supplementary Data 5.  Dataset for ABE panel3 of the activity guide-target.
Supplementary Data 6. Dataset for ABE panel4 of the activity guide-target.
Supplementary Data 7. Dataset for the nuclease mismatch library.
Supplementary Data 8. Dataset for the ABE mismatch library.
Supplementary Data 9. Oligonucleotides and target sites used within the study.
