## Supplementary Code for "Rational Design of Enhanced Nme2Cas9 and Nme2^Smu^Cas9 Nucleases and Base Editors": demuxMM.py

```
#!/usr/bin/env python
# coding: utf-8

## Run first - Import all required programs
import sys
from collections import defaultdict
import gzip
from Bio.Align import PairwiseAligner
import numpy as np
import pandas as pd
import Bio.Seq
import Bio.SeqIO
from Bio.SeqRecord import SeqRecord
import os
from thefuzz import fuzz
from thefuzz import process
from Levenshtein import distance
from Bio import pairwise2
from Bio.Seq import Seq


# Load in inputs
# Reference .csv File with library elements
input_df = pd.read_csv('/home/nathan.bamidele-umw/libm_demux_test/demux_Ref.csv')

# Get the folder name from input filenames
folder_name = os.path.splitext(os.path.basename(sys.argv[1]))[0].rsplit('_', 3)[0]

spacer_sequences = np.array(list(input_df['sp']))
barcode_sequences_OG = np.array(input_df['bc'])
barcode_sequences = np.array(input_df['bc_rc'])
target_sequences = np.array(list(input_df['tgt']))

spacer_list = input_df['sp'].tolist()
barcode_list = input_df['bc_rc'].tolist()
target_list = input_df['tgt'].tolist()

homology_5p = 'GTGGAAAGGACGAAACACCG'
scaff_seq1 = 'gttgtagctccctttctcatttcgg'
scaff_seq_2 = 'AACGATGCCCC'
homology_3p = 'TCTAGAGGTACGGCCGCTTCGA'


# To get my sequencing files loaded in
def readFastq(filename):
    """Reads gzip-compressed FASTQ file."""
    sequences = []
    qualities = []
    with gzip.open(filename, 'rt') as fh:  # 'rt' mode reads the file as text
        while True:
            fh.readline()  # skip name line
            seq = fh.readline().rstrip()   # read base sequence
            fh.readline()                  # skip placeholder line
            qual = fh.readline().rstrip()  # read base quality line
            if len(seq) == 0:
                break
            sequences.append(seq)
            qualities.append(qual)
    return sequences, qualities


# Get the sequences loaded in
seqsR1, qualsR1 = readFastq(sys.argv[1])  # First argument is R1 fastq.gz filename
seqsR2, qualsR2 = readFastq(sys.argv[2])  # Second argument is R2 fastq.gz filename


# Make the folder you need
os.makedirs(folder_name, exist_ok=True)

# Programs for quality and to write the fastq
def quality_checker(qual):
    arr = bytes(qual, 'utf-8')
    """Checks quality score of the reads"""
    qual1 = (np.frombuffer(arr, dtype=np.uint8) - 33).mean()
    return qual1


def to_IOSeq_rec(sensor_seq, i, q1):
    qname = 'read_' + str(i)
    record = SeqRecord(Bio.Seq.Seq(sensor_seq), id=qname, name=qname, description='', dbxrefs=[])
    arr = bytes(q1, 'utf-8')
    qual1 = (np.frombuffer(arr, dtype=np.uint8) - 33)
    record.letter_annotations["phred_quality"] = qual1[0:len(sensor_seq)]
    return record


# All the programs for finding the barcode, the spacer and if there are multiple barcodes
def barcode_identifier(r2, barcode_sequences, scaff_seq_2):
    output1 = None
    score2 = None
    bc_end = None
    bc_start = None
    barcode_seq = None
    bc_idx = None
    bc_idx_1 = None
    Barcode_Match = None
    quality = None
    alignments_1 = pairwise2.align.localms(r2, "CAAAAAATAAACGATGCCCCTTAAA", 2, -1, -2, -2, one_alignment_only=True)
    best_alignment_1 = alignments_1[0]
    bc_end = best_alignment_1[3]
    bc_start = best_alignment_1[3] - 13
    barcode_seq = r2[bc_start:bc_end]
    output1, score1 = process.extractOne(barcode_seq, barcode_list)
    if score1 >= 88:
        bc_idx = np.where(barcode_sequences == output1)[0]
        if len(bc_idx) == 1:
            Barcode_Match = 'Yes'
            bc_idx_1 = int(str(bc_idx)[1:-1])
        if len(bc_idx) > 1:
            Barcode_Match = 'Multiple'
            bc_idx_1 = bc_idx
    else:
        Barcode_Match = 'No'
        bc_idx_1 = 'No_Barcode_Match'
    return bc_idx_1, Barcode_Match


def spacer_matcher(r1, bc_idx_1, Barcode_Match):
    index = None
    if Barcode_Match == 'Yes':
        alignments_2 = pairwise2.align.localms(r1, 'GTTGTAGCTCCCTTTCTCATTTCGG',  2, -1, -2, -2, one_alignment_only=True)
        best_alignment_2 = alignments_2[0]
        sp_start = best_alignment_2[3] - 23
        sp_end = best_alignment_2[3]
        sp_seq = r1[sp_start:sp_end]
        spacer = spacer_sequences[bc_idx_1]
        Ratio1 = fuzz.ratio(spacer, sp_seq)
        if Ratio1 >= 92:
            out = 'Match_Sp_Bc'
            index = bc_idx_1
        else:
            out = 'Recombined_Sp_Bc'
            index = 'None'
    else:
        out = 'No_Barcode_Match'
    return index, out


def multiple_barcode(r1, bc_idx_1, Barcode_Match):
    target_idx = None
    if Barcode_Match == 'Multiple':
        for i in range(0, len(bc_idx_1)):
            bc_idx_i = bc_idx_1[i]
            alignments_2 = pairwise2.align.localms(r1, 'GTTGTAGCTCCCTTTCTCATTTCGG',  2, -1, -2, -2, one_alignment_only=True)
            best_alignment_2 = alignments_2[0]
            sp_start = best_alignment_2[3] - 23
            sp_end = best_alignment_2[3]
            sp_seq = r1[sp_start:sp_end]
            spacer = spacer_sequences[bc_idx_i]
            Ratio1 = fuzz.ratio(spacer, sp_seq)
            if Ratio1 >= 92:
                out = 'Match_Sp_Bc'
                index = bc_idx_i
                break
            else:
                out = 'Recombined_Sp_Bc'
                index = 'None'
                continue
    return out, index


def run_program(r1, r2, quallity1, qual2, barcode_sequences, spacer_sequences):
    cols = ['good_quality', 'bad_quality', 'useable', 'recombined', 'No_Barcode']
    z = np.zeros((len(cols), 1))
    d2 = pd.DataFrame(dict(zip(cols, z)))
    for i in range(1, len(seqsR1)):
        q1 = quality_checker(qualsR1[i])
        q2 = quality_checker(qualsR2[i])
        if q1 >= 26 and q2 >= 26:
            d2.loc[0, 'good_quality'] += 1
            A, B = barcode_identifier(r2[i], barcode_sequences, scaff_seq_2)
            if B == 'Yes':
                C, D = spacer_matcher(r1[i], A, B)
                if D == 'Match_Sp_Bc':
                    d2.loc[0, 'useable'] += 1
                    out_file = folder_name + '/' + 'M' + str(A) + '.fastq'
                    record = to_IOSeq_rec(r2[i], i, qualsR2[i])
                    with open(out_file, 'a') as fq:
                        Bio.SeqIO.write(record, fq, 'fastq')
                if D == 'Recombined_Sp_Bc':
                    d2.loc[0, 'recombined'] += 1
            if B == 'No':
                d2.loc[0, 'No_Barcode'] += 1
            if B == 'Multiple':
                E, F = multiple_barcode(r1[i], A, B)
                if E == 'Match_Sp_Bc':
                    d2.loc[0, 'useable'] += 1
                    out_file = folder_name + '/' + 'M' + str(F) + '.fastq'
                    record = to_IOSeq_rec(r2[i], i, qualsR2[i])
                    with open(out_file, 'a') as fq:
                        Bio.SeqIO.write(record, fq, 'fastq')
                if E == 'Recombined_Sp_Bc':
                    d2.loc[0, 'recombined'] += 1
        else:
            d2.loc[0, 'bad_quality'] += 1
    return d2


# Run this for the big one
d2 = run_program(seqsR1, seqsR2, qualsR1, qualsR2, barcode_sequences, spacer_sequences)

# This will tell you where what your output is
print(d2)
```
